# Protein-ligand binding kinetics are primarily controlled by the protein, not the ligand

**DOI:** 10.64898/2026.05.20.726507

**Authors:** Bharath Srinivasan, Ana Corrionero, Marco Barone, Patricia Alfonso, Niall Prendiville, Tatiana Cazorla, Anza Suneer Rahiyanath, Adrian Whitty, Peter Tonge

**Author notes:** These authors contributed equally (Bharath Srinivasan, Ana Corrionero).

## Abstract

Protein–ligand interactions underpin biological regulation and drug action, with both binding affinity and binding kinetics shaping functional outcomes. By analysing kinetic data for 4,311 protein-small-molecule pairs, we find that when association occurs below the diffusion-controlled limit, the rates of ligand association (*k*_on_) and dissociation (*k*_off_) are primarily determined by how the initial encounter complex reorganizes into the final bound state, and that this reorganization is governed chiefly by the intrinsic dynamic properties of the protein rather than by structural features of the ligand. Counterintuitively, therefore, *k*_off_ exhibits minimal dependence on ligand structure, so that dissociation proceeds through protein-gated conformational transitions rather than through direct rupture of protein–ligand contacts. This mechanistic behaviour stands in marked contrast to that for protein–protein complexes, based on an analysis of 1,561 interactions. Together, these findings challenge prevailing assumptions regarding the molecular determinants of small-molecule binding kinetics, and have broad implications for rationally modulating protein–ligand interactions and drug-target residence times.

## INTRODUCTION

It is now widely recognized that considering only equilibrium binding affinity (*K*_d_) or half-maximal inhibitory concentration (IC_50_) provides an incomplete picture of the biochemical basis for the pharmacological potency of drugs, and that the temporal dynamics of drug-target engagement can critically influence pharmacodynamics, duration of efficacy, and off-target effects. Consequently, direct measurement of the rates of drug-target binding (*k*_on_) and dissociation (*k*_off_) is increasingly recognized as essential for rational lead optimization, leading to a corresponding increase in the availability of high quality kinetic data for protein-ligand binding interactions.^1-12^

The initial hit compound that serves as the starting point for many drug discovery efforts is typically only a weak binder (*K*_d_ = 10^-3^-10^-7^ M), and therefore one of many aspects of its advancement to a drug is to increase its potency. This is most often achieved by progressively modifying the structure of the compound to increase its binding affinity for the target, and the relative importance of changes in *k*_on_ versus *k*_off_ in these affinity increases have been the subject of considerable discussion.^4,6-9,11,13-15^

In this study, we report kinetic data for 606 protein-kinase complexes showing that, contrary to expectations, for interactions weaker than *K*_d_ ∼10^-8^ M differences in binding affinity result primarily from changes in the association rate constant, *k*_on_, and not in *k*_off_. The dissociation rate constant becomes the driving factor only for high affinity interactions where *k*_on_ is diffusion controlled. We show that this observation holds true for 3,238 additional kinase-ligand pairs from the KIND database of protein-ligand binding kinetics, as well as for other target classes such as some GPCRs, HSP90, and the hERG ion channels.^14,16^

The results can be explained, in part, by considering that ligand binding is a multi-step process in which the initial encounter complex can partition forwards to give the final inhibited complex or backwards to reform free ligand and protein. However, applying this model reveals the surprising result that the activation barrier for the initial step in the dissociation of the final complex is largely insensitive to the structure of the ligand. This result implies that protein-ligand contacts remain fully or mostly intact in the transition state for the initial step in ligand dissociation. Reaching the first transition state in ligand dissociation therefore involves a ligand-independent process–presumably an energetically costly rearrangement of the protein– that precedes any significant displacement of the ligand from its bound location. While protein-gated ligand dissociation has been postulated previously for a small number of specific systems,^17-19^ our results indicate that such protein-gated dissociation may be a general feature of protein-small ligand binding. These small ligands therefore bind via a mechanism akin to that seen for slow-binding inhibitors, and quite distinct from the binding mechanism of protein-protein interactions.

## RESULTS & DISCUSSION

### Affinity-kinetics relationship for inhibitors of Bruton’s tyrosine kinase (BTK)

To explore how incremental improvements in ligand binding affinity affect the rate constants for ligand binding, *k*_on_, and dissociation, *k*_off_, we revisited a kinetic data set from the design of novel Bruton’s tyrosine kinase (BTK) inhibitors in which the on- and off-rates were measured using a time-resolved Förster Resonance Energy Transfer (TR-FRET) assay in the KINETICfinder platform, with the dissociation constant, *K*_d_, for each compound being calculated from the ratio *k*_off_/*k*_on_.^20^ **Figure 1 A-B** shows the *k*_on_ and *k*_off_ values for 38 small-molecule inhibitors binding to BTK plotted as a function of their *K*_d_ values, measured separately for the unphosphorylated and phosphorylated forms of the enzyme at 25°C (**Table S1**).

**Figure 1.**
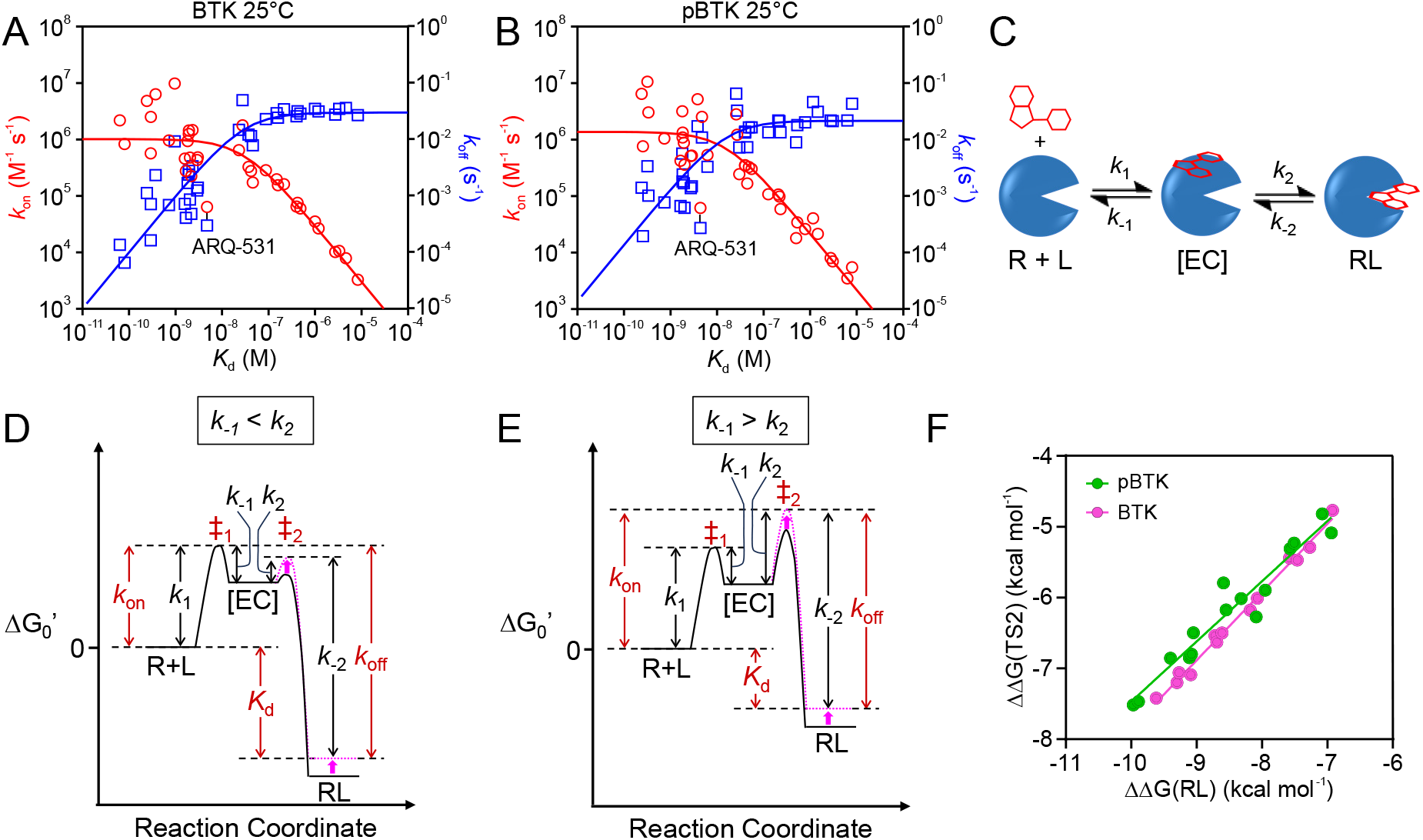
Plots of *k*_on_ (red circles) and *k*_off_ (blue squares) against *K*_d_ for 38 small molecules binding to (**A**) unphosphorylated BTK (BTK) and (**B**) phosphorylated BTK (pBTK), determined using KINETICfinder TR-FRET binding assays. The data are fitted to *Equation 1* for *k*_on_ and *Equation 2* for *k*_off_. (**C**) Two-step binding mechanism for a ligand, L (red) with a receptor, R (blue), where the initial diffusional encounter with rate constant *k*_1_ involves a ‘sticky’ collision giving an initial encounter complex, [EC], that can partition backwards to regenerate free R and L, or forwards to give the final complex, RL. (**D**) Free energy profile for the two-step binding mechanism from (C) in which the first step is rate limiting, causing EC to partition mostly forward, as might be expected for the highest affinity ligands. A small change in binding affinity (magenta arrow) will change *k*_off_ but not *k*_on_ provided *k*_1_ remains constant. (**E**) Free energy profile for the alternative case in which the second step is rate limiting, showing that a small change in affinity will change *k*_on_ but not *k*_off_ provided *k*_-2_ remains constant. For (D) and (E), the rate constant(s) controlled by each energy barrier are indicated, with a higher activation barrier corresponding to a lower value for the rate constant. (**F**) Linear Free Energy Relationship showing how the free energy of the transition state for the second step, ‡_2_, varies with that of the RL complex for the lower affinity inhibitors from panel (A) (BTK, pink) and (B) (pBTK, green). The solid lines represent linear regression fits to the data, with slopes of ϕ = 0.96 ± 0.03 and 0.85 ± 0.06, respectively.

**Figure 1A** shows that, for weak inhibitors of unphosphorylated BTK (*K*_d_ ≥ 10 nM), *k*_on_ increases in proportion to 1/*K*_d_ and then plateaus at 10^6^-10^7^ M^-1^s^-1^ for higher affinity binders. Because *K*_d_ = *k*_off_/*k*_on_, the data for *k*_off_ necessarily show the complementary trend in which, for the stronger binders, *k*_off_ increases proportionately with *K*_d_, with a break at *K*_d_ ∼ 10 nM above which the *k*_off_ values appeared invariant at ∼0.03 s^-1^ (t_1/2_ ∼ 20 s). Similar behavior was seen for BTK that has been phosphorylated (pBTK) (**Figure 1B**), and for unphosphorylated and phosphorylated BTK at 37°C (**Figure S1**). That protein-ligand association rate constants (*k*_on_) generally do not exceed ∼10^6^-10^7^ M^-1^s^-1^ is known, and is attributable to binding having reached the effective diffusion-controlled encounter limit.^21-23^ However, the observation that below a certain affinity threshold *k*_off_ becomes invariant is unexpected, with no comparable physical limit to explain this asymptote.

These results can be understood, in part, by recognizing that binding of a ligand to a specific site on a protein is generally not a one-step process. Instead, the initial collisional encounter is ‘sticky’, resulting in a metastable encounter complex that can partition forward to give the final inhibited complex or backward to give the free ligand and target (**Figure 1C**). The existence of a transient encounter complex as an intermediate on the binding path is well-accepted for protein-protein binding,^24^ but is less widely discussed in the literature on protein-small molecule complexes.^23,25-27^ The fraction, *F*, of encounter complexes that partition forward to give the final complex is governed by the relative heights of the energy barriers for *k*_2_ versus *k*_-1_, as defined in **Figure 1C**, according to the expression *F* = *k*_2_/(*k*_-1_ + *k*_2_). The data in **Figures 1A** and **B** indicate that, at *K*_d_ ∼ 10^-8^ M, there is a change in rate limiting step such that, for the stronger-binding ligands, binding is fast with the encounter complex partitioning mostly forward such that *k*_on_ ∼ *k*_1_ (**Figure 1D**), while for weaker-binding ligands the *k*_2_ step has become rate-limiting so that differences in *K*_d_ now result primarily from differences in *k*_on_ (**Figure 1E**). This change in rate-limiting step can be explained if the incremental destabilization of the bound state for progressively weaker binders (i.e. increasing values of *K*_d_) also destabilizes the transition state for *k*_2_ such that, at some affinity threshold, *k*_2_ becomes the rate-limiting step for binding. This behavior is described by *Equations 1*-*3*. The curve fits in **Figures 1A** and **1B** were derived from this model, showing that it can account for the observed trends in *k*_on_ and *k*_off_ with *K*_d_, albeit with some scatter around the trendlines from one BTK-inhibitor complex to another, especially among the high affinity binders.

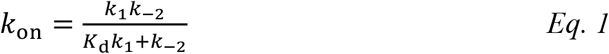

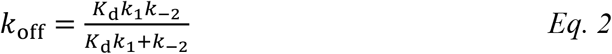

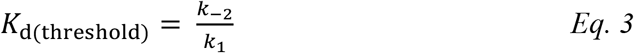

An unexpected feature of the equilibrium-kinetic trends shown in **Figures 1A** and **B** is that, for the inhibitors with *K*_d_ ≥ 10^-8^ M, *k*_off_ has a constant value. According to the above model, in this affinity regime *k*_off_ corresponds to *k*_-2_ (**Figure 1E**). The observed invariance of *k*_off_ with *K*_d_ in this regime therefore implies that the value of *k*_-2_, which reflects the relative energies of the bound complex versus the transition state for the *k*_-2_ step, is essentially insensitive to the structure of the ligand across a set of complexes that span ∼3 logs of affinity. The extent to which changes in ligand structure affect the free energy of the transition state for *k*_-2_ versus that of the bound complex can be quantified by constructing a linear free energy relationship between these two quantities.^28^ Changes in the energy of the bound receptor-ligand complex, relative to the reference state of free protein plus free ligand, are simply given by ΔΔG^‡^ = -RTln(1/*K*_d_). The corresponding changes in the free energy of the transition state for *k*_-2_ are given by the Eyring-Polanyi equation as ΔΔG^‡^ = -RTln(*k*_off_/*K*_d_) = -RTln(*k*_on_). Such a plot can have a slope ranging from zero, indicating the free energy of transition state 2 (‡_2_) is insensitive to changes in the energy of the RL complex, to a value of 1, indicating that any change in the free energy of RL is exactly mirrored in ‡_2_. For the data from **Figure 1** the relationship has a slope of ϕ = 0.96 ± 0.03 for unphosphorylated BTK, and ϕ = 0.85 ± 0.06 for the phosphorylated enzyme (**Figure 1F**), indicating that the effects of ligand structure on the stability of the bound complex are almost quantitatively mirrored in the transition state for the initial step in ligand dissociation. This observation is difficult to account for in terms of our conventional understanding of the process of ligand dissociation, in which the transition state is presumed to involve disengagement of the ligand from its binding site with concomitant loss of protein-ligand binding energy.^27,29-33^ Instead, our analysis implies that, at least for the BTK ligands with *K*_d_ ≥ 10^-8^ M, the binding contacts between protein and ligand are essentially fully retained in the transition state for dissociation.

Application of the two-step binding model to the affinity-kinetics relationships observed in **Figure 1A** and **B** thus poses a number of intriguing questions: (1) What is the nature of the structural differences between the RL complex and the transition state for dissociation that account for the latter being the highest energy state of the system, if it is not loss of binding energy with the ligand? (2) What determines the ligand affinity threshold at which affinity optimization ceases to increase *k*_on_ and instead begins to decrease *k*_off_? (3) Why is there more scatter in the data on the high affinity side of this affinity threshold than to the low affinity side? (4) What factors cause the occasional protein-ligand pair to show as an outlier in the analysis? (5) Is the observed affinity-kinetic relationship particular to BTK, or perhaps to kinases in general, or do other protein-ligand classes behave similarly?

### Extension to additional kinase-inhibitor complexes

To provide insight into the questions posed above, and to understand the generality of the affinity-kinetic trends observed for the BTK inhibitors, we first extended the analysis to kinetic data for a set of 455 kinase-inhibitor binding interactions measured using the same KINETICfinder TR-FRET binding assays used in **Figure 1** (**Table S1** and **S2**) that broadly span the phylogenetic distribution of the human kinome (**Figure S2**). Rather than plotting rate constants against *K*_d_ for multiple inhibitors with a particular enzyme, we instead compared data for 7 individual inhibitors, danusertib, dasatinib, encorafenib, foretinib, tivozanib, tucatinib, and zimlovisertib binding to a range of kinase targets numbering anywhere between 19 (for tucatinib) to 108 (for danusertib) (**Tables S3-S9**).

**Figure 2A** shows the results plotted for all 455 kinase-ligand pairs, demonstrating that the trends identified in **Figure 1** hold true for this broad set of kinase targets and compounds. Analysis of subsets of the data for individual highly represented inhibitors also shows behaviors similar to those seen above for inhibitors of BTK. In particular, for the broad spectrum inhibitors dasatinib and danusertib, which bind to the ‘DFG-in’ conformation of the kinase (‘Type I’ inhibitors), the plots clearly show that, for the weaker interactions, differences in affinity are primarily controlled by variation in *k*_on_, while for the higher affinity interactions (*K*_d_ ≤ ∼10^-8^ M) affinity differences are driven primarily by differences in *k*_off_ (**Figure 2B, C**). The broad-spectrum inhibitors foretinib and tivozanib, which are Type II inhibitors that bind to the ‘DFG-out’ kinase conformation, show similar trends for the weaker complexes but with greater variation in *k*_on_ for the high affinity binders (**Figure 2D, E**). The highly selective kinase inhibitors encorafenib, tucatinib, and zimlovisertib, all of Type I, by definition have few or no high affinity complexes beyond those with their specific targets, but the weaker complexes again show relatively invariant *k*_off_ values, with *K*_d_ primarily depending on *k*_on_ (**Figure 2F-H**). These results show that the affinity-kinetics relationship described in **Figure 1** is not unique to BTK as a kinase target.

**Figure 2.**
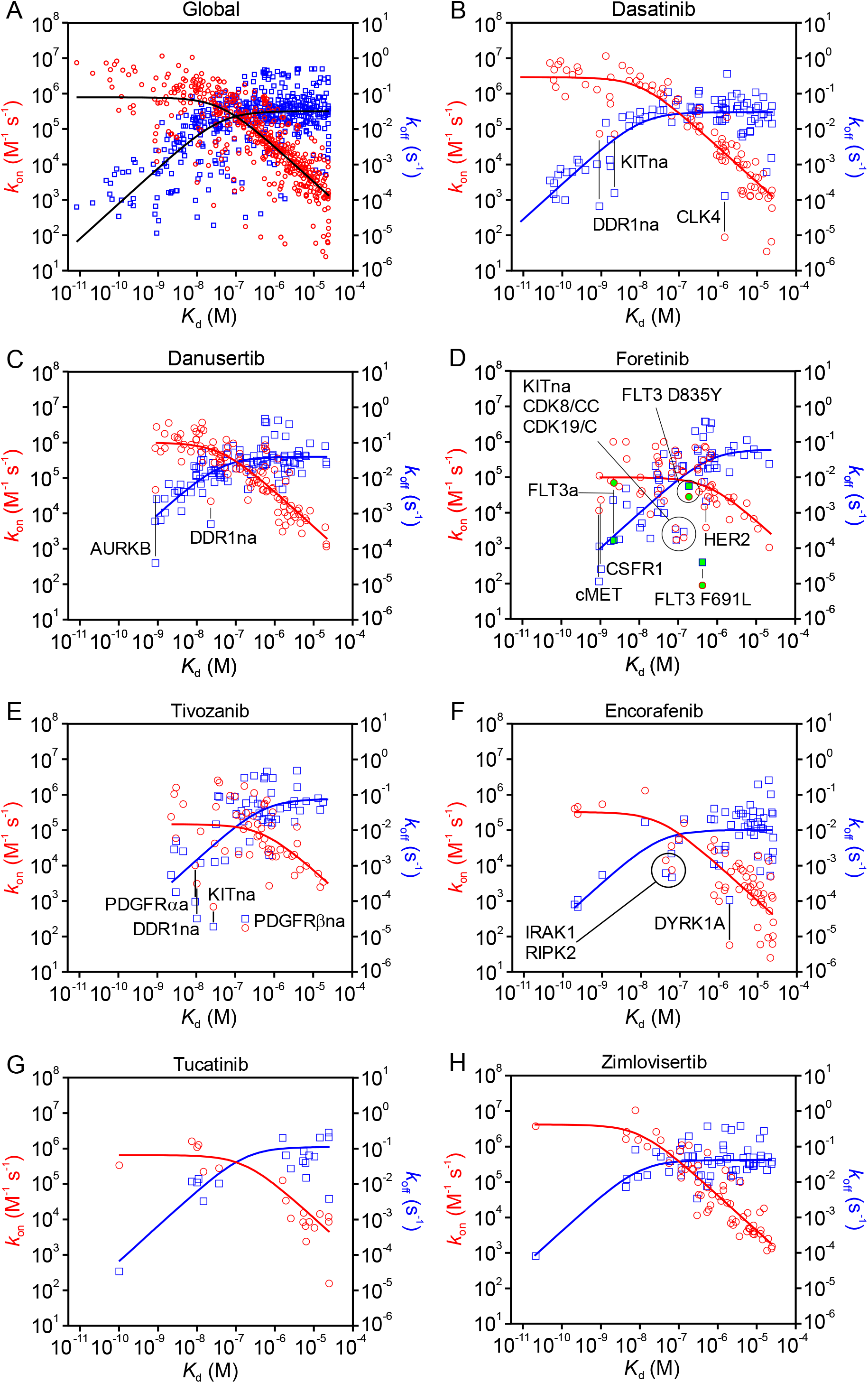
Plots of *k*_on_ (red circles) and *k*_off_ (blue squares) against *K*_d_ for various kinase-inhibitor combinations. (**A**) Data for 455 kinase-inhibitor combinations measured using the same KINETICfinder TR-FRET binding assay used in **Figures 1A** and **B**. Data for the individual inhibitors are shown in (**B**) dasatinib (89 kinases), (**C**) danusertib (108 kinases), (**D**) foretinib (64 kinases), (**E**) tivozanib (56 kinases), (**F**) encorafenib (55 kinases), (**G**) tucatinib (19 kinases), and (**H**) zimlovisertib (64 kinases). The data are fitted to *Equation 1* for *k*_on_ and *Equation 2* for *k*_off_ (solid lines). In panel **A** the fitted lines are shown in black for clarity. Selected outlier kinases are labeled on the plots. The suffixes ‘na’ and ‘a’ indicate the non-activated and activated forms of the kinase, respectively (**Table S2**); the absence of a suffix indicates that the activation status is unknown. Data for FLT3 wild-type and two mutants, D835Y and F691Y, are colored green.

### Extension to other target classes and assay methods

To assess the broader generality of the behavior seen in **Figures 1** and **2** we expanded the study to include data from the KIND database,^6,16^ which contains kinetic measurements for the interactions of 3,812 compounds with 78 different targets and has no overlap with the data plotted in **Figures 1** and **2**. The majority of the data (3,238 target-ligand pairs) are for small molecule inhibitors of different kinases. The other target classes most highly represented in this database include GPCRs (242 small molecule ligands), heat shock protein-90 (HSP90, 160 inhibitors), and the human ether-a-go-go-related (hERG) potassium voltage-gated ion channel (45 inhibitors). The methods used to measure the kinetic parameters encompass several different biochemical and cellular techniques such as radioligand binding assays, surface plasmon resonance, fluorescent ligand binding assays, and xCELLidence.^6,14,16^

Analysis of the kinase inhibitor subset from KIND (**Figure 3A**) showed a similar affinity-kinetics trend to those seen in **Figures 1** and **2**. Across this diverse set of kinases and ligands the limiting values for *k*_on_ seen at high affinities and for *k*_off_ seen at low affinities vary over 2-3 logs, as was seen with the kinase dataset in **Figure 2A** determined using KINETICfinder TR-FRET binding assays. Broadly similar affinity-kinetics trends are seen for the KIND data subset involving GPCRs (**Figure 3B**), though with greater scatter around the trend. However, examining the different GPCRs represented in **Figure 3B** individually (**Figure S3**) showed that some, such as the dopamine receptor D2 (**Figure 3C**), displayed a clear affinity-kinetics relationship qualitatively similar to that seen for the kinases. For other GPCRs, such as the metabotropic glutamate receptor, mGlu (**Figure 3D**), the behaviour was consistent with the same model, but the available data spanned a small affinity range that encompassed only *k*_on_-controlled or *k*_off_-controlled behaviour. The data for HSP90 (**Figure 3E**) and hERG (**Figure 3F**) are similarly consistent with the binding model described above. Although there is insufficient data to fully define both limbs of the affinity-kinetics relationship, it can clearly be seen that the affinity of weak binding inhibitors is controlled by *k*_on_ rather than *k*_off_.

**Figure 3.**
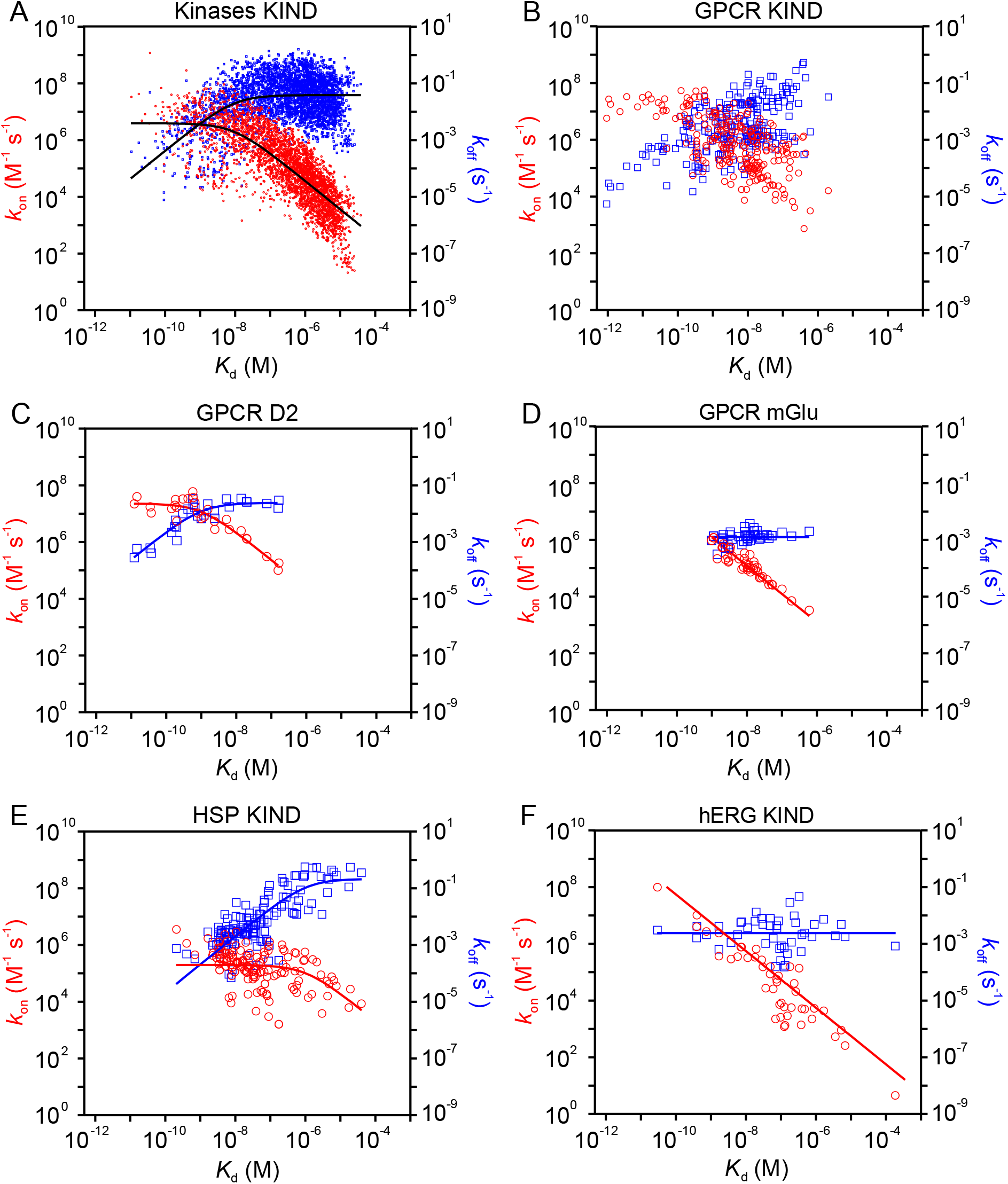
Plots of *k*_on_ (red) and *k*_off_ (blue) versus *K*_d_ for small molecules binding to four distinct target classes, using data from the KIND database: (**A**) 3,238 kinase-inhibitor complexes, (**B**) 242 G-protein coupled receptors (GPCR)-ligand complexes, including (**C**) the dopamine receptor, D2, and (**D**) the metabotropic glutamate receptor, mGlu; (**E**) 160 heat shock protein-90 (HSP90) complexes, and (**F**) 45 hERG voltage-gated ion channel complexes. The solid lines are fits to Equations 1 (for *k*_on_ values) or 2 (for *k*_off_), or to the linear equations these simplify to at high values of *K*_d_. In panel A the fitted lines are shown in black for clarity.

These results show that the observation that *K*_d_ is mainly controlled by *k*_on_ for low affinity interactions is independent of assay method and applies across several unrelated target types. The limiting values observed for *k*_on_ and *k*_off_ differ from one target type to another, however, as does the affinity threshold that differentiates the two kinetic regimes (**Table S10**). In particular, compared to the kinase-inhibitor complexes, which tend to show limiting *k*_off_ values in the range of 10^-1^-10^-2^ s^-1^, the HSP90 inhibitors tend to dissociate somewhat faster while the hERG ligands dissociate more slowly. The GPCRs tend to bind their ligands 1-2 orders of magnitude faster than is seen for the kinases and have *k*_off_ values that span a wide range extending both above and below the limiting value typical for kinases.

### Causes of outliers from the observed affinity-kinetics trends

#### High affinity binders

Several of the datasets in **Figures 1**-**3** show substantially greater scatter for the high affinity interactions, where *k*_on_ ∼ *k*_1_, than for the weaker binders where *k*_on_ is rate-limited by *k*_2_. For example, **Figures 1A** and **B**, where the target is invariant and only the ligand differs, show that *k*_1_ is quite sensitive to differences in ligand structure while *k*_-2_ is remarkably insensitive. These findings imply that *k*_1_ is generally more susceptible to variation than *k*_-2_, at least for different ligands binding to the same kinase protein. The diffusional encounter rate is sensitive to a number of factors, including the size and conformation of the unbound ligand, and long-range electrostatic interactions between ligand and receptor.^24^ Insight into how small differences in protein structure contribute to variations in these two rate constants can be gained from **Figure 2B-H**, in which both *k*_1_ and *k*_-2_ show modest sensitivity to the identity of the kinase.

#### Slow-binding inhibitors

Due to the emerging appreciation of drug-target residence time as an important factor in pharmacological activity,^8,34^ particular attention has been paid in recent years to slow-binding inhibitors. In most cases, slow-binding occurs via a multi-step mechanism in which the initial bound complex slowly converts to a thermodynamically more stable final bound state over a time frame of minutes or hours (**Figure 4A, B**).^35-37^ This final step in binding typically involves a slow conformational change that results in increased protein-ligand binding energy.^38-41^ Because the rate-limiting step for ligand binding and dissociation is the final step (‡_3_ in **Figure 4B**), for slow-binding inhibitors *k*_on_ << *k*_1_, and *k*_off_ = *k*_-3_ regardless of affinity.

**Figure 4.**
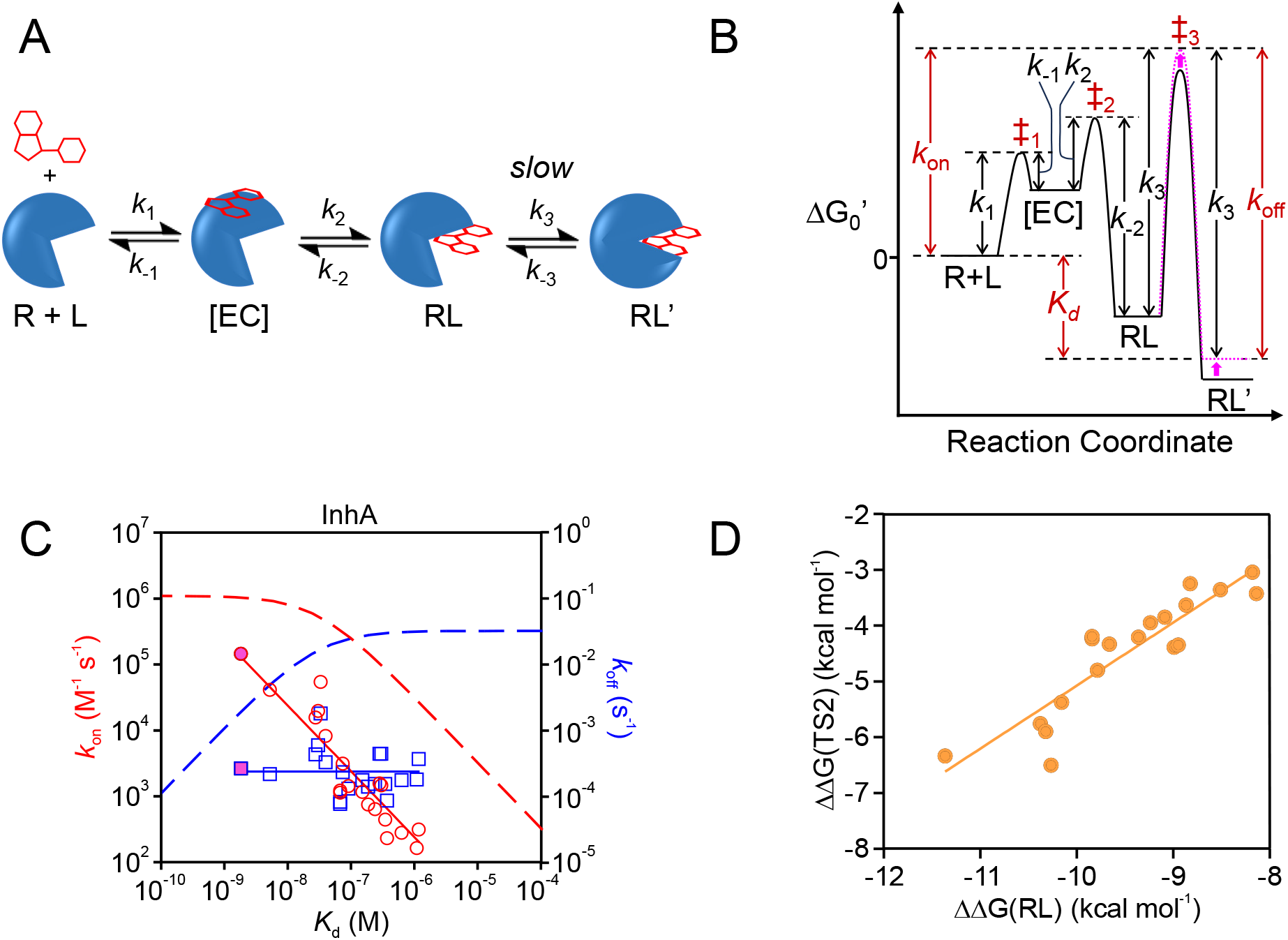
Affinity-kinetics relationship for slow-binding inhibitors. Mechanism (**A**) and free energy profile (**B**) for a slow-binding inhibitor. The rate constant(s) controlled by each energy barrier are indicated, with a higher activation barrier corresponding to a lower value for the rate constant. A small change in binding affinity (*magenta* arrow) will change *k*_on_ but not *k*_off_ provided *k*_*-3*_ remains constant. (**C**) Plot of *k*_*on*_ (red circles) and *k*_*off*_ (blue squares) versus *K*_d_ for a set of slow-binding inhibitors of *M. tuberculosis* InhA. The solid lines represent the best fits of the data to *k*_*off*_ = *k*_*-3*_ (blue line), and *k*_*on*_ = *k*_*-3*_/*K*_d_ (red line). The magenta data points are for binding of the isoniazid-NAD adduct. The dashed lines represent the corresponding relationships observed for the kinase BTK in **Figure 1A**, included as a reference for the behavior of conventional inhibitors. (**D**) Free energy profile showing how the free energy of the transition state for step 3, ‡_3_, compares to that of the RL’ complex for the InhA inhibitors from (**C**), showing a slope of ϕ = 1.13*±* 0.13.

**Figure 4C** shows kinetic data for a set of inhibitors for the *M. tuberculosis* enoyl-acyl carrier protein reductase, InhA, that have been shown to act as slow-binding inhibitors and include the INH-NAD adduct which is the active form of the tuberculosis drug isoniazid.^39,42-45^ The data show that, as was the case for the mostly conventional inhibitors described above, *k*_off_ for these compounds is roughly invariant with *K*_d_, implying that *k*_-3_ has a value of ∼10^-4^ M^-1^s^-1^, approximately 2 logs slower than for the previously discussed interactions, notably for kinases. Nonetheless, as was the case for the other target-ligand systems, replotting the data as an LFER for *k*_off_ versus *K*_d_ shows that ϕ ∼ 1.0 (**Figure 4D**), indicating that differences in the stability of the final inhibited complex, RL’, among the different compounds are closely mirrored in the energy of the transition state for the rate-limiting step, ‡_3_.

#### Additional outliers with low k_on_ and k_off_ values

The plots in **Figures 1-3** contain a number of outliers with low *k*_on_ and *k*_off_ values. These low outliers may be slow-binding inhibitors for which binding involves a rate-limiting third step, as discussed above for InhA. Indeed, several of these low outliers have previously been shown to display a slow-binding inhibition mechanism, such as ARQ-531 (**Figure 1**),^46^ and others may be previously unrecognized examples of slow-binding inhibition. Alternatively, some low outliers could instead be compounds that bind through the two-step mechanism described in **Figure 1** but which, for unknown reasons, have a suppressed value for *k*_-2_. Several of the outliers in **Figure 2** are non-activated kinases, and it is possible that in these cases reorganization of the encounter complex to give the final inhibitor complex is slowed by the need to displace the unphosphorylated activation loop from a position that occludes the inhibitor’s binding site.^47^

### Relation to protein-protein and protein-peptide binding

To better understand the kinetic trends observed for protein–small ligand interactions, we performed a comparative analysis of protein–protein and protein–peptide interactions, including some antibody-antigen interactions. The results show that the behavior for the protein-small molecule complexes that we presented in the previous sections differs from that observed for protein-protein complexes. Examination of protein-protein interaction kinetic data from the SKEMPI-2 database which includes 1,561 entries encompassing mutational variations of 66 distinct pairs of interaction partners (**Figure 5A, Figure S4**),^48^ shows that differences in binding affinity achieved by mutating interacting proteins was most commonly driven by changes in *k*_off_. The binding of *B. anthracis* β-lactamase to BLIP-II provides one of many examples (**Figure 5B**). Similar behavior was seen for peptides binding to the T-cell receptor, as illustrated by the data for HLA-A2/Tax peptide complex binding to the A6 T-cell receptor in **Figure 5C**. As described above, this finding that changes in *K*_d_ are due to differences in *k*_off_ is expected if binding is diffusion controlled, or if for other reasons mutational effects on the stability of the bound complex do not affect the energy of the transition state for dissociation. In some cases, changes in *K*_d_ were accompanied by small or moderate changes in *k*_on_. Thus, while most protein-protein interactions in the SKEMPI2 database showed linear free energy relationships between the free energy of the bound complex and the transition state for dissociation with ϕ ∼ 0, a continuum of values ranging up to ϕ ∼ 0.5 was seen. However, only rarely was *k*_on_ a dominant driver of affinity differences (ϕ > 0.5). Closer examination shows that, in these cases, the changes in *k*_on_ correlated with mutational changes in the net charges on the interacting proteins (**Figure 5D**), suggesting that binding is diffusion-controlled and that the effects on *k*_on_ result from changes in the diffusional encounter rate due to electrostatic funneling or perhaps to changes in the number of strongly solvated groups affecting the protein’s hydrodynamic radius. For example, examination of *k*_on_ versus *K*_d_ for the interaction of the TEM1 β-lactamase to BLIP shows that the mutants fall into two classes of behavior, one involving changes in the net charge on the protein for which effects on *K*_d_ depend primarily on differences in *k*_on_, and another subset the change in charge introduced by the mutations is zero or small for which *k*_on_ is invariant and affinity differences are driven by *k*_off_.^24^ Thus, the protein-protein or protein-peptide interactions display quite different kinetic behavior from the protein-small ligand dissociation described above, where for the corresponding linear free energy relationships ϕ ∼ 1.0.

**Figure 5.**
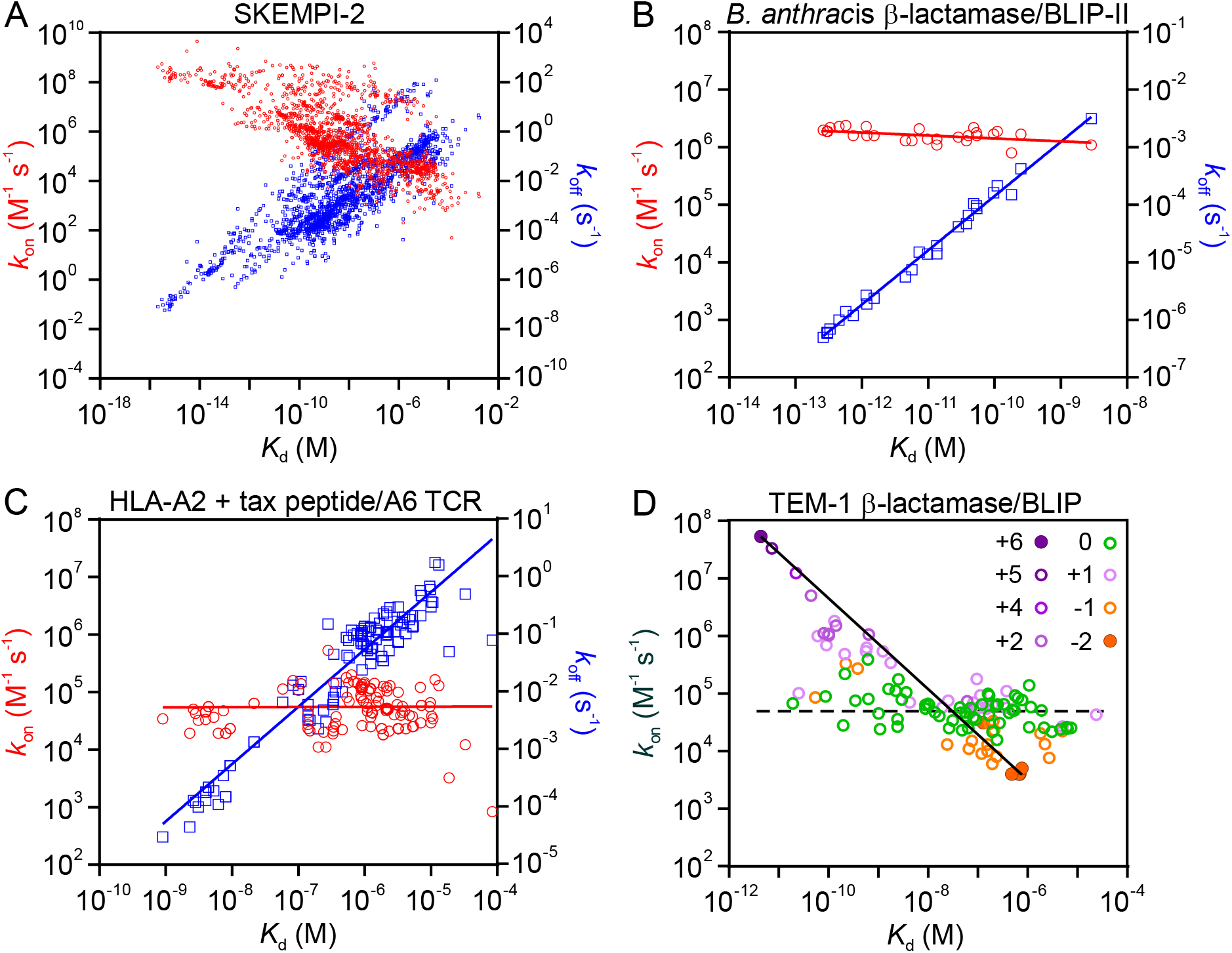
Plots of *k*_on_ and *k*_off_ against *K*_d_ for protein-protein complexes from the SKEMPI-2 database. (A) Global data for all mutants in the SKEMPI-2 database. (B) Mutated forms of *B. anthracis* β-lactamase binding to BLIP-II and (C) HLA-A2/Tax peptide binding to A6 TCR, showing affinity differences dominated by changes in *k*_off_ as typical of most interactions in the database. (D) Plot of *k*_on_ only versus *K*_d_ for mutant forms of TEM-1 β-lactamase binding to BLIP (the corresponding data for *k*_off_ were omitted for clarity), as a rare example of an interaction with a strong dependence of *K*_d_ on *k*_on_. The data are color-coded to indicate the net change in overall charge on the TEM-1 β-lactamase resulting from the mutations. Trend lines are shown to highlight the correlation between charge, *k*_on_ and *K*_d_ for two subsets of mutants, one for which differences in affinity are driven by *k*_on_ (solid line), and the other for which *k*_on_ is invariant (dashed line).

### Implications for the mechanism of ligand dissociation

The observation that for the slow-binding inhibitors of InhA the off-rate is relatively invariant across a significant range of inhibitor affinity values is easily understood if the slow conformational reorganization involved in progression of the intermediate complex, RL, to the final inhibited complex, RL’, is primarily regulated by the intrinsic dynamics properties of the protein. In such cases it is plausible that dissociation of structurally distinct ligands that bind with different affinities might occur with similar rates if the same slow conformational change in the protein is rate-limiting.

For the low affinity binders in **Figures 1-3**, the invariance in *k*_off_ values similarly implies that the activation barrier for *k*_-2_ (**Figure 1D**) depends on dynamic motions of the protein that precede disengagement of the ligand from its binding site (**Figure 6**). Any substantial loss of binding energy with the ligand would differentially destabilize this transition state compared to the bound complex, reducing the off-rate and giving ϕ values substantially less than one, in contradiction to our observations. Our results therefore indicate that ligand can dissociate only from rare, high energy conformational states of the RL complex, that these states form slowly, and that subsequent dissociation of the ligand is fast so that achieving the excited state is rate limiting for *k*_*-2*_. Whether formation of the high energy conformation and disengagement of the ligand are discrete steps, or instead involve a concerted process but with ligand disengagement lagging behind the protein conformational changes, hinges on whether the high energy state has a sufficient lifetime to be considered a discrete intermediate. Our results do not distinguish these possibilities and so this detail remains unknown. A protein-gated mechanism for the ligand’s exit from its binding site likely obtains for high affinity binders as well as low, though it becomes kinetically masked for interactions in which binding is encounter controlled, because when *k*_*1*_ is rate limiting *k*_*-2*_ ceases to influence the observed on- and off-rates. The principle of microscopic reversibility requires that, if ligand can exit the binding site only in high energy states of the protein, then ligand can enter the binding site only in the same high energy conformational states. Thus, during ligand binding, the rate of conversion of the initial encounter complex to the final bound state is also gated by waiting for the protein to adopt the required high energy conformation to open the binding site for ligand entry. The observation that several different target classes display broadly similar affinity-kinetic relationships suggests that this protein gated mechanism for ligand binding and dissociations is a common phenomenon.

**Figure 6.**
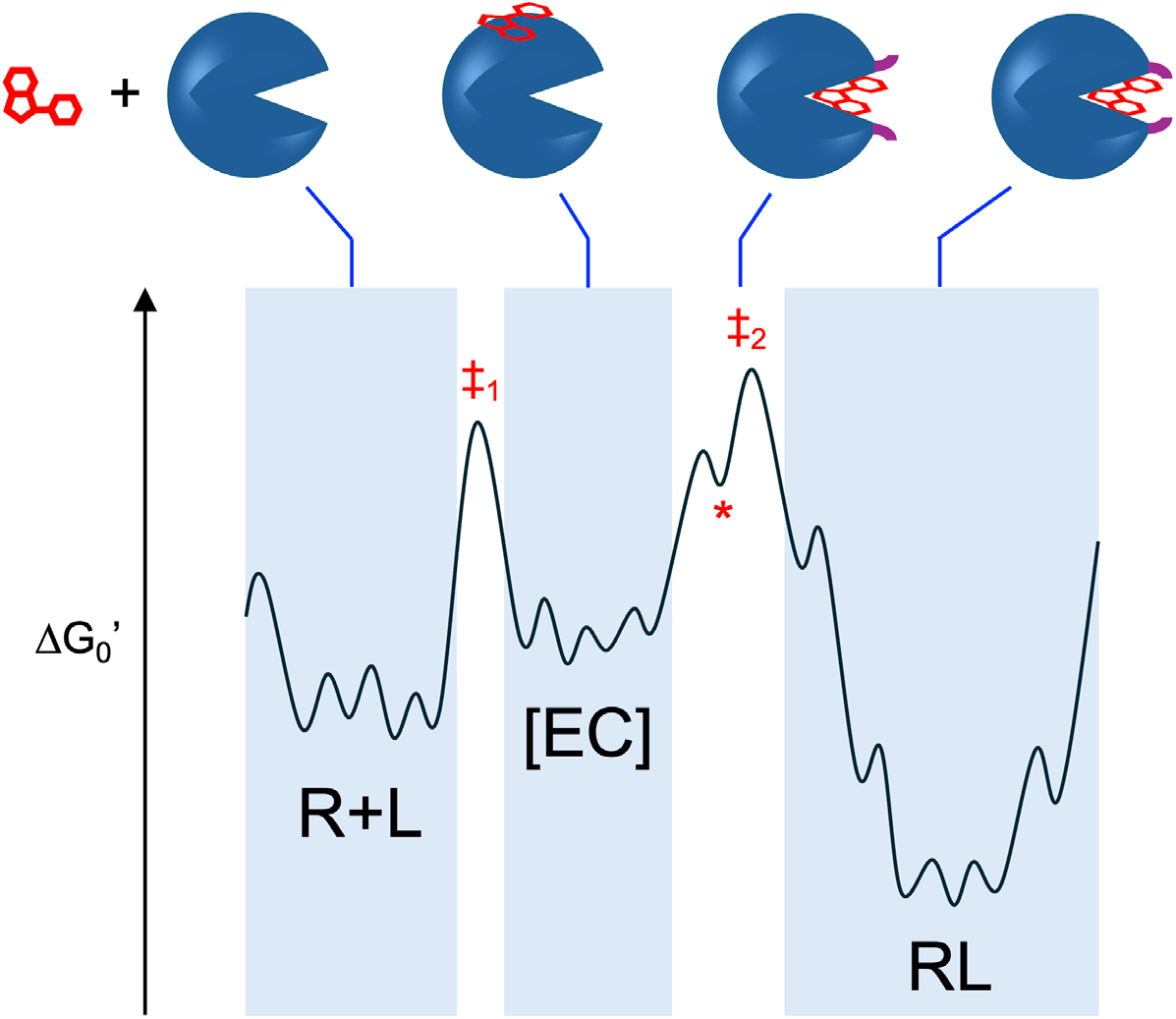
The proposed protein-gated mechanism for ligand binding and dissociation. The free receptor and ligand (R+L) state, the encounter complex ([EC]), and the bound complex (RL) each exist as an ensemble of conformational microstates. The transition states for the formation of [EC] and its subsequent reorganization to give RL are designated ‡_1_ And ‡_2_. Ligand can exit the binding site in RL only from a high energy conformational state (indicated *) in which protein motions have opened a path for the ligand to depart, represented by movement of the purple-highlighted regions in the upper cartoon. Adopting the high energy state is rate-limiting compared to the subsequent rapid exit of the ligand from its binding site, and so in ‡_2_ the ligand is fully bound.

The nature of the conformational changes involved in achieving the high energy, ligand entry/exit-competent state of the protein is unclear. Compared to the ground state RL complex, the high energy state may be enthalpically destabilized by strain or by disruption of internal interactions, may be entropically unlikely due to a requirement for multiple low energy motions to occur in concert to open a path for the ligand to escape, or a combination of both. The protein motions do not, however, substantially disrupt interactions between the protein and the ligand; our measured ϕ values indicate that these interactions remain substantially intact in the transition state. Therefore, involvement of large-scale motions that uncover substantial portions of the ligand surface that were previously in contact with the protein appears unlikely.

Limits can be placed on the free energy of the high energy state of RL by considering that it is likely less stable than the R + L ground state (i.e. formation of the encounter complex is likely energetically up-hill even at 1 M ligand), and must be lower in energy than the transition state for *k*_*-2*_. The former energy difference can be calculated from the *K*_d_ value that defines the threshold between the *k*_on_- and *k*_off_-driven limbs of the structure-kinetic relationship, and the latter can be estimated from *k*_*-2*_ using the Eyring-Polanyi equation. For BTK, therefore, we can say that the high energy state RL* is likely higher in free energy, compared to the RL complex, by somewhere between 11 and 20 kcal/mol. Thus, the ligand entry/exit competent state is indeed rare, comprising only 1 in 10^8^–10^14^ RL complexes. The differences in limiting *k*_off_ values seen for the different protein classes in **Table S2** provide a measure of the variation in the energy barriers for achieving the high energy state. These range over ∼3 logs among the systems we examined, with complexes involving hERG and certain GPCRs showing the highest barriers, while those with HSP90 and kinases with the inhibitor tucatinib involve significantly faster processes. Our results show that the rate of formation of the high energy state depends primarily on the identity of the protein but can also be affected by the nature of the ligand, consistent with the properties of slow-binding inhibitor systems.^49-53^ Hence, the structure-kinetic relationship for a single protein binding to set of homologous ligands (e.g. **Figures 1A, B**) tend to show very little scatter, while those for the same ligand binding to homologous proteins (e.g. **Figures 2B-H**) show considerable variation around the mean behaviour; unsurprisingly, plots that encompass both different proteins and different ligands show the most scatter (e.g. **Figures 2A**, and **3A, B**). In addition, in systems where *k*_-2_ is insensitive to ligand structure, as illustrated in **Figure 1**, this observation implies that conversion of the encounter complex to the final bound state proceeds predominantly through a conformational selection mechanism, while those cases in which *k*_-2_ varies with ligand identity most likely incorporate an element of induced fit, with ligand-specific features influencing the reorganization step.^54^

The requirement for a protein conformational change prior to ligand dissociation may be because small ligands tend to bind in deep pockets that extensively envelop them,^55^ and thus require some flexion on the part of the protein to enable entry into and exit from the tight confines of the binding site. In contrast, many PPI interfaces are relatively open,^56^ allowing disengagement of the binding partners without prior conformational change. The continuum of ϕ values from 0-0.5 observed for PPI dissociation, in contrast to the near-unit values seen with small ligands, may reflect earlier versus later transition states for PPI dissociation in which the protein-protein contacts are disrupted to different extents.^24^

The current study significantly advances our understanding of protein-ligand binding in several ways. Although it has previously been noted that the affinity of small ligands correlates with *k*_on_ in certain circumstances,^7,14^ we provide a theoretical framework for qualitatively and quantitatively understanding this phenomenon. We additionally show that protein-gated ligand dissociation is true not only in occasional rare instances,^17-19,57,58^ but is widespread, applying to a range of classic small molecule drug targets including kinases, GPCRs, and others. In particular, we show that to understand experimentally observed on- and off-rates it is necessary to explicitly consider the initial encounter complex as an intermediate on the pathway for binding, and that the protein exists as an ensemble of conformational microstates at each stage of the binding process. We further show that conversion of the encounter complex to the final bound complex is kinetically gated by the slow formation of particular high energy conformational states that allow the ligand to enter and exit the binding site, and that in the transition state for this process the ligand is essentially fully (≥85%) bound in its final binding site. Finally, we show that this structure-kinetic relationship is mechanistically related to that seen for classic slow-binding inhibitors, but contrasts sharply with the mechanism for dissociation of protein-protein complexes.

These findings have several important implications for the study of ligand binding and the interpretation and application of binding kinetics. For example, with regard to the computational study of protein-ligand dissociation, our findings imply that the degree of ligand displacement from its bound location is not an appropriate reaction coordinate for modelling the dissociation reaction, as is sometimes done in ‘forced MD’-type computations. This is because the most energetically significant events occur along orthogonal, protein related coordinates, before the ligand has significantly moved from its bound position. Instead, the process of ligand dissociation may be best described using a reaction coordinate that involves primarily motions of the protein atoms, leading to the formation of the high energy conformational state(s) from which the ligand can rapidly exit the binding site. In drug discovery, our findings set an expectation that, counter to current dogma, in the early stages of hit optimization, affinity enhancement is achieved primarily by increasing *k*_on_ rather than by decreasing *k*_off_, with the latter becoming true only once affinity has increased to the point where binding is diffusion controlled. Measuring binding kinetics from an early stage, and interpreting them according to the framework described here, can yield important information about drug binding, such as the magnitude of drug-target residence time that might be achieved if a series were optimized to a given affinity level, and provides a means for the clear identification of kinetic outliers that could provide starting points for the development of slow-binding inhibitors. Our findings additionally raise questions that may stimulate future lines of research. For example, for a given protein, what are the conformational changes required to achieve the excited state that allows ligand binding and dissociation, and can these be modeled to allow more accurate computational prediction of ligand on- and off-rates? Are there general principles that determine how one class of proteins differs from another in this respect? How does the structure of the ligand affect the rate of achieving the excited state, and how can an understanding of these effects be used to enable the rational design of slow-binding inhibitors?

## CONCLUSIONS

The binding of proteins with small molecule ligands pervades biology, pharmacology, and medicine. A detailed understanding of how such binding occurs is therefore of broad importance. We show here that the entry and exit of a small ligand from its binding site can occur only in an excited conformational state of the protein. Consequently, over certain affinity ranges, the kinetics of ligand binding and dissociation are regulated by protein dynamics independent of the strength of ligand binding. We demonstrate this phenomenon across a diversity of protein and ligand classes, suggesting that the mechanism of protein-gated binding and dissociation may be quite general. The above conclusions represent a significant departure from the conventional view that ligand dissociation is best thought of in terms of incremental disengagement from the binding site. Instead, we show that to understand experimentally observed ligand on- and off-rates it is necessary to consider the bound complex as an ensemble of conformational microstates, with the rate-limiting step in a ligand’s entry to or departure from its binding site being the slow adoption an excited conformational state that creates a path for the subsequent, rapid movement of the ligand. In the transition state for this process, the ligand remains essentially fully bound in its original position. The binding of small ligands in general is therefore mechanistically related to that of classical slow-binding inhibitors, but quite distinct from that of most protein-protein complexes.

## MATERIALS AND METHODS

### High-Throughput Kinetic Screening Assay

TR-FRET binding assays conducted using the KINETICfinder platform were performed as described previously.^20^ In brief, experiments were performed at room temperature in black 384 well microplates using a buffer (50 mM HEPES, pH 7.5) containing 10 mM MgCl_2_, 0.01% Brij-35, 1 mM DTT. Kinases (**Table S2**) were incubated with Tb-streptavidin, Tb-anti-GST or Tb-anti-His antibody and kinase tracers. For all experiments, a 4-point 10-fold serial dilution of 100x concentrated test compounds were prepared in DMSO. Positive control wells contained no test compound and negative control wells contained no kinase. The kinetic assays were read continuously in a PHERAstar FSX plate reader (BMG LABTECH) and the specific TR–FRET signals were fitted to the Motulsky-Mahan equation. The affinity (*K*_d_), kinetic constants (*k*_on_, *k*_off_) and residence time of each test compound were calculated using KINPy^®^ software (Enzymlogic). Errors in the calculated values are SEM. Off-rates were set to > 0.5 s^-1^ if values are above assay quantitation limit. The kinetic data for seven reported kinase inhibitors, danusertib (ApexBio), dasatinib monohydrate (Santa Cruz), encorafenib (ApexBio), foretinib (MedChemExpress), tivozanib (ApexBio), tucatinib (MedChemExpress) and zimlovisertib (MedChemExpress) for 134, 136, 135, 123, 133, 87 and 86 kinases, respectively, are reported in **Tables S3-S9**. A subset of the kinetic data obtained using the KINETICfinder TR-FRET binding assay is compared with values obtained using the orthogonal method of SPR in **Figures S5** and **S6, Table S11**, showing good agreement between the two methods. The SPR data was either taken from the literature or determined here for BTK (see below)

### Surface Plasmon Resonance (SPR) Kinetics

#### Cloning and expression of BTK

SPR kinetics were performed using the kinase domain from mouse BTK (UNIPROT CODE: P35991) which is 98.9% identical to the human enzyme (UNIPROT CODE: Q06187), and includes the Y617P mutation necessary to make the protein soluble in *E. coli*. The cDNA encoding the sequence for the catalytic domain (396-659) of mouse BTK Y617P containing a TwinStrep^®^ tag at the C-terminus was purchased from GenScript and inserted into a pET-24a(+) plasmid. BL21 (DE3) cells were co-transformed with the BTK plasmid (100 ng/µL) together with a plasmid encoding the *Y. enterocolitica* tyrosine-protein phosphatase YopH (100 ng/µL) using the heat-shock method (30 s, 42°C). The transformed cells were plated on LB-agar plate containing 50 µg/mL of kanamycin and 100 µg/mL of spectinomycin, and incubated overnight at 37°C. Subsequently, one colony was picked and used to inoculate 150 mL of LB starter culture containing both antibiotics, which was then incubated overnight at 37°C with shaking at 220 rpm. The overnight culture was then used to inoculate (1:50) LB media which was incubated at 37°C (200 rpm) until the OD_600_ reached 0.6-0.8. The flasks were chilled on ice for approximately 30 min and then induced with 0.4 mM of IPTG at 18°C for 20 h. Cells were harvested by centrifugation at 5000g for 15 min and the cell pellets stored at -80°C.

#### Protein purification

All procedures were carried out on ice or at 4°C. The cell pellet generated by overexpressing mBTKY617P (396-659)-TwinStrep^®^ tag was resuspended in lysis buffer (100 mM Tris/HCl, pH 8, 5% glycerol with a mixture of proteases inhibitors (GoldBio) and DNase I (Sigma Aldrich), cells were lysed using sonication, and the resulting lysate was centrifuged at 45,000g for 1 h. The supernatant was loaded onto a 1ml StrepTactin^®^ XT 4Flow^®^ gravity column previously equilibrated with wash buffer (100 mM Tris/HCl pH 8, 150 mM NaCl, 1 mM EDTA, 10% glycerol) and washed with 5CV of wash buffer. Protein was then eluted using 3CV of elution buffer (100 mM Tris/HCl pH 8, 150 mM NaCl, 50 mM biotin, 1 mM EDTA, 10% glycerol), concentrated to 1 mL and loaded onto a HiLoad 16/600 Superdex™ 75 column preequilibrated with gel-filtration buffer (20 mM HEPES pH 8, 300 mM NaCl, 5% glycerol). Fractions containing mBTKY617P (396-659)-TwinStrep^®^ were combined, concentrated and stored at -80°C. Protein concentration was determined using UV-Vis nanospectroscopy (ND-1000, ThermoFisher), and protein purity was assessed using SDS-PAGE.

#### Surface Plasmon Resonance

SPR experiments were performed on a Biacore T200 Instrument (GE Healthcare) at 25°C. A solution of 50 µg/mL StrepTactin^®^ XT (IBA Lifescience) solution in 10 mM sodium acetate pH 4.5, was immobilized on a Series S CM5 Chip (Cytiva), that was previously activated for 10 min using a 1:1 mixture of 0.1M EDC(3-(N,N-dimethylamino) propyl-N-ethylcarbodiimide) and 0.1 M (N-hydroxysuccinimide) NHS, using a flow rate of 10 µL/min. The ligand was immobilized at a density of about 2500 RU, and the surface was then deactivated using 1 M of ethanolamine and conditioned using 10 mM of NaOH.

TwinStrep^®^ tagged BTK (100 nM in HBS-P supplemented with 0.5% DMSO) was captured on the chip at a flow rate of 10 µL/min, resulting in a final surface density of about 1000-2000 RU. Binding experiments were performed using Single-Cycle Kinetic (SCK) mode. A 5-point serial dilution of compounds were prepared in HBS-P + 0.5% DMSO and were injected at a flow rate of 50 μL/min. The surface was regenerated using 30 s injection of 50 mM NaOH. Data were fitted using the Biacore T200 Evaluation software using a 1:1 model or a two-state model (**Figure S6**). The SPR kinetic constants are given in **Table S11** and plotted together with the TR-FRET and SPR data for other kinases in **Figure S5**.

### The KIND Dataset

The KIND dataset comprising 78 biological targets spanning kinases, GPCRs, heat shock proteins (HSPs), selected enzymes, and ion channels with a total of 3,812 interactions of which 3,685 were assessed including *k*_on_, *k*_off_ and *K*_d_) was downloaded from https://kbbox.h-its.org/toolbox/data/kind-dataset/.

### The SKEMPI-2 Dataset

The SKEMPI-2 database containing equilibrium and/or kinetic data for a total of 7,085 wild-type and mutant proteins involved in protein-protein or protein-peptide binding was downloaded from https://life.bsc.es/pid/skempi2.^48^ Entries that lacked *k*_on_ or *k*_off_ values were discarded, and 1,561 entries encompassing mutational variations of 66 distinct pairs of interaction partners were analyzed. Those interactions represented by the largest number of mutational variants were examined individually (**Figure S4**) with three examples included in **Figure 5**.

### Statistical Tests

The datasets were assessed for their normality/lognormality using D’Agostino-Pearson, Anderson-darling, Shapiro-Wilk and Kolmogorov-Smirnov tests. All subsequent statistical analysis, performed in GraphPad Prism version 10.1.2, were determined by the nature of distribution. As expected, the datasets corresponding to *K*_d_, *k*_on_ and *k*_off_ were lognormally distributed. Spearman correlation analysis was performed taking into account the non-gaussian nature of distribution for some parameters in the dataset.

## Abbreviations

BTK: Bruton’s tyrosine kinase
TR-FRET: time-resolved Förster resonance energy transfer
GPCR: G-protein coupled receptors
HSP90: Heat shock protein-90
K4DD: kinetics for drug discovery

## ACKNOWLEDGMENTS

The research was supported by the National Institutes of Health grant GM149297 to PJT and by the Centre for the Development of Industrial Technology (CDTI) through the Horizon Europe EIC Accelerator Seal of Excellence SME grant (SoE-20211014) to AC and PA. ASR was supported by a National Institutes of Health Chemistry-Biology Interface Training Grant (GM136572), and MB was supported by an Abroad Fellowship (31189) from Associazione Italiana per la Ricerca sul Cancro (AIRC). The authors thank Rachel Grimley for her constant support, enthusiasm and critical input, and Joanna Brookfield, Helena Danielson and Fredrik Svensson for helpful discussions. Rachel Grimley is also acknowledged for reading the manuscript and providing comments that have substantially improved the narrative.

## CONFLICT OF INTEREST STATEMENT

The authors declare no conflicts of interest.

## Notes

### Competing Interest Statement

The authors have declared no competing interest.

