## Supplementary Information for "Protein-ligand binding kinetics are primarily controlled by the protein, not the ligand"

[#]These authors contributed equally

| **Figures/Tables** | **Content** | **Page** |
|---|---|---|
| Figure S1 | Binding kinetic data for BTK at 37°C (TR-FRET, KINETICfinder). | S3 |
| Figure S2 | Phylogenetic kinase tree. | S4 |
| Figure S3 | Binding kinetic data for GPCRs from the KIND database. | S5-S6 |
| Figure S4 | Binding kinetic data for protein-protein interactions from the SKEMPI 2.0 database. | S7-S11 |
| Figure S5 | Comparison of binding kinetic data determined using TR-FRET (KINETICfinder) and SPR. | S12 |
| Figure S6 | SPR data for BTK binding kinetics. | S13 |
| Table S1 | Protein-ligand data sets used in this study | S14 |
| Table S2 | Kinases used in this study | S15-S16 |
| Table S3 | TR-FRET (KINETICfinder) binding kinetic data: Danusertib | S17-S18 |
| Table S4 | TR-FRET (KINETICfinder) binding kinetic data: Dasatinib | S19-S20 |
| Table S5 | TR-FRET (KINETICfinder) binding kinetic data: Encorafenib | S21-S22 |
| Table S6 | TR-FRET (KINETICfinder) binding kinetic data: Foretinib | S23-S24 |
| Table S7 | TR-FRET (KINETICfinder) binding kinetic data: Tivozenib | S25-S26 |
| Table S8 | TR-FRET (KINETICfinder) binding kinetic data: Tucatinib | S27-S28 |
| Table S9 | TR-FRET (KINETICfinder) binding kinetic data: Zimlovisertib | S29-S30 |
| Table S10 | Limiting $k_{on}$ and $k_{off}$ values and affinity thresholds for the different target classes and inhibitors | S31 |
| Table S11 | SPR and TR-FRET (KINETICfinder) binding kinetic data. | S32-S33 |
| References | | S34-S35 |

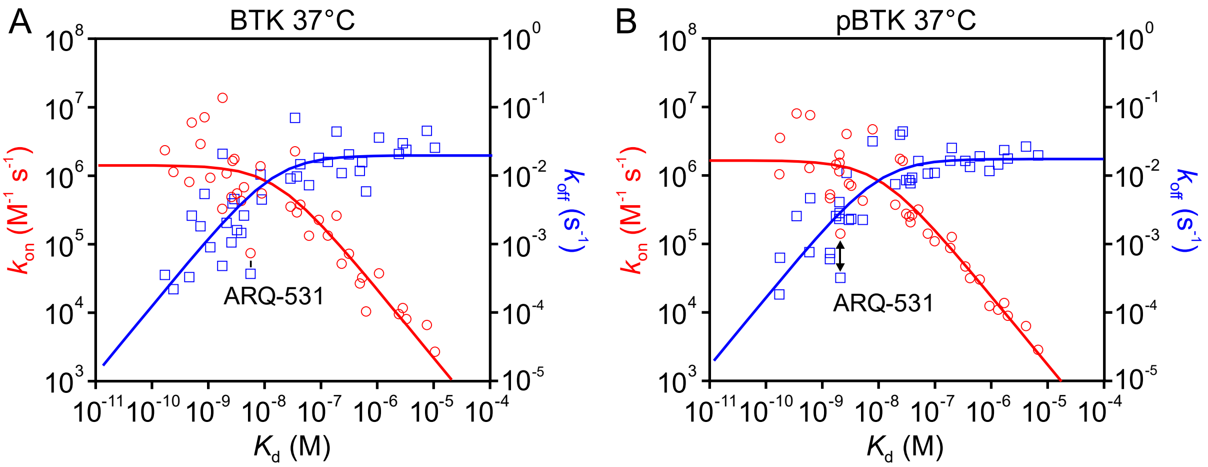

**Figure S1**. Plots of $k_{on}$ (red circles) and $k_{off}$ (blue circles) against $K_d$ for 36 small molecules binding to (**A**) unphosphorylated BTK and (**B**) phosphorylated BTK at 37°C.

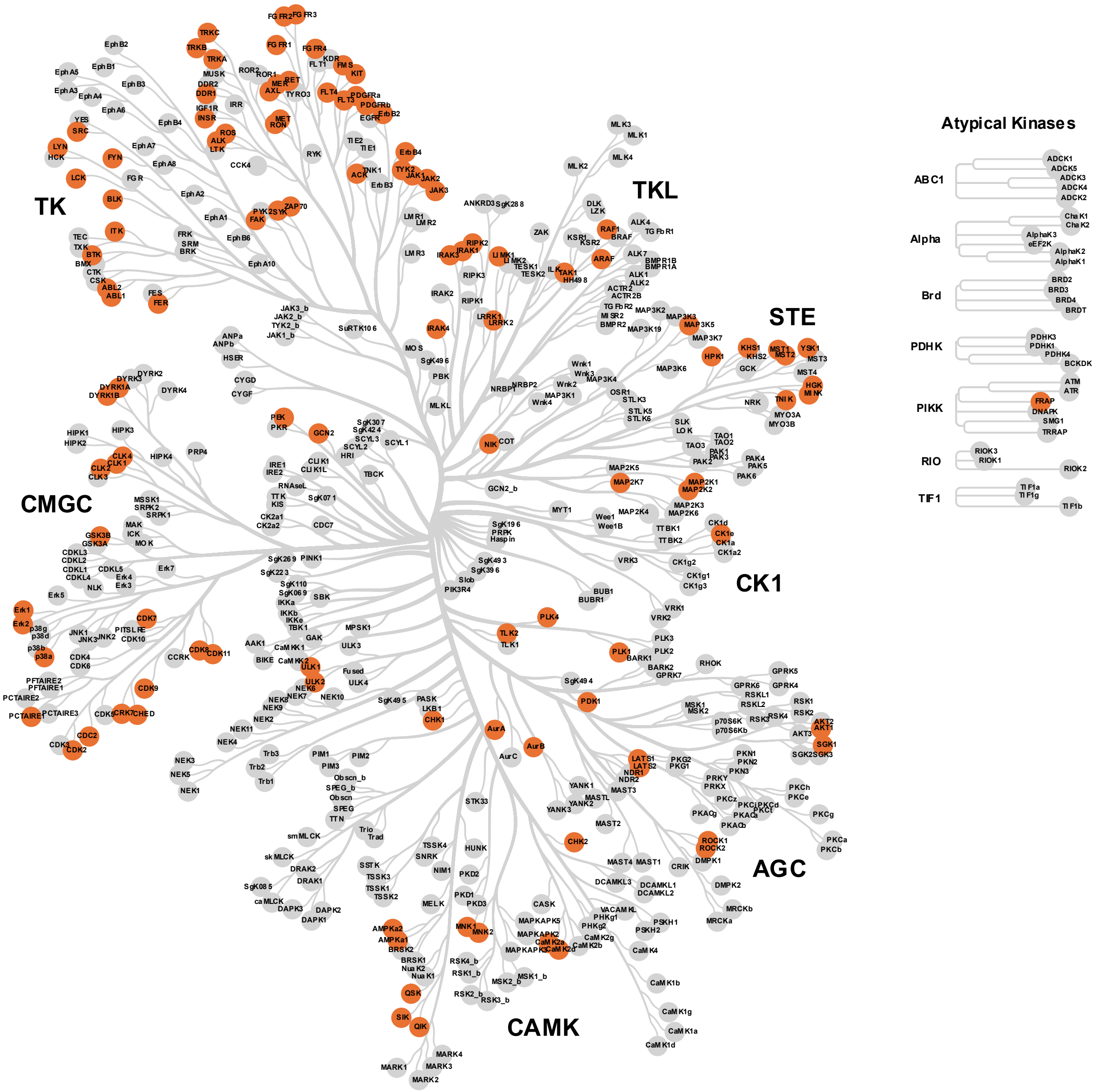

**Figure S2.** Phylogenetic tree of different kinases employed in this study (orange). The kinome tree was generated using the CORAL package.

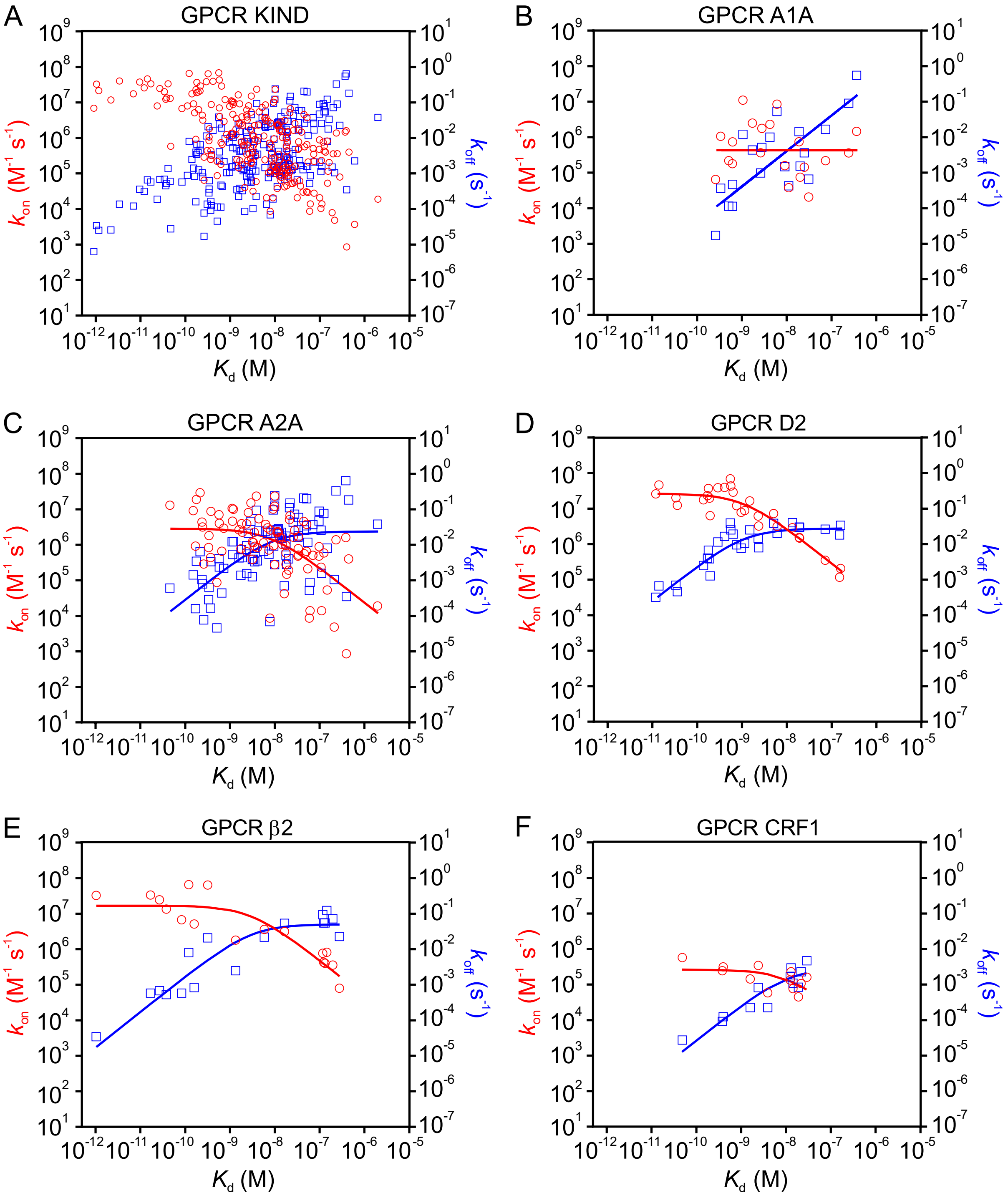
A
GPCR KIND
B
GPCR A1A
C
GPCR A2A
D
GPCR D2
E
GPCR β2
F
GPCR CRF1
$k_{on}$ ($M^{-1}$ $s^{-1}$)
$k_{off}$ ($s^{-1}$)
$K_d$ (M)

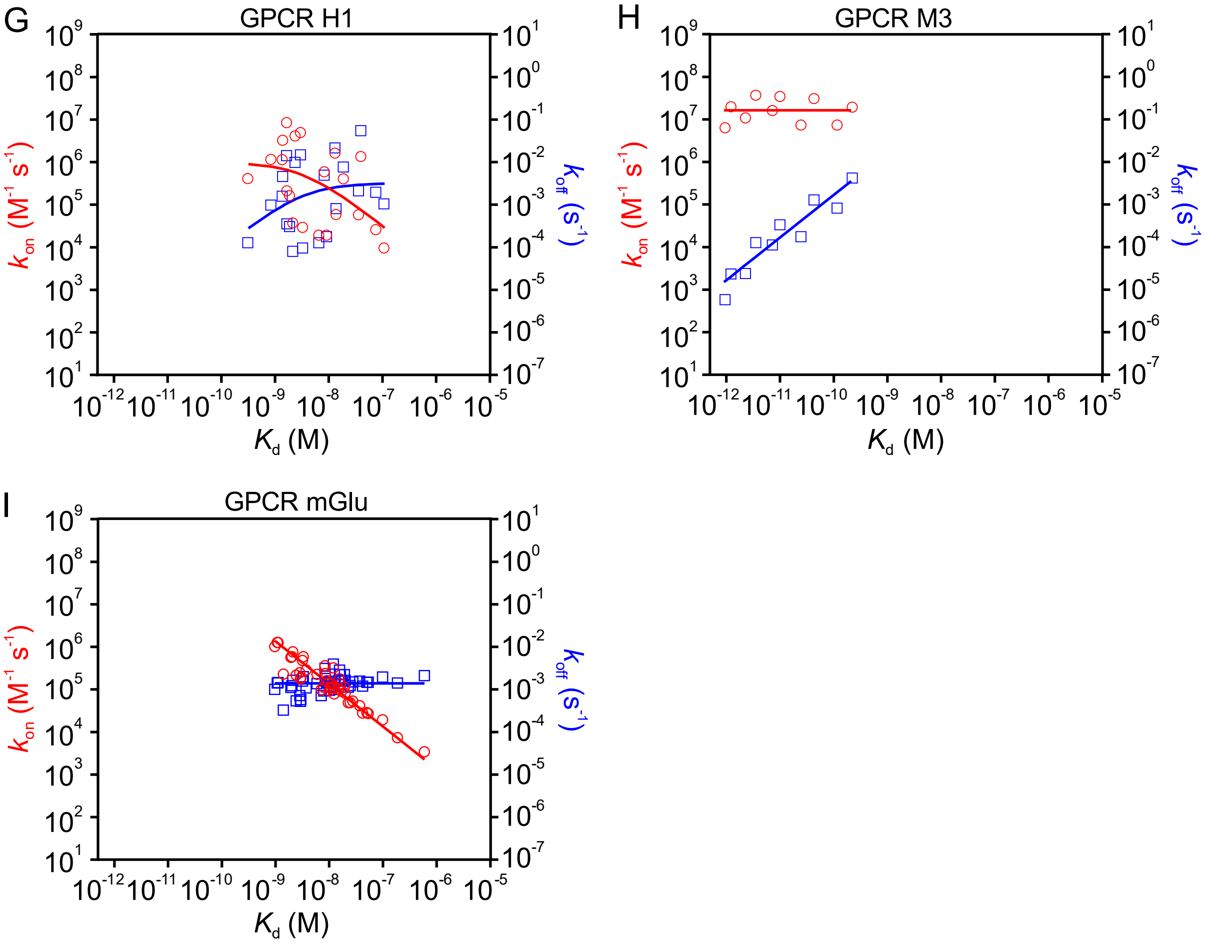

**Figure S3.** Scatter plot of binding kinetic data for GPCRs from the KIND data set.

**Figure S4.** Binding kinetic data for protein-protein interactions.
**A) Affinity changes driven primarily by changes in $k_{off}$**

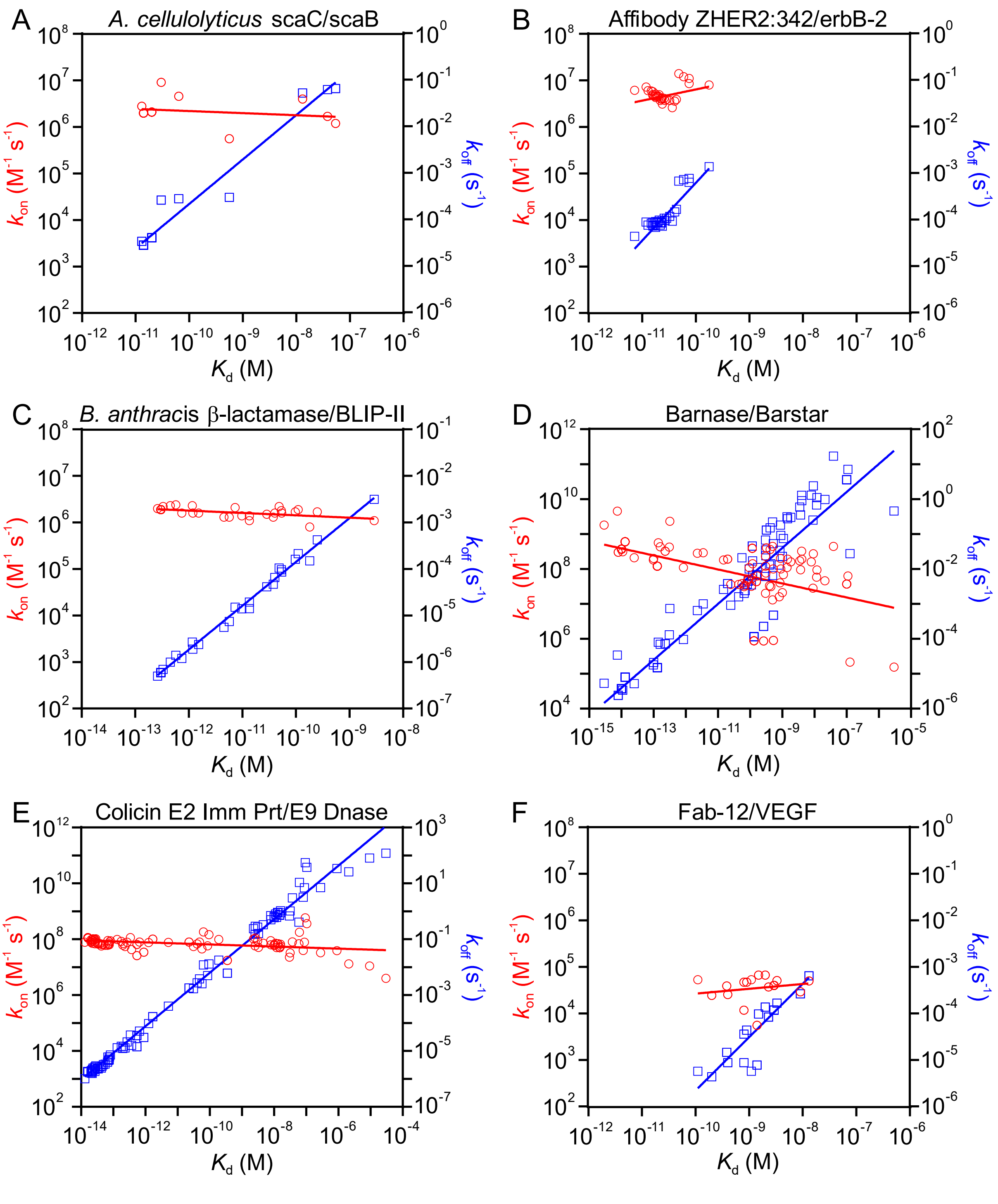

**A) Affinity changes driven primarily by changes in $k_{off}$ (continued)**

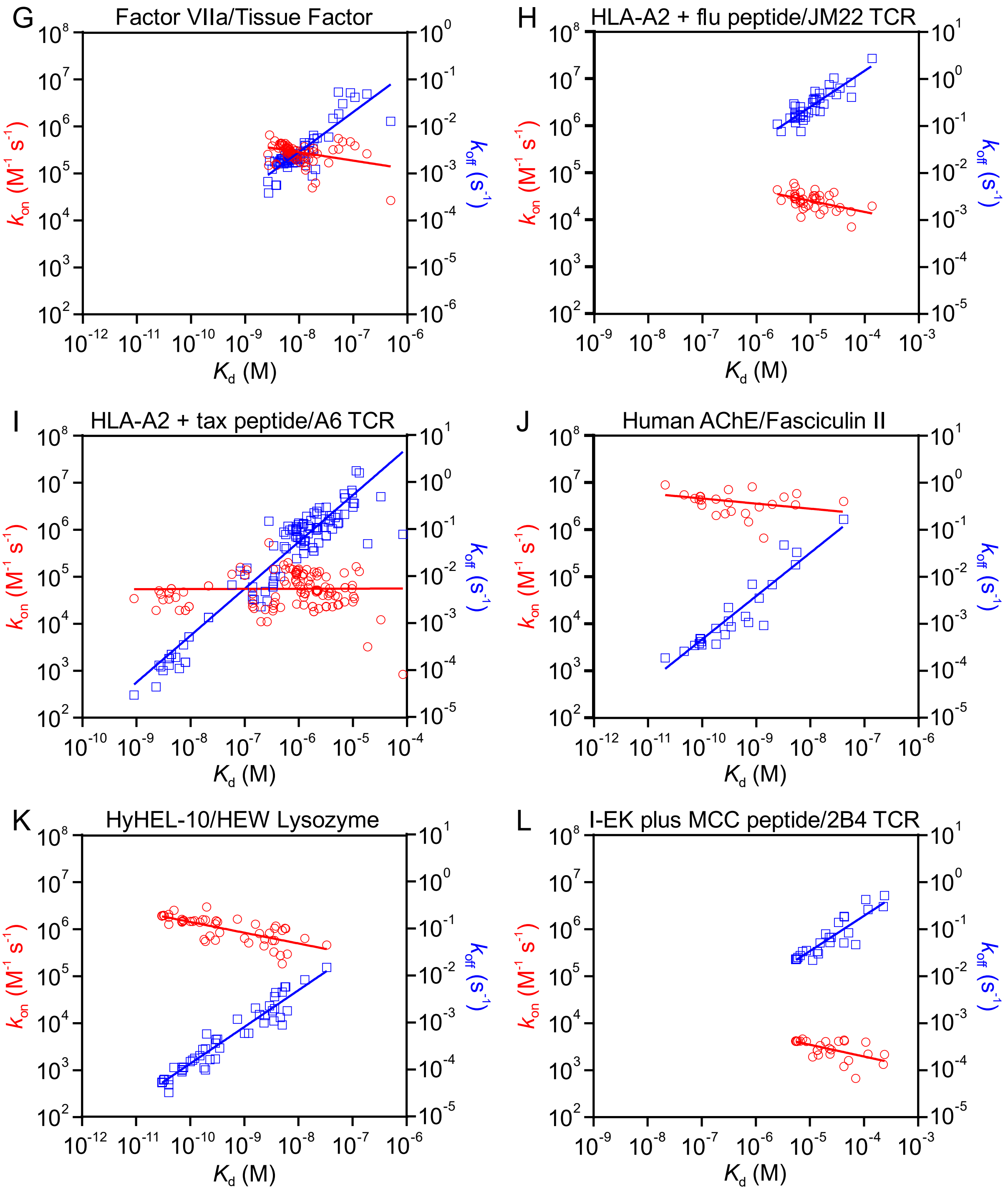

### A) Affinity changes driven primarily by changes in $k_{off}$ (continued)

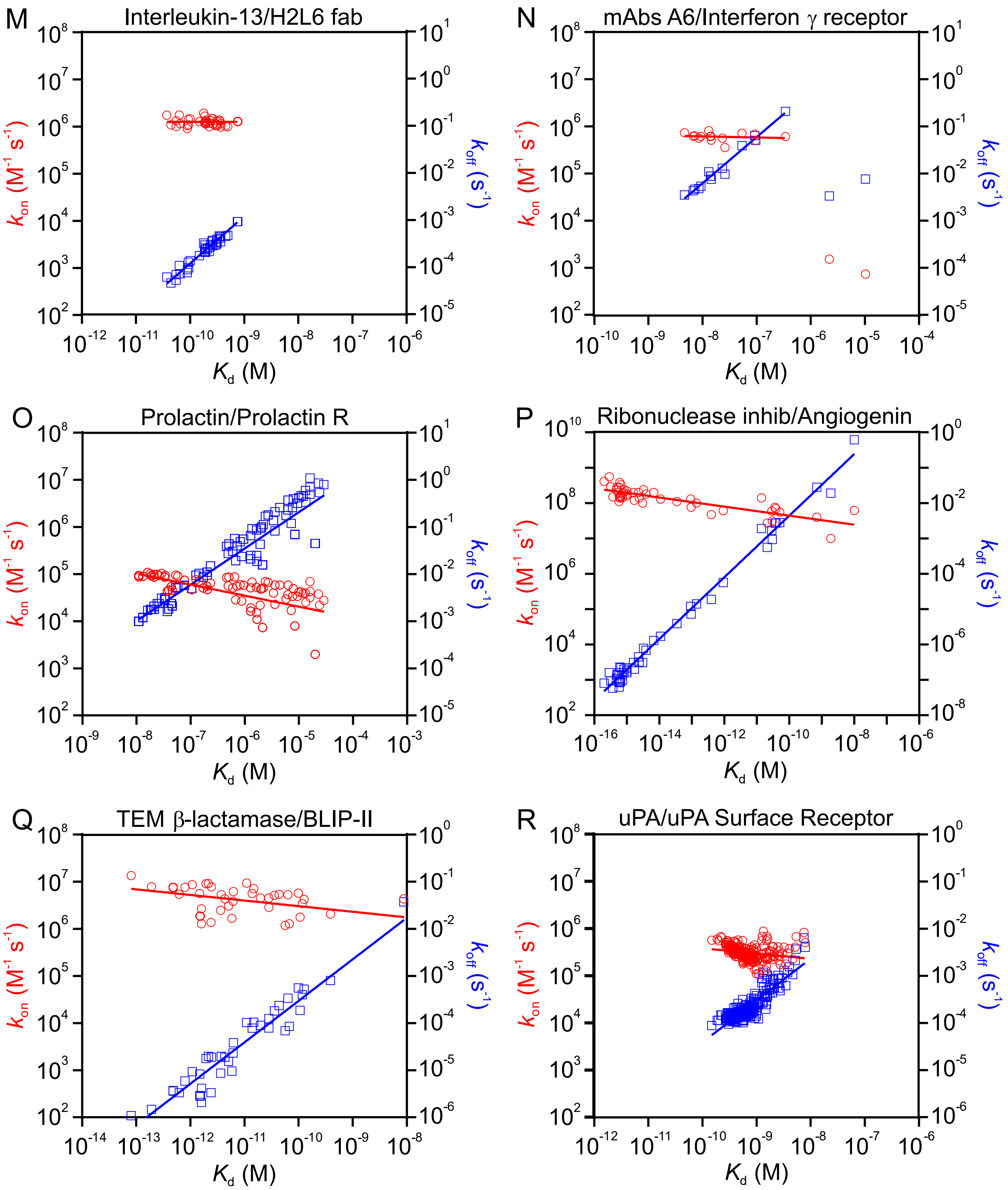

**B) Affinity changes driven primarily by changes in $k_{on}$**

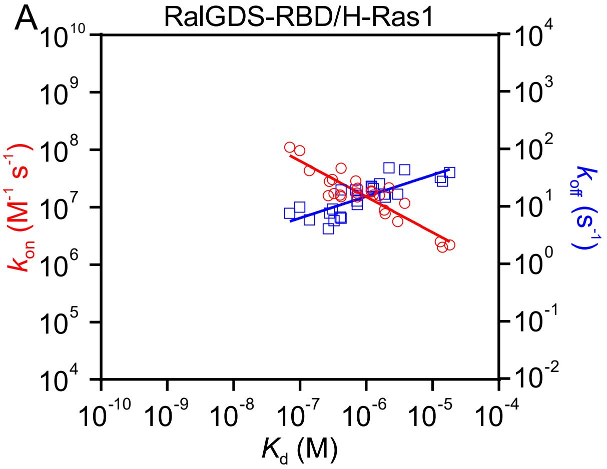

### C) Affinity changes driven by changes in both $k_{on}$ and $k_{off}$

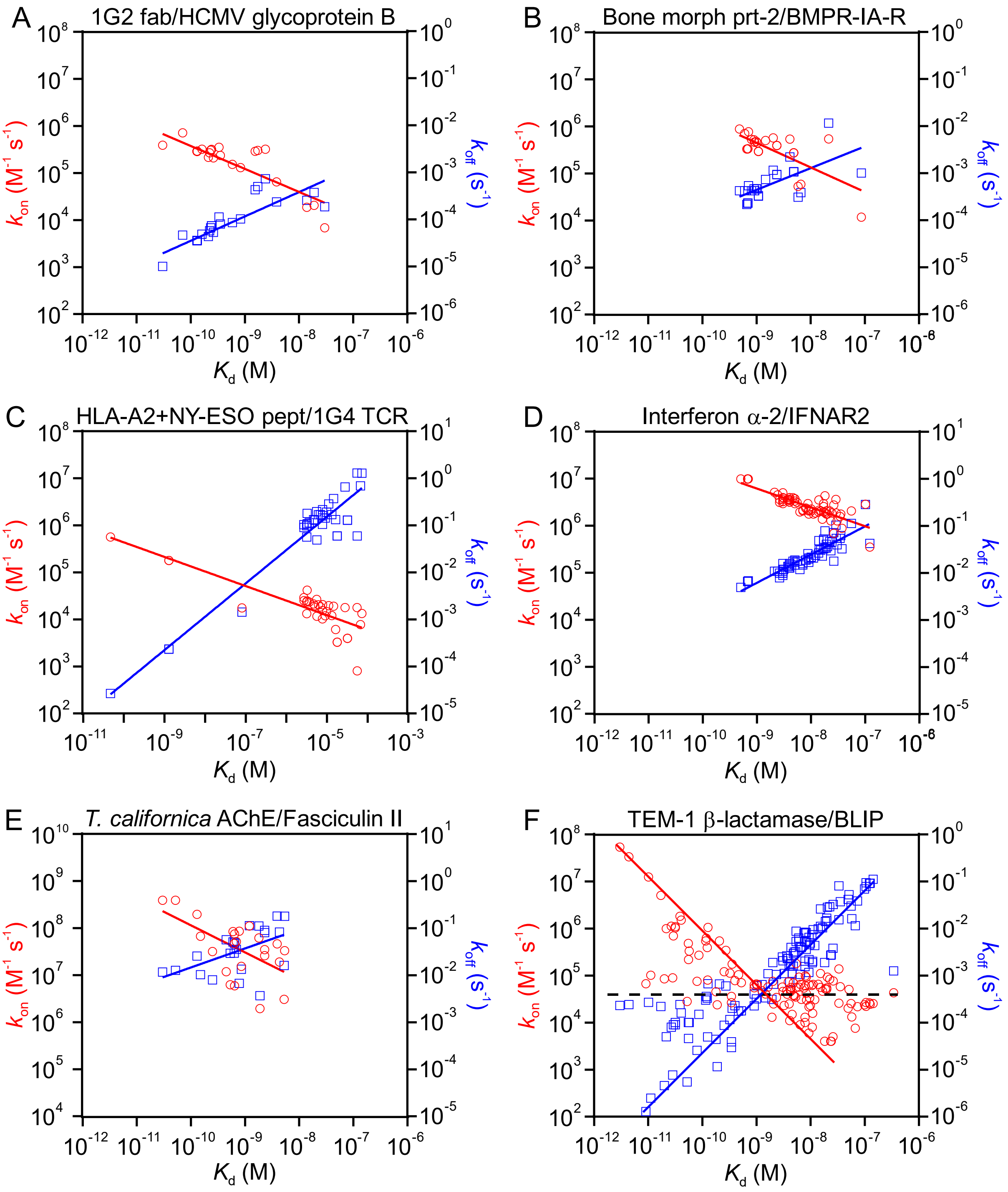

**Figure S4.** Binding kinetic data for protein-protein interactions. A) Affinity changes driven primarily by changes in $k_{off}$. B) Affinity changes driven primarily by changes in $k_{on}$. C) Affinity changes driven by changes in both $k_{on}$ and $k_{off}$. Data were taken from the SKEMPI 2.0 database.[1] In Figure S4C panel F (TEM-1 β-lactamase/BLIP) lines with unit or zero slope are shown to highlight the data trends.

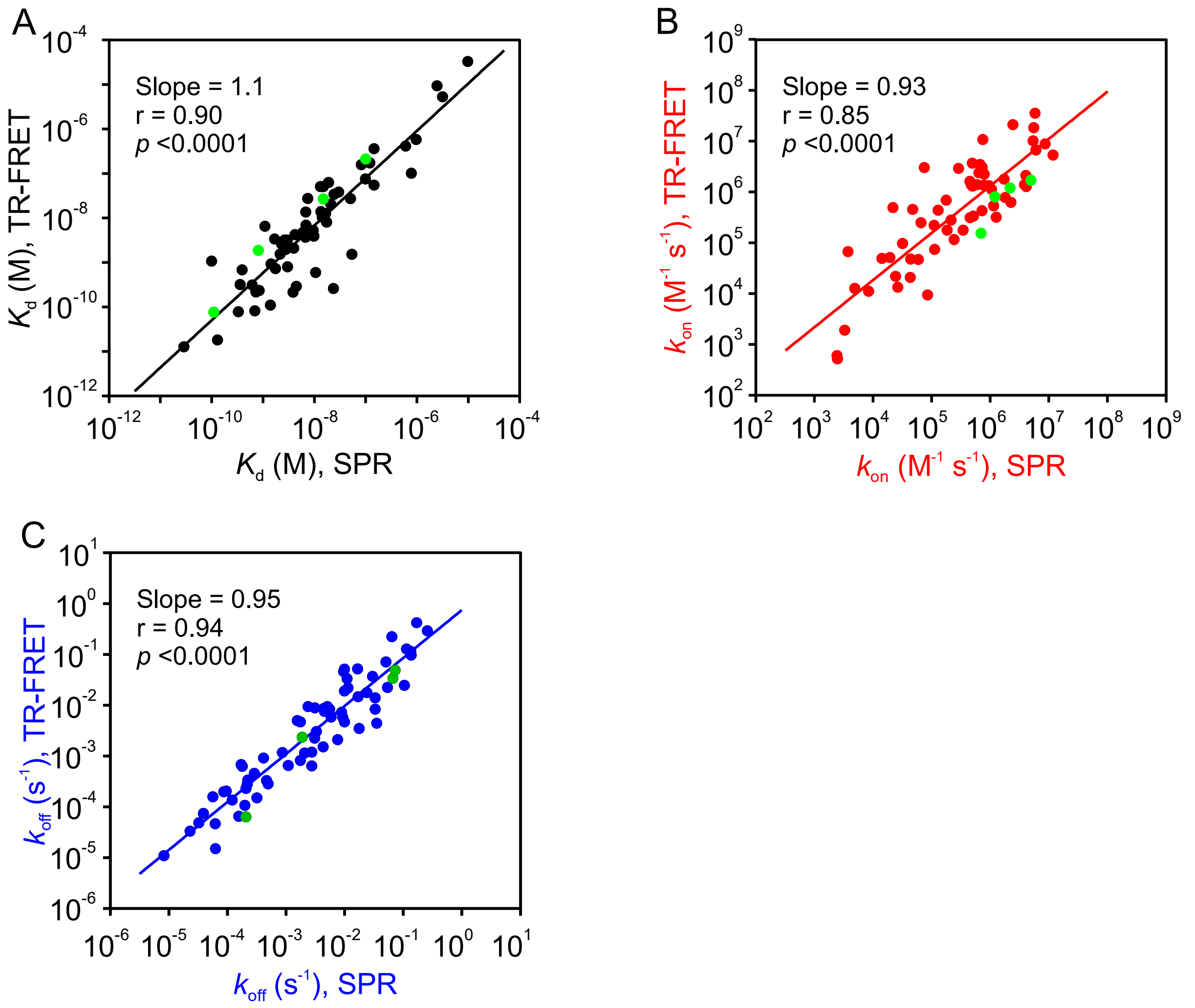

**Figure S5**. Comparison of SPR and TR-FRET (KINETICfinder) binding kinetic data. For A) $K_d$ and B) $k_{on}$ there are 63 pairs of values, whereas for C) $k_{off}$ there are 70 pairs. The slopes and statistical parameters are given in the figures and include the Pearson correlation (r) and two-tailed $p$-value. Data for the unphosphorylated BTK (25°C) were determined in this study and include GDC-0853, GDC-0834, BTK-In-1, P301390 (green). The remaining data were taken from published sources (**Table S11**).

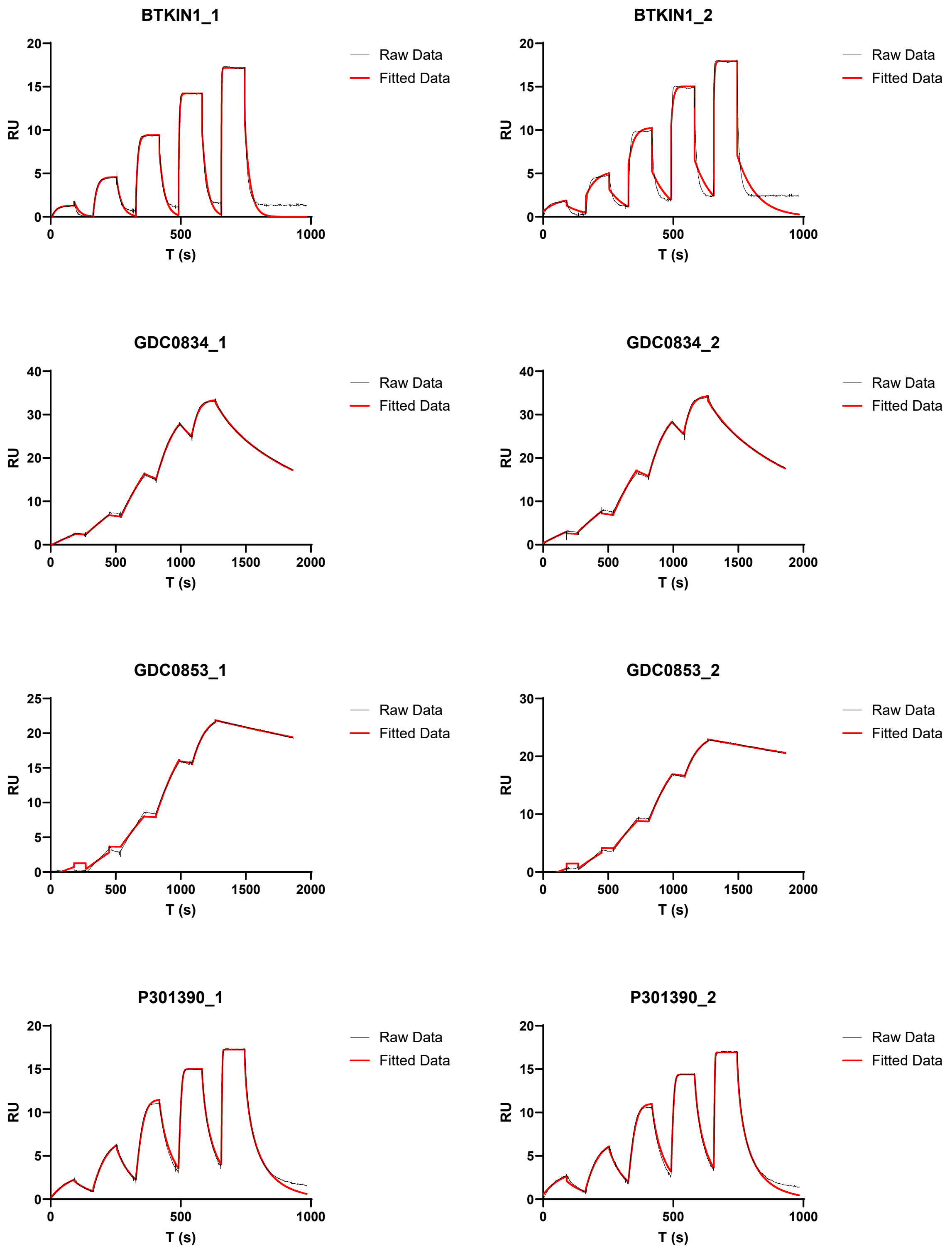

**Figure S6.** SPR data for BTK binding kinetics.

**Table S1.** Protein-ligand and protein-protein interaction datasets used in this study.[a]

| Data Type | Subclassification | Interactions with | Source |
|---|---|---|---|
| Bruton's tyrosine kinase inhibitors | unphosphorylated 25 °C | 38 small molecules | *Bravo et al.*[2] |
| | phosphorylated 25 °C | 38 small molecules | |
| | unphosphorylated 37 °C | 37 small molecules | |
| | phosphorylated 37 °C | 38 small molecules | |
| | | ∑ = 151 kinase-ligand pairs | |
| Kinase inhibitor promiscuity panel | danusertib | 108 kinases | Generated for this study (**Tables S3-S9**) |
| | dasatinib | 89 kinases | |
| | encorafenib | 55 kinases | |
| | foretinib | 64 kinases | |
| | tivozanib | 56 kinases | |
| | tucatinib | 19 kinases | |
| | zimlovisertib | 64 kinases | |
| | | ∑ = 455 kinase-ligand pairs | |
| KIND database (K4DD)[b] | kinases | 3,238 small molecules | *Schuetz et al.*[3-5] |
| | GPCRs | 242 small molecules | |
| | heat shock proteins | 160 small molecules | |
| | voltage gated ion channel | 45 small molecules | |
| | | ∑ = 3,685 protein-ligand pairs | |
| InhA | Slow binding inhibitors | 20 small molecules | *Rawat et al.*[6] *Spagnuolo et al.*[7] |
| **Grand Total** (protein-small ligand systems) | | **4,311** protein-ligand pairs = **8,622** independent kinetic measurements | |
| SKEMPI2 Database | Protein-Protein interactions involving wild-type and mutational variants of 25 different proteins.[c] | 1,561 wild-type and mutational variants of the corresponding binding partners | *Jankauskaite et al.*[1] |
| **Grand Total** (protein-protein systems) | | **1,561** protein-protein interaction pairs = **3,122** independent kinetic measurements | |

[a]Parameters that either could not be estimated or were estimated only as lower or upper limits were excluded from the analysis presented. [b]The KIND database contains data for 3,812 interactions of which 3,685 are analyzed here. [c]Reflects only those interactions represented by with a sufficient number of mutational variants to warrant inclusion in the current analysis.

**Table S2.** Table of kinases used in this study.

| Target Name | GenBank Accession Number | Construct | Activation Procedure |
|---|---|---|---|
| ABL1 | NP_005148.2 | Partial length | No special measures |
| ABL2 (ARG) | NP_009298 | Full length | No special measures |
| ACK (TNK2) | NP_005772.3 | Partial length | No special measures |
| AKT1 | NP_005154.2 | Full length | Activated by coexpression with PDK1 |
| ALK | NP_004295.2 | Partial length | Activated in vitro via autophosphorylation |
| ALK1 (ACVRL1) | NP_000011.2 | Partial length | No special measures |
| ALK5 (TGFBR1) | NP_004603 | Partial length | Activated in vitro via autophosphorylation |
| AMPK (a1/b1/g1) (activated) | NP_006242.4/NP_006244.2/NP_002724.1 | Full length | Activated in vitro by CAMKK1 |
| AMPK (a2/b1/g1) | NP_006244.2/NP_006244.2/NP_002724.1 | Full length | Activated in vitro by CAMKK1 |
| ARAF | NP_001645.1 | Partial length | Constitutive kinase activity |
| ASK1 (MAP3K5) | NP_005914 | Full length | No special measures |
| AURKA (AURORA A) | NP_940839 | Full length | Activated in vitro via autophosphorylation |
| AURKB (AURORA B)/INCENP | NP_004208/AAU04398.1 | Full length | Activated in vitro via autophosphorylation |
| AXL | NP_068713.2 | Partial length | No special measures |
| BLK | NP_001706 | Full length | No special measures |
| BRAF | NP_004324.2 | Full length | Activated in vitro via autophosphorylation |
| BRAF V600E | NP_004324.2 | Partial length | Activated in vitro via autophosphorylation |
| BTK (activated) | NP_000052 | Full length | Activated in vitro via autophosphorylation |
| BTK (non-activated) | NP_000052 | Full length | No special measures |
| CAMK2a | NP_741960 | Full length | No special measures |
| CAMK2d | NP_742113.1 | Full length | No special measures |
| CDK1/cyclin B1 | NP_001777/NP_114172 | Full length | Activated by coexpression with cyclin B1 |
| CDK12/cyclin K | NP_057591.2/NP_001092872.1 | Partial length | Activated by coexpression with cyclin K and CAK1 |
| CDK13/cyclin K | NP_003709.3/NP_001092872.1 | Partial length | Activated by coexpression with cyclin K |
| CDK16/cyclin Y | NP_006192.1/NP_659449.3 | Full length | Activated by coexpression with cyclin Y |
| CDK19/cyclin C | NP_055891.1/NP_005181.2 | Full length | Activated by coexpression with cyclin C |
| CDK2/cyclin A2 | NP_001789.2/NP_001228.1 | Full length | Activated by coexpression with cyclin A2 |
| CDK2/cyclin E1 | NP_001789.2/NP_001229.1 | Full length | Activated by coexpression with cyclin E1 |
| CDK7/cyclin H/MNAT1 | NP_001790.1/NP_001230.1/NP_002422.1 | Full length | Activated by coexpression with cyclin H and MNAT1 |
| CDK8/cyclin C | NP_001251/NP_005181 | Full length | Activated by coexpression with cyclin H and in vitro via autophosphorylation |
| CDK9/cyclin T1 | NP_001252/NP_001231 | Full length | Activated by coexpression with cyclin T1 |
| CHEK1 (CHK1) | NP_001265.2 | Full length | No special measures |
| CHEK2 (CHK2) | NP_009125 | Full length | No special measures |
| CK1e (CSNK1E) | NP_001885 | Full length | No special measures |
| CLK1 | NP_004062 | Partial length | No special measures |
| CLK2 | NP_003984 | Full length | No special measures |
| CLK4 | NP_065717 | Full length | No special measures |
| CRAF (RAF1) | NP_002871 | Partial length | Constitutive kinase activity |
| CSF1R (FMS) | NP_005202 | Partial length | Activated in vitro via autophosphorylation |
| DDR1 (activated) | NP_001945 | Partial length | Activated in vitro via autophosphorylation |
| DDR1 (non-activated) | NP_001945 | Partial length | No special measures |
| DYRK1A | NP_001387 | Full length | No special measures |
| DYRK1B | NP_004705 | Full length | Activated in vitro via autophosphorylation |
| EGFR (ERBB1) (activated) | NP_005219.2 | Partial length | Activated in vitro via autophosphorylation |
| EGFR (ERBB1) (non-activated) | NP_005219.2 | Partial length | No special measures |
| EGFR (ERBB1) E746_A750del | NP_005219.2 | Partial length | No special measures |
| EGFR (ERBB1) L858R | NP_005219.2 | Partial length | Activated in vitro via autophosphorylation |
| ERK1 (MAPK3) | NP_002737 | Full length | Activated by coexpression with MAP2K1 |
| ERK2 (MAPK1) | NP_620407 | Full length | Activated by coexpression with MAP2K1 |
| FAK1 (PTK2) | NP_722560 | Full length | Activated in vitro by SCR |
| FER | NP_005237.1 | Full length | No special measures |
| FGFR1 (activated) | NP_000595 | Partial length | Activated in vitro via autophosphorylation |
| FGFR1 (non-activated) | NP_000595 | Partial length | No special measures |
| FGFR2 | NP_075261 | Partial length | No special measures |
| FGFR3 (activated) | NP_000133 | Partial length | Activated in vitro via autophosphorylation |
| FGFR3 (non-activated) | NP_000133 | Partial length | No special measures |
| FGFR4 | NP_002002 | Partial length | No special measures |
| FLT3 (activated) | NP_004110 | Partial length | Activated in vitro via autophosphorylation |
| FLT3 (non-activated) | NP_004110 | Partial length | No special measures |
| FLT3 D835Y | NP_004110 | Partial length | Activated in vitro via autophosphorylation |
| FLT3 F691L | NP_004110 | Partial length | No special measures |
| FLT3 ITD | NP_004110 | Partial length | No special measures |
| FLT4 (VEGFR3) | NP_891555.1 | Partial length | Activated in vitro via autophosphorylation |
| FYN B | NP_694592 | Full length | No special measures |
| GCN2 (EIF2AK4) | NP_001013725 | Full length | No special measures |
| GSK3b | NP_002084 | Full length | No special measures |
| HER2 (ERBB2) | NP_004439 | Partial length | No special measures |

| | | | |
|---|---|---|---|
| HER4 (ERBB4) | NP_005226 | Partial length | No special measures |
| HGK (MAP4K4) | NP_004825 | Partial length | No special measures |
| HPK1 (MAP4K1) | NP_009112.1 | Partial length | No special measures |
| INSR | NP_000199 | Partial length | Activated in vitro via autophosphorylation |
| IRAK1 | NP_001560 | Full length | No special measures |
| IRAK3 | AAH57800.1 | Full length | No special measures |
| IRAK4 | NP_057207 | Full length | No special measures |
| ITK | NP_005537.3 | Full length | Activated in vitro via autophosphorylation |
| JAK1 | NP_002218.2 | Partial length | No special measures |
| JAK2 | NP_004963 | Partial length | Activated in vitro via autophosphorylation |
| JAK3 | NP_000206 | Partial length | No special measures |
| KDR (VEGFR2) | NP_002244 | Partial length | Activated in vitro via autophosphorylation |
| KHS1 (MAP4K5) | NP_942089 | Full length | No special measures |
| KIT (activated) | NP_000213 | Partial length | Activated in vitro via autophosphorylation |
| KIT (non-activated) | NP_000213 | Partial length | No special measures |
| KIT D816V | NP_000213 | Partial length | Activated in vitro via autophosphorylation |
| LATS1 | NP_004681.1 | Partial length | Activated by coexpression with MOBKL1A |
| LATS2 | NP_055387.2 | Partial length | No special measures |
| LCK | NP_005347 | Full length | No special measures |
| LIMK1 | NP_002305 | Partial length | Activated by coexpression with ROCK2 |
| LOK (STK10) | NP_005981.3 | Full length | No special measures |
| LRRK2 | NP_940980.3 | Full length | No special measures |
| LRRK2 G2019S | NP_940980.3 | Full length | No special measures |
| LYNA | NP_002341.1 | Full length | No special measures |
| MEK1 (MAP2K1) | NP_002746 | Full length | Activated in vitro by RAF1 |
| MEK2 (MAP2K2) | NP_109587 | Full length | Activated by coexpression with RAF1 |
| MEK7 (MAP2K7) | NP_00660186.1 | Full length | Activated by coexpression with MAP3K3 |
| MER (MERTK) | NP_006334.2 | Partial length | No special measures |
| MET (cMET) | NP_000236.2 | Partial length | No special measures |
| MINK1 | NP_056531 | Partial length | No special measures |
| MNK1 (MKNK1) | NP_945324 | Full length | Activated in vitro by MAPK12 |
| MNK2 (MKNK2) | NP_060042.2 | Full length | Activated in vitro by MAPK12 |
| MTOR | NP_004949.1 | Partial length | No special measures |
| MYLK2 (SKMLCK) | NP_149109 | Full length | No special measures |
| NIK (MAP3K14) | NP_003945.2 | Partial length | Constitutive kinase activity |
| p38a (MAPK14) | NP_620581 | Full length | Activated in vitro by MAP2K6 |
| PDGFRa | NP_006197 | Partial length | Activated in vitro via autophosphorylation |
| PDGFRb (activated) | NP_002600 | Partial length | Activated in vitro via autophosphorylation |
| PDGFRb (non-activated) | NP_002600 | Partial length | No special measures |
| PDK1 | NP_002604 | Full length | No special measures |
| PERK (EIF2AK3) | NP_004827.3 | Partial length | No special measures |
| PI3Ka/P85a (PIK3CA/PIK3R1) | NP_006209.2/NP_852664.1 | Full length | Activated by coexpression with P85a |
| PI3Ka/P85a (PIK3CA/PIK3R1) H1047R | NP_006209.2/NP_852664.1 | Full length | Activated by coexpression with P85a |
| PI3Kb/P85a (PIK3CB/PIK3R1) | NP_006210/NP_852664.1 | Full length | Activated by coexpression with P85a |
| PI3Kd/P85a (PIK3CD/PIK3R1) | NP_005017.2/NP_852664.1 | Full length | Activated by coexpression with P85a |
| PLK1 | NP_005021.2 | Full length | No special measures |
| PLK4 | NP_055079.3 | Partial length | Activated in vitro |
| RET | NP_066124 | Partial length | Activated in vitro via autophosphorylation |
| RIPK2 | NP_003812 | Partial length | No special measures |
| ROCK1 | NP_005397 | Partial length | No special measures |
| ROCK2 | NP_004841 | Partial length | No special measures |
| RON (MST1R) | NP_002438 | Partial length | Activated in vitro via autophosphorylation |
| ROS1 | NP_002935 | Partial length | No special measures |
| SGK1 (SGK) | NP_005618 | Partial length | Constituitive kinase activity |
| SIK1 (SNF1LK) | NP_775490.2 | Full length | No special measures |
| SIK2 (QIK) | NP_056006.1 | Full length | Activated in vitro |
| SIK3 (QSK) | P_079440.3 | Partial length | Constitutive kinase activity |
| SRC | NP_005408 | Full length | No special measures |
| STK3 (MST2) | NP_006272.2 | Full length | No special measures |
| STK4 (MST1) | NP_006273 | Full length | No special measures |
| SYK | NP_003168 | Full length | Activated in vitro via autophosphorylation |
| TAK1-TAB1 (MAP3K7-MAP3K7IP1) | NP_663304 | Partial length | Activated in vitro via autophosphorylation |
| TLK2 | NP_006843.2 | Partial length | No special measures |
| TNIK | NP_055843.1 | Partial length | No special measures |
| TRKA (NTRK1) | NP_002520 | Partial length | No special measures |
| TRKB (NTRK2) | NP_006171 | Partial length | Activated in vitro via autophosphorylation |
| TRKC (NTRK3) | NP_002521.2 | Partial length | Activated in vitro via autophosphorylation |
| TYK2 | NP_003322.2 | Partial length | No special measures |
| ULK1 | NP_003556.1 | Partial length | No special measures |
| ULK2 | NP_055498.3 | Partial length | Activated in vitro |
| YSK1 (STK25) | NP_006365 | Full length | Activated in vitro via autophosphorylation |
| ZAP70 | NP_001070 | Full length | Activated in vitro via autophosphorylation |

**Table S3.** Table of TR-FRET (KINETICfinder) binding kinetic parameters for danusertib against a panel of kinases

| Compound ID | Target | $k_{on}$ ($M^{-1}$ $s^{-1}$) | $k_{off}$ ($s^{-1}$) | τ (min) | $K_d$ (M) |
|---|---|---|---|---|---|
| Danusertib (PHA-739358) | RET | 6.98E+05 | 5.92E-04 | 28.13 | 8.49E-10 |
| Danusertib (PHA-739358) | AURKB (AURORA B)/INCENP | 4.62E+04 | 3.96E-05 | 420.94 | 8.58E-10 |
| Danusertib (PHA-739358) | ABL1 | 2.75E+06 | 2.51E-03 | 6.64 | 9.15E-10 |
| Danusertib (PHA-739358) | FGFR2 | 5.51E+05 | 6.44E-04 | 25.90 | 1.17E-09 |
| Danusertib (PHA-739358) | ABL2 (ARG) | 1.76E+06 | 2.60E-03 | 6.42 | 1.48E-09 |
| Danusertib (PHA-739358) | PLK4 | 6.97E+05 | 1.05E-03 | 15.93 | 1.50E-09 |
| Danusertib (PHA-739358) | HPK1 (MAP4K1) | 3.56E+06 | 6.75E-03 | 2.47 | 1.89E-09 |
| Danusertib (PHA-739358) | FGFR3 (activated) | 1.43E+06 | 3.81E-03 | 4.38 | 2.66E-09 |
| Danusertib (PHA-739358) | FGFR1 (activated) | 1.39E+06 | 4.37E-03 | 3.82 | 3.14E-09 |
| Danusertib (PHA-739358) | SRC | 1.24E+06 | 4.20E-03 | 3.96 | 3.38E-09 |
| Danusertib (PHA-739358) | AMPK (A1/B1/G1) | 1.74E+06 | 6.56E-03 | 2.54 | 3.78E-09 |
| Danusertib (PHA-739358) | FGFR1 (non-activated) | 6.52E+05 | 2.57E-03 | 6.49 | 3.94E-09 |
| Danusertib (PHA-739358) | AURKA (AURORA A) | 2.87E+05 | 1.15E-03 | 14.55 | 3.99E-09 |
| Danusertib (PHA-739358) | FAK1 (PTK2) | 8.74E+05 | 4.05E-03 | 4.11 | 4.63E-09 |
| Danusertib (PHA-739358) | FGFR3 (non-activated) | 5.52E+05 | 3.86E-03 | 4.31 | 7.00E-09 |
| Danusertib (PHA-739358) | JAK3 | 8.61E+05 | 6.12E-03 | 2.72 | 7.10E-09 |
| Danusertib (PHA-739358) | FLT4 (VEGFR3) | 2.54E+06 | 2.16E-02 | 0.77 | 8.52E-09 |
| Danusertib (PHA-739358) | AMPK (A2/B1/G1) | 1.20E+06 | 1.14E-02 | 1.46 | 9.47E-09 |
| Danusertib (PHA-739358) | CDK7/cyclin H/MNAT1 | 2.03E+06 | 1.94E-02 | 0.86 | 9.55E-09 |
| Danusertib (PHA-739358) | TRKB (NTRK2) | 1.23E+06 | 1.19E-02 | 1.40 | 9.67E-09 |
| Danusertib (PHA-739358) | BLK | 1.78E+06 | 2.06E-02 | 0.81 | 1.15E-08 |
| Danusertib (PHA-739358) | CSF1R (FMS) | 7.91E+05 | 9.60E-03 | 1.74 | 1.21E-08 |
| Danusertib (PHA-739358) | KIT D816V | 5.43E+05 | 6.87E-03 | 2.43 | 1.26E-08 |
| Danusertib (PHA-739358) | LIMK1 | 2.44E+06 | 3.11E-02 | 0.54 | 1.27E-08 |
| Danusertib (PHA-739358) | ROS1 | 3.67E+06 | 5.00E-02 | 0.33 | 1.36E-08 |
| Danusertib (PHA-739358) | DDR1 (activated) | 2.70E+05 | 3.76E-03 | 4.44 | 1.39E-08 |
| Danusertib (PHA-739358) | ALK | 2.22E+06 | 3.29E-02 | 0.51 | 1.48E-08 |
| Danusertib (PHA-739358) | FYN B | 6.66E+05 | 1.09E-02 | 1.52 | 1.64E-08 |
| Danusertib (PHA-739358) | DDR1 (non-activated) | 2.20E+04 | 4.97E-04 | 33.52 | 2.26E-08 |
| Danusertib (PHA-739358) | JAK2 | 6.95E+05 | 1.60E-02 | 1.00 | 2.31E-08 |
| Danusertib (PHA-739358) | ACK (TNK2) | 5.52E+05 | 1.36E-02 | 1.22 | 2.47E-08 |
| Danusertib (PHA-739358) | FLT3 D835Y | 1.03E+06 | 2.65E-02 | 0.63 | 2.58E-08 |
| Danusertib (PHA-739358) | SIK1 (SNF1LK) | 2.69E+05 | 7.59E-03 | 2.20 | 2.82E-08 |
| Danusertib (PHA-739358) | LCK | 5.88E+05 | 1.66E-02 | 1.00 | 2.82E-08 |
| Danusertib (PHA-739358) | LYNA | 5.28E+05 | 1.60E-02 | 1.04 | 3.04E-08 |
| Danusertib (PHA-739358) | KHS1 (MAP4K5) | 3.63E+05 | 1.35E-02 | 1.23 | 3.73E-08 |
| Danusertib (PHA-739358) | STK4 (MST1) | 3.40E+05 | 1.32E-02 | 1.26 | 3.90E-08 |
| Danusertib (PHA-739358) | TYK2 | 2.67E+05 | 1.05E-02 | 1.59 | 3.93E-08 |
| Danusertib (PHA-739358) | TRKA (NTRK1) | 4.59E+05 | 1.86E-02 | 0.90 | 4.05E-08 |
| Danusertib (PHA-739358) | KDR (VEGFR2) | 2.93E+05 | 1.21E-02 | 1.37 | 4.14E-08 |
| Danusertib (PHA-739358) | CDK2/cyclin A2 | 2.01E+05 | 1.02E-02 | 1.64 | 5.05E-08 |
| Danusertib (PHA-739358) | BTK (activated) | 3.80E+05 | 1.99E-02 | 0.84 | 5.25E-08 |
| Danusertib (PHA-739358) | LOK (STK10) | 1.21E+06 | 6.86E-02 | 0.24 | 5.66E-08 |
| Danusertib (PHA-739358) | TRKC (NTRK3) | 5.07E+05 | 2.98E-02 | 0.56 | 5.89E-08 |
| Danusertib (PHA-739358) | ITK | 7.59E+05 | 4.68E-02 | 0.36 | 6.17E-08 |
| Danusertib (PHA-739358) | KIT (activated) | 1.29E+05 | 8.01E-03 | 2.08 | 6.18E-08 |
| Danusertib (PHA-739358) | FER | 9.51E+05 | 8.10E-02 | 0.21 | 8.52E-08 |
| Danusertib (PHA-739358) | ULK2 | 1.82E+05 | 1.85E-02 | 0.90 | 1.02E-07 |
| Danusertib (PHA-739358) | FGFR4 | 3.09E+05 | 3.73E-02 | 0.45 | 1.21E-07 |
| Danusertib (PHA-739358) | YSK1 (STK25) | 4.86E+05 | 6.10E-02 | 0.27 | 1.25E-07 |
| Danusertib (PHA-739358) | BTK (non-activated) | 1.77E+05 | 2.22E-02 | 0.75 | 1.26E-07 |
| Danusertib (PHA-739358) | ALK5 (TGFBR1) | 2.69E+05 | 3.50E-02 | 0.48 | 1.30E-07 |
| Danusertib (PHA-739358) | FLT3 ITD | 1.44E+05 | 2.66E-02 | 0.63 | 1.84E-07 |
| Danusertib (PHA-739358) | ULK1 | 1.91E+05 | 3.63E-02 | 0.46 | 1.90E-07 |
| Danusertib (PHA-739358) | CDK2/cyclin E1 | 2.36E+04 | 4.78E-03 | 3.49 | 2.03E-07 |
| Danusertib (PHA-739358) | AXL | 7.68E+04 | 1.68E-02 | 0.99 | 2.19E-07 |
| Danusertib (PHA-739358) | CDK13/cyclin K | 5.35E+04 | 1.28E-02 | 1.31 | 2.39E-07 |
| Danusertib (PHA-739358) | SIK2 (QIK) | 6.90E+04 | 1.83E-02 | 0.91 | 2.65E-07 |
| Danusertib (PHA-739358) | FLT3 (activated) | 1.45E+05 | 4.15E-02 | 0.40 | 2.85E-07 |
| Danusertib (PHA-739358) | CLK2 | 8.40E+04 | 2.41E-02 | 0.69 | 2.86E-07 |
| Danusertib (PHA-739358) | ALK1 (ACVRL1) | 8.95E+04 | 3.09E-02 | 0.54 | 3.45E-07 |
| Danusertib (PHA-739358) | STK3 (MST2) | 7.38E+04 | 2.70E-02 | 0.62 | 3.65E-07 |
| Danusertib (PHA-739358) | MST1R (RON) | 6.91E+04 | 2.95E-02 | 0.56 | 4.27E-07 |
| Danusertib (PHA-739358) | EGFR (ERBB1) L858R | 1.56E+05 | 6.79E-02 | 0.25 | 4.36E-07 |
| Danusertib (PHA-739358) | TNIK | 4.57E+04 | 2.21E-02 | 0.80 | 4.82E-07 |

| | | | | | |
|---|---|---|---|---|---|
| Danusertib (PHA-739358) | CLK4 | 1.39E+05 | 6.81E-02 | 0.24 | 4.89E-07 |
| Danusertib (PHA-739358) | MYLK2 (SKMLCK) | 4.58E+04 | 2.33E-02 | 0.72 | 5.08E-07 |
| Danusertib (PHA-739358) | SYK | 6.92E+05 | 3.83E-01 | 0.04 | 5.53E-07 |
| Danusertib (PHA-739358) | CLK1 | 8.42E+05 | 4.69E-01 | 0.04 | 5.57E-07 |
| Danusertib (PHA-739358) | EGFR (ERBB1) E746_A750del | 2.07E+05 | 1.16E-01 | 0.14 | 5.61E-07 |
| Danusertib (PHA-739358) | RIPK2 | 8.42E+04 | 5.01E-02 | 0.33 | 5.96E-07 |
| Danusertib (PHA-739358) | EGFR (activated) | 4.10E+04 | 2.46E-02 | 0.68 | 6.00E-07 |
| Danusertib (PHA-739358) | PDGFRα | 6.10E+05 | 3.69E-01 | 0.05 | 6.05E-07 |
| Danusertib (PHA-739358) | EGFR (non-activated) | 4.40E+04 | 2.83E-02 | 0.59 | 6.43E-07 |
| Danusertib (PHA-739358) | IRAK3 | 4.92E+04 | 4.13E-02 | 0.40 | 8.38E-07 |
| Danusertib (PHA-739358) | JAK1 (JH1 JH2) | 5.22E+04 | 4.57E-02 | 0.36 | 8.76E-07 |
| Danusertib (PHA-739358) | CDK1/cyclin B1 | 4.01E+04 | 3.74E-02 | 0.45 | 9.32E-07 |
| Danusertib (PHA-739358) | CDK12/cyclin K | 1.69E+05 | 1.65E-01 | 0.10 | 9.74E-07 |
| Danusertib (PHA-739358) | CHEK2 (CHK2) | 2.84E+04 | 3.28E-02 | 0.51 | 1.15E-06 |
| Danusertib (PHA-739358) | TAK1/TAB1 (MAP3K7/MAP3K7IP1) | 1.06E+04 | 1.26E-02 | 1.33 | 1.19E-06 |
| Danusertib (PHA-739358) | SIK3 | 3.52E+05 | 4.23E-01 | 0.04 | 1.20E-06 |
| Danusertib (PHA-739358) | MINK1 | 5.55E+03 | 7.01E-03 | 2.38 | 1.26E-06 |
| Danusertib (PHA-739358) | BRAF V600E | 2.50E+04 | 3.30E-02 | 0.51 | 1.32E-06 |
| Danusertib (PHA-739358) | CAMK2a | 3.05E+03 | 4.74E-03 | 3.52 | 1.56E-06 |
| Danusertib (PHA-739358) | IRAK1 | 1.60E+04 | 2.68E-02 | 0.62 | 1.67E-06 |
| Danusertib (PHA-739358) | MER (MERTK) | 2.22E+04 | 3.84E-02 | 0.43 | 1.73E-06 |
| Danusertib (PHA-739358) | CDK9/cyclin T1 | 2.42E+04 | 4.29E-02 | 0.39 | 1.77E-06 |
| Danusertib (PHA-739358) | PLK1 | 1.07E+04 | 1.92E-02 | 0.87 | 1.79E-06 |
| Danusertib (PHA-739358) | ARAF | 7.95E+04 | 1.49E-01 | 0.11 | 1.88E-06 |
| Danusertib (PHA-739358) | INSR | 4.59E+03 | 8.68E-03 | 1.92 | 1.89E-06 |
| Danusertib (PHA-739358) | HGK (MAP4K4) | 8.42E+03 | 1.78E-02 | 0.94 | 2.11E-06 |
| Danusertib (PHA-739358) | PDGFRβ (activated) | 1.41E+05 | 3.29E-01 | 0.05 | 2.33E-06 |
| Danusertib (PHA-739358) | BRAF | 1.29E+04 | 3.06E-02 | 0.54 | 2.37E-06 |
| Danusertib (PHA-739358) | CAMK2d | 5.81E+03 | 1.49E-02 | 1.12 | 2.56E-06 |
| Danusertib (PHA-739358) | FLT3 F691L | 1.66E+04 | 4.32E-02 | 0.39 | 2.60E-06 |
| Danusertib (PHA-739358) | MET (cMET) | 9.34E+03 | 2.86E-02 | 0.58 | 3.06E-06 |
| Danusertib (PHA-739358) | GSK3B (GSK3 BETA) | 7.84E+03 | 2.59E-02 | 0.64 | 3.30E-06 |
| Danusertib (PHA-739358) | GCN2 (EIF2AK4) | 1.20E+04 | 4.48E-02 | 0.37 | 3.73E-06 |
| Danusertib (PHA-739358) | SGK1 (SGK) | 1.39E+04 | 5.77E-02 | 0.29 | 4.14E-06 |
| Danusertib (PHA-739358) | KIT (non-activated) | 9.38E+03 | 3.90E-02 | 0.43 | 4.16E-06 |
| Danusertib (PHA-739358) | HER4 (ERBB4) | 5.32E+03 | 2.38E-02 | 0.70 | 4.48E-06 |
| Danusertib (PHA-739358) | DYRK1B | 2.71E+03 | 1.33E-02 | 1.25 | 4.91E-06 |
| Danusertib (PHA-739358) | CDK16/cyclin Y | 7.36E+03 | 4.34E-02 | 0.38 | 5.90E-06 |
| Danusertib (PHA-739358) | ZAP70 | 6.33E+03 | 4.08E-02 | 0.41 | 6.45E-06 |
| Danusertib (PHA-739358) | LATS1 | >7.44E+04 | >5.00E-01 | <0.033 | 6.72E-06 |
| Danusertib (PHA-739358) | CHEK1 (CHK1) | >6.95E+04 | >5.00E-01 | <0.033 | 7.19E-06 |
| Danusertib (PHA-739358) | PDK1 | >6.33E+04 | >5.00E-01 | <0.033 | 7.90E-06 |
| Danusertib (PHA-739358) | NIK (MAP3K14) | 1.19E+04 | 9.49E-02 | 0.18 | 7.98E-06 |
| Danusertib (PHA-739358) | LATS2 | >3.12E+04 | >5.00E-01 | <0.033 | 1.60E-05 |
| Danusertib (PHA-739358) | ROCK1 | 4.06E+03 | 7.92E-02 | 0.21 | 1.95E-05 |
| Danusertib (PHA-739358) | p38a (MAPK14) | 1.34E+03 | 2.79E-02 | 0.60 | 2.08E-05 |
| Danusertib (PHA-739358) | FLT3 (non-activated) | 1.15E+03 | 2.40E-02 | 0.70 | 2.09E-05 |
| Danusertib (PHA-739358) | AKT1 | NE | NE | NE | >2.50E-05 |
| Danusertib (PHA-739358) | ASK1 (MAP3K5) | NE | NE | NE | >2.50E-05 |
| Danusertib (PHA-739358) | CDK8/cyclin C | NE | NE | NE | >2.50E-05 |
| Danusertib (PHA-739358) | CK1e (CSNK1E) | NE | NE | NE | >2.50E-05 |
| Danusertib (PHA-739358) | CRAF (RAF1) | NE | NE | NE | >2.50E-05 |
| Danusertib (PHA-739358) | ERK1 (MAPK3) | NE | NE | NE | >2.50E-05 |
| Danusertib (PHA-739358) | ERK2 (MAPK1) | NE | NE | NE | >2.50E-05 |
| Danusertib (PHA-739358) | HER2 (ERBB2) | NE | NE | NE | >2.50E-05 |
| Danusertib (PHA-739358) | IRAK4 | NE | NE | NE | >2.50E-05 |
| Danusertib (PHA-739358) | MEK1 (MAP2K1) | NE | NE | NE | >2.50E-05 |
| Danusertib (PHA-739358) | MEK2 (MAP2K2) | NE | NE | NE | >2.50E-05 |
| Danusertib (PHA-739358) | MEK7 (MAP2K7) | NE | NE | NE | >2.50E-05 |
| Danusertib (PHA-739358) | MNK1 (MKNK1) | NE | NE | NE | >2.50E-05 |
| Danusertib (PHA-739358) | MNK2 (MKNK2) | NE | NE | NE | >2.50E-05 |
| Danusertib (PHA-739358) | PDGFRβ (non-activated) | NE | NE | NE | >2.50E-05 |
| Danusertib (PHA-739358) | PERK (EIF2AK3) | NE | NE | NE | >2.50E-05 |
| Danusertib (PHA-739358) | PI3Ka/P85a (PIK3CA/PIK3R1) | NE | NE | NE | >2.50E-05 |
| Danusertib (PHA-739358) | PI3Ka/P85a (PIK3CA/PIK3R1) H1047R | NE | NE | NE | >2.50E-05 |
| Danusertib (PHA-739358) | PI3Kb/P85a (PIK3CB/PIK3R1) | NE | NE | NE | >2.50E-05 |
| Danusertib (PHA-739358) | PI3Kd/P85a (PIK3CD/PIK3R1) | NE | NE | NE | >2.50E-05 |
| Danusertib (PHA-739358) | ROCK2 | NE | NE | NE | >2.50E-05 |
| Danusertib (PHA-739358) | TLK2 | NE | NE | NE | >2.50E-05 |

**Table S4.** Table of TR-FRET (KINETICfinder) binding kinetic parameters for dasatinib against a panel of kinases

| Compound ID | Target | $k_{on}$ ($M^{-1}$ $s^{-1}$) | $k_{off}$ ($s^{-1}$) | τ (min) | $K_d$ (M) |
|---|---|---|---|---|---|
| Dasatinib Monohydrate (BMS-354825) | ABL1 | 7.46E+06 | 6.27E-05 | 265.86 | 8.40E-12 |
| Dasatinib Monohydrate (BMS-354825) | BLK | 7.33E+06 | 3.52E-04 | 47.37 | 4.80E-11 |
| Dasatinib Monohydrate (BMS-354825) | LYNA | 2.44E+06 | 1.46E-04 | 114.35 | 5.98E-11 |
| Dasatinib Monohydrate (BMS-354825) | ABL2 (ARG) | 4.75E+06 | 2.94E-04 | 56.76 | 6.18E-11 |
| Dasatinib Monohydrate (BMS-354825) | FYN B | 1.52E+06 | 1.04E-04 | 160.82 | 6.80E-11 |
| Dasatinib Monohydrate (BMS-354825) | SIK1 | 6.63E+06 | 5.22E-04 | 31.91 | 7.87E-11 |
| Dasatinib Monohydrate (BMS-354825) | SRC | 8.25E+06 | 7.43E-04 | 22.43 | 9.01E-11 |
| Dasatinib Monohydrate (BMS-354825) | KIT (activated) | 8.27E+05 | 9.84E-05 | 169.38 | 1.19E-10 |
| Dasatinib Monohydrate (BMS-354825) | LCK | 4.62E+06 | 5.98E-04 | 27.86 | 1.29E-10 |
| Dasatinib Monohydrate (BMS-354825) | CSF1R (FMS) | 3.42E+06 | 6.22E-04 | 26.81 | 1.82E-10 |
| Dasatinib Monohydrate (BMS-354825) | BTK (non-activated) | 5.62E+06 | 1.37E-03 | 12.20 | 2.43E-10 |
| Dasatinib Monohydrate (BMS-354825) | BTK (activated) | 4.27E+06 | 1.13E-03 | 14.72 | 2.65E-10 |
| Dasatinib Monohydrate (BMS-354825) | KIT D816V | 1.79E+06 | 7.05E-04 | 23.66 | 3.93E-10 |
| Dasatinib Monohydrate (BMS-354825) | SIK2 (QIK) | 1.30E+06 | 1.01E-03 | 16.49 | 7.79E-10 |
| Dasatinib Monohydrate (BMS-354825) | DDR1 (non-activated) | 7.37E+04 | 6.62E-05 | 251.65 | 8.98E-10 |
| Dasatinib Monohydrate (BMS-354825) | RIPK2 | 1.15E+07 | 1.51E-02 | 1.11 | 1.31E-09 |
| Dasatinib Monohydrate (BMS-354825) | ACK (TNK2) | 8.19E+05 | 1.35E-03 | 12.35 | 1.65E-09 |
| Dasatinib Monohydrate (BMS-354825) | DDR1 (activated) | 5.16E+05 | 8.87E-04 | 18.79 | 1.72E-09 |
| Dasatinib Monohydrate (BMS-354825) | PDGFRα | 2.89E+06 | 5.05E-03 | 3.30 | 1.75E-09 |
| Dasatinib Monohydrate (BMS-354825) | KIT (non-activated) | 7.15E+04 | 1.56E-04 | 106.92 | 2.18E-09 |
| Dasatinib Monohydrate (BMS-354825) | EGFR (ERBB1) E746 A750del | 6.24E+06 | 2.17E-02 | 0.77 | 3.47E-09 |
| Dasatinib Monohydrate (BMS-354825) | HER4 (ERBB4) | 2.31E+06 | 1.21E-02 | 1.38 | 5.23E-09 |
| Dasatinib Monohydrate (BMS-354825) | EGFR (ERBB1) L858R | 4.45E+06 | 2.42E-02 | 0.69 | 5.43E-09 |
| Dasatinib Monohydrate (BMS-354825) | EGFR (ERBB1) (non-activated) | 2.28E+06 | 1.85E-02 | 0.90 | 8.11E-09 |
| Dasatinib Monohydrate (BMS-354825) | SIK3 | 1.75E+06 | 1.70E-02 | 0.98 | 9.70E-09 |
| Dasatinib Monohydrate (BMS-354825) | EGFR (ERBB1) (activated) | 3.08E+06 | 3.07E-02 | 0.54 | 9.97E-09 |
| Dasatinib Monohydrate (BMS-354825) | ALK5 (TGFBR1) | 1.60E+06 | 2.18E-02 | 0.76 | 1.36E-08 |
| Dasatinib Monohydrate (BMS-354825) | BRAF V600E | 1.20E+06 | 2.37E-02 | 0.70 | 1.97E-08 |
| Dasatinib Monohydrate (BMS-354825) | KHS1 (MAP4K5) | 1.73E+06 | 3.58E-02 | 0.47 | 2.07E-08 |
| Dasatinib Monohydrate (BMS-354825) | ARAF | 1.02E+06 | 3.34E-02 | 0.50 | 3.29E-08 |
| Dasatinib Monohydrate (BMS-354825) | LIMK1 | 1.39E+06 | 5.28E-02 | 0.32 | 3.79E-08 |
| Dasatinib Monohydrate (BMS-354825) | BRAF | 8.73E+05 | 3.45E-02 | 0.48 | 3.95E-08 |
| Dasatinib Monohydrate (BMS-354825) | CRAF (RAF1) | 2.14E+06 | 1.10E-01 | 0.15 | 5.12E-08 |
| Dasatinib Monohydrate (BMS-354825) | p38a (MAPK14) | 2.41E+05 | 1.54E-02 | 1.08 | 6.40E-08 |
| Dasatinib Monohydrate (BMS-354825) | HER2 (ERBB2) | 5.89E+05 | 4.82E-02 | 0.35 | 8.18E-08 |
| Dasatinib Monohydrate (BMS-354825) | ALK1 (ACVRL1) | 7.08E+05 | 6.58E-02 | 0.25 | 9.29E-08 |
| Dasatinib Monohydrate (BMS-354825) | JAK1 (JH1 JH2) | 1.17E+05 | 1.90E-02 | 0.88 | 1.62E-07 |
| Dasatinib Monohydrate (BMS-354825) | RET | 2.22E+05 | 4.30E-02 | 0.39 | 1.94E-07 |
| Dasatinib Monohydrate (BMS-354825) | IRAK3 | 1.87E+05 | 3.87E-02 | 0.43 | 2.07E-07 |
| Dasatinib Monohydrate (BMS-354825) | HPK1 (MAP4K1) | 1.20E+05 | 3.18E-02 | 0.52 | 2.65E-07 |
| Dasatinib Monohydrate (BMS-354825) | SYK | 3.15E+05 | 9.88E-02 | 0.17 | 3.14E-07 |
| Dasatinib Monohydrate (BMS-354825) | JAK3 | 8.42E+04 | 2.72E-02 | 0.61 | 3.23E-07 |
| Dasatinib Monohydrate (BMS-354825) | JAK2 | 3.75E+05 | 1.21E-01 | 0.10 | 3.24E-07 |
| Dasatinib Monohydrate (BMS-354825) | FGFR2 | 1.23E+05 | 4.07E-02 | 0.41 | 3.30E-07 |
| Dasatinib Monohydrate (BMS-354825) | LOK (STK10) | 1.15E+05 | 3.93E-02 | 0.42 | 3.42E-07 |
| Dasatinib Monohydrate (BMS-354825) | KDR (VEGFR2) | 8.11E+04 | 6.57E-02 | 0.25 | 8.10E-07 |
| Dasatinib Monohydrate (BMS-354825) | MEK1 (MAP2K1) | 1.84E+04 | 1.70E-02 | 0.98 | 9.20E-07 |
| Dasatinib Monohydrate (BMS-354825) | MEK2 (MAP2K2) | 3.65E+04 | 3.51E-02 | 0.50 | 9.63E-07 |
| Dasatinib Monohydrate (BMS-354825) | FLT3 D835Y | 1.69E+04 | 1.84E-02 | 0.91 | 1.09E-06 |
| Dasatinib Monohydrate (BMS-354825) | FER | 2.31E+04 | 2.64E-02 | 0.63 | 1.14E-06 |
| Dasatinib Monohydrate (BMS-354825) | GCN2 (EIF2AK4) | 4.08E+04 | 5.09E-02 | 0.33 | 1.25E-06 |
| Dasatinib Monohydrate (BMS-354825) | FGFR1 (non-activated) | 5.29E+04 | 6.66E-02 | 0.25 | 1.26E-06 |
| Dasatinib Monohydrate (BMS-354825) | TNIK | 1.32E+04 | 1.66E-02 | 1.00 | 1.26E-06 |
| Dasatinib Monohydrate (BMS-354825) | CK1e (CSNK1E) | 2.92E+04 | 3.96E-02 | 0.42 | 1.36E-06 |
| Dasatinib Monohydrate (BMS-354825) | CLK4 | 8.83E+01 | 1.30E-04 | 128.30 | 1.47E-06 |
| Dasatinib Monohydrate (BMS-354825) | LRRK2 G2019S | 2.47E+04 | 3.76E-02 | 0.44 | 1.52E-06 |
| Dasatinib Monohydrate (BMS-354825) | LRRK2 | 1.25E+04 | 1.92E-02 | 0.87 | 1.53E-06 |
| Dasatinib Monohydrate (BMS-354825) | FGFR1 (activated) | 3.47E+04 | 5.38E-02 | 0.31 | 1.55E-06 |
| Dasatinib Monohydrate (BMS-354825) | MYLK2 (SKMLCK) | 2.13E+04 | 3.42E-02 | 0.49 | 1.61E-06 |
| Dasatinib Monohydrate (BMS-354825) | FGFR3 (activated) | 2.14E+05 | 3.63E-01 | 0.05 | 1.70E-06 |
| Dasatinib Monohydrate (BMS-354825) | FLT4 (VEGFR3) | 3.78E+04 | 8.40E-02 | 0.20 | 2.22E-06 |
| Dasatinib Monohydrate (BMS-354825) | TYK2 | >2.01E+05 | >5.00E-01 | <0.033 | 2.49E-06 |
| Dasatinib Monohydrate (BMS-354825) | YSK1 (STK25) | >1.71E+05 | >5.00E-01 | <0.033 | 2.92E-06 |
| Dasatinib Monohydrate (BMS-354825) | AURKB (AURORA B)/INCENP | 3.33E+04 | 1.00E-01 | 0.17 | 3.01E-06 |
| Dasatinib Monohydrate (BMS-354825) | IRAK4 | 1.22E+04 | 3.68E-02 | 0.45 | 3.03E-06 |
| Dasatinib Monohydrate (BMS-354825) | FGFR3 (non-activated) | 1.40E+04 | 4.45E-02 | 0.37 | 3.19E-06 |

| | | | | | |
|---|---|---|---|---|---|
| Dasatinib Monohydrate (BMS-354825) | STK4 (MST1) | 8.08E+03 | 2.89E-02 | 0.58 | 3.58E-06 |
| Dasatinib Monohydrate (BMS-354825) | PLK4 | >1.29E+05 | >5.00E-01 | <0.033 | 3.86E-06 |
| Dasatinib Monohydrate (BMS-354825) | FLT3 ITD | 1.83E+03 | 7.11E-03 | 2.34 | 3.90E-06 |
| Dasatinib Monohydrate (BMS-354825) | FLT3 (activated) | 1.96E+04 | 7.70E-02 | 0.22 | 3.93E-06 |
| Dasatinib Monohydrate (BMS-354825) | AURKA (AURORA A) | >1.27E+05 | >5.00E-01 | <0.033 | 3.93E-06 |
| Dasatinib Monohydrate (BMS-354825) | TAK1-TAB1 (MAP3K7-MAP3K7IP1) | 1.46E+03 | 6.11E-03 | 2.73 | 4.20E-06 |
| Dasatinib Monohydrate (BMS-354825) | TRKB (NTRK2) | >1.18E+05 | >5.00E-01 | <0.033 | 4.25E-06 |
| Dasatinib Monohydrate (BMS-354825) | MINK1 | 1.79E+03 | 7.85E-03 | 2.12 | 4.38E-06 |
| Dasatinib Monohydrate (BMS-354825) | AXL | 1.00E+04 | 4.58E-02 | 0.36 | 4.57E-06 |
| Dasatinib Monohydrate (BMS-354825) | MER (MERTK) | 9.92E+03 | 5.47E-02 | 0.30 | 5.51E-06 |
| Dasatinib Monohydrate (BMS-354825) | HGK (MAP4K4) | 1.27E+03 | 8.37E-03 | 1.99 | 6.59E-06 |
| Dasatinib Monohydrate (BMS-354825) | MET (cMET) | 4.98E+03 | 3.33E-02 | 0.50 | 6.69E-06 |
| Dasatinib Monohydrate (BMS-354825) | CLK1 | 1.32E+03 | 8.86E-03 | 1.88 | 6.73E-06 |
| Dasatinib Monohydrate (BMS-354825) | ZAP70 | 2.40E+03 | 1.67E-02 | 1.00 | 6.94E-06 |
| Dasatinib Monohydrate (BMS-354825) | CLK2 | 2.24E+03 | 1.94E-02 | 0.86 | 8.67E-06 |
| Dasatinib Monohydrate (BMS-354825) | TRKA (NTRK1) | 1.19E+04 | 1.09E-01 | 0.15 | 9.10E-06 |
| Dasatinib Monohydrate (BMS-354825) | AMPK (A2/B1/G1) | 4.34E+03 | 4.02E-02 | 0.41 | 9.26E-06 |
| Dasatinib Monohydrate (BMS-354825) | ITK | 4.19E+03 | 4.47E-02 | 0.37 | 1.07E-05 |
| Dasatinib Monohydrate (BMS-354825) | TRKC (NTRK3) | >4.48E+04 | >5.00E-01 | <0.033 | 1.12E-05 |
| Dasatinib Monohydrate (BMS-354825) | MNK2 (MKNK2) | 1.27E+03 | 1.43E-02 | 1.16 | 1.12E-05 |
| Dasatinib Monohydrate (BMS-354825) | FGFR4 | 1.96E+03 | 2.54E-02 | 0.66 | 1.30E-05 |
| Dasatinib Monohydrate (BMS-354825) | ROS1 | 5.50E+03 | 7.95E-02 | 0.21 | 1.45E-05 |
| Dasatinib Monohydrate (BMS-354825) | RON (MST1R) | 1.82E+03 | 2.66E-02 | 0.63 | 1.46E-05 |
| Dasatinib Monohydrate (BMS-354825) | PERK (EIF2AK3) | 3.48E+01 | 6.13E-04 | 27.18 | 1.76E-05 |
| Dasatinib Monohydrate (BMS-354825) | STK3 (MST2) | 1.11E+03 | 2.13E-02 | 0.78 | 1.92E-05 |
| Dasatinib Monohydrate (BMS-354825) | AMPK (A1/B1/G1) | >2.53E+04 | >5.00E-01 | <0.033 | 1.98E-05 |
| Dasatinib Monohydrate (BMS-354825) | IRAK1 | 1.32E+03 | 2.94E-02 | 0.57 | 2.23E-05 |
| Dasatinib Monohydrate (BMS-354825) | FLT3 (non-activated) | 6.58E+01 | 1.53E-03 | 10.91 | 2.32E-05 |
| Dasatinib Monohydrate (BMS-354825) | FAK1 (PTK2) | 1.86E+03 | 4.46E-02 | 0.37 | 2.40E-05 |
| Dasatinib Monohydrate (BMS-354825) | MNK1 (MKNK1) | 5.89E+02 | 1.42E-02 | 1.17 | 2.41E-05 |
| Dasatinib Monohydrate (BMS-354825) | CDK19/cyclin C | NE | NE | NE | >1.00E-06 |
| Dasatinib Monohydrate (BMS-354825) | AKT1 | NE | NE | NE | >2.50E-05 |
| Dasatinib Monohydrate (BMS-354825) | ALK | NE | NE | NE | >2.50E-05 |
| Dasatinib Monohydrate (BMS-354825) | ASK1 (MAP3K5) | NE | NE | NE | >2.50E-05 |
| Dasatinib Monohydrate (BMS-354825) | CAMK2a | NE | NE | NE | >2.50E-05 |
| Dasatinib Monohydrate (BMS-354825) | CAMK2d | NE | NE | NE | >2.50E-05 |
| Dasatinib Monohydrate (BMS-354825) | CDK1/cyclin B1 | NE | NE | NE | >2.50E-05 |
| Dasatinib Monohydrate (BMS-354825) | CDK12/cyclin K | NE | NE | NE | >2.50E-05 |
| Dasatinib Monohydrate (BMS-354825) | CDK13/cyclin K | NE | NE | NE | >2.50E-05 |
| Dasatinib Monohydrate (BMS-354825) | CDK16/cyclin Y | NE | NE | NE | >2.50E-05 |
| Dasatinib Monohydrate (BMS-354825) | CDK2/cyclin A2 | NE | NE | NE | >2.50E-05 |
| Dasatinib Monohydrate (BMS-354825) | CDK2/cyclin E1 | NE | NE | NE | >2.50E-05 |
| Dasatinib Monohydrate (BMS-354825) | CDK7/cyclin H/MNAT1 | NE | NE | NE | >2.50E-05 |
| Dasatinib Monohydrate (BMS-354825) | CDK8/cyclin C | NE | NE | NE | >2.50E-05 |
| Dasatinib Monohydrate (BMS-354825) | CDK9/cyclin T1 | NE | NE | NE | >2.50E-05 |
| Dasatinib Monohydrate (BMS-354825) | CHEK1 (CHK1) | NE | NE | NE | >2.50E-05 |
| Dasatinib Monohydrate (BMS-354825) | CHEK2 (CHK2) | NE | NE | NE | >2.50E-05 |
| Dasatinib Monohydrate (BMS-354825) | DYRK1A | NE | NE | NE | >2.50E-05 |
| Dasatinib Monohydrate (BMS-354825) | DYRK1B | NE | NE | NE | >2.50E-05 |
| Dasatinib Monohydrate (BMS-354825) | ERK1 (MAPK3) | NE | NE | NE | >2.50E-05 |
| Dasatinib Monohydrate (BMS-354825) | ERK2 (MAPK1) | NE | NE | NE | >2.50E-05 |
| Dasatinib Monohydrate (BMS-354825) | FLT3 F691L | NE | NE | NE | >2.50E-05 |
| Dasatinib Monohydrate (BMS-354825) | GSK3b (GSK3 BETA) | NE | NE | NE | >2.50E-05 |
| Dasatinib Monohydrate (BMS-354825) | INSR | NE | NE | NE | >2.50E-05 |
| Dasatinib Monohydrate (BMS-354825) | LATS1 | NE | NE | NE | >2.50E-05 |
| Dasatinib Monohydrate (BMS-354825) | LATS2 | NE | NE | NE | >2.50E-05 |
| Dasatinib Monohydrate (BMS-354825) | MTOR | NE | NE | NE | >2.50E-05 |
| Dasatinib Monohydrate (BMS-354825) | NIK (MAP3K14) | NE | NE | NE | >2.50E-05 |
| Dasatinib Monohydrate (BMS-354825) | PDK1 | NE | NE | NE | >2.50E-05 |
| Dasatinib Monohydrate (BMS-354825) | PI3Ka/P85a (PIK3CA/PIK3R1) | NE | NE | NE | >2.50E-05 |
| Dasatinib Monohydrate (BMS-354825) | PI3Ka/P85a (PIK3CA/PIK3R1) H1047R | NE | NE | NE | >2.50E-05 |
| Dasatinib Monohydrate (BMS-354825) | PI3Kb/P85a (PIK3CB/PIK3R1) | NE | NE | NE | >2.50E-05 |
| Dasatinib Monohydrate (BMS-354825) | PI3Kd/P85a (PIK3CD/PIK3R1) | NE | NE | NE | >2.50E-05 |
| Dasatinib Monohydrate (BMS-354825) | PLK1 | NE | NE | NE | >2.50E-05 |
| Dasatinib Monohydrate (BMS-354825) | ROCK1 | NE | NE | NE | >2.50E-05 |
| Dasatinib Monohydrate (BMS-354825) | ROCK2 | NE | NE | NE | >2.50E-05 |
| Dasatinib Monohydrate (BMS-354825) | SGK1 (SGK) | NE | NE | NE | >2.50E-05 |
| Dasatinib Monohydrate (BMS-354825) | TLK2 | NE | NE | NE | >2.50E-05 |
| Dasatinib Monohydrate (BMS-354825) | ULK1 | NE | NE | NE | >2.50E-05 |
| Dasatinib Monohydrate (BMS-354825) | ULK2 | NE | NE | NE | >2.50E-05 |

**Table S5.** Table of TR-FRET (KINETICfinder) binding kinetic parameters for encorafenib against a panel of kinases

| Compound ID | Target | $k_{on}$ ($M^{-1}$ $s^{-1}$) | $k_{off}$ ($s^{-1}$) | τ (min) | $K_d$ (M) |
|---|---|---|---|---|---|
| Encorafenib (LGX818) | BRAF | 4.01E+05 | 8.01E-05 | 208.01 | 2.00E-10 |
| Encorafenib (LGX818) | BRAF V600E | 2.85E+05 | 6.81E-05 | 244.82 | 2.39E-10 |
| Encorafenib (LGX818) | ARAF | 4.55E+05 | 1.09E-04 | 153.30 | 2.39E-10 |
| Encorafenib (LGX818) | CRAF (RAF1) | 5.39E+05 | 5.48E-04 | 30.42 | 1.02E-09 |
| Encorafenib (LGX818) | LIMK1 | 1.31E+06 | 1.67E-02 | 1.00 | 1.28E-08 |
| Encorafenib (LGX818) | RIPK2 | 1.41E+04 | 6.13E-04 | 27.19 | 4.34E-08 |
| Encorafenib (LGX818) | GCN2 (EIF2AK4) | 3.58E+04 | 2.18E-03 | 7.66 | 6.08E-08 |
| Encorafenib (LGX818) | IRAK1 | 7.50E+03 | 4.68E-04 | 35.64 | 6.23E-08 |
| Encorafenib (LGX818) | GSK3b (GSK3 BETA) | 5.53E+04 | 5.21E-03 | 3.20 | 9.42E-08 |
| Encorafenib (LGX818) | p38a (MAPK14) | 1.58E+05 | 2.03E-02 | 0.82 | 1.28E-07 |
| Encorafenib (LGX818) | TNIK | 3.35E+04 | 2.07E-02 | 0.80 | 6.20E-07 |
| Encorafenib (LGX818) | CLK4 | 1.47E+03 | 1.10E-03 | 15.22 | 7.47E-07 |
| Encorafenib (LGX818) | KHS1 (MAP4K5) | 5.33E+04 | 4.04E-02 | 0.41 | 7.58E-07 |
| Encorafenib (LGX818) | CK1e (CSNK1E) | 2.59E+04 | 1.99E-02 | 0.84 | 7.70E-07 |
| Encorafenib (LGX818) | HGK (MAP4K4) | 2.82E+03 | 2.82E-03 | 5.90 | 1.00E-06 |
| Encorafenib (LGX818) | CLK1 | 9.84E+02 | 9.90E-04 | 16.84 | 1.01E-06 |
| Encorafenib (LGX818) | TAK1/TAB1 (MAP3K7/MAP3K7IP1) | 2.01E+03 | 2.83E-03 | 5.89 | 1.40E-06 |
| Encorafenib (LGX818) | MNK2 (MKNK2) | 2.21E+04 | 3.24E-02 | 0.51 | 1.46E-06 |
| Encorafenib (LGX818) | MINK1 | 2.43E+03 | 3.60E-03 | 4.63 | 1.48E-06 |
| Encorafenib (LGX818) | DYRK1A | 5.70E+01 | 1.07E-04 | 156.30 | 1.87E-06 |
| Encorafenib (LGX818) | HPK1 (MAP4K1) | 6.81E+03 | 1.40E-02 | 1.19 | 2.05E-06 |
| Encorafenib (LGX818) | ERK2 (MAPK1) | 9.51E+03 | 2.65E-02 | 0.63 | 2.78E-06 |
| Encorafenib (LGX818) | FLT4 (VEGFR3) | 1.16E+04 | 3.77E-02 | 0.44 | 3.26E-06 |
| Encorafenib (LGX818) | CDK2/cyclin A2 | 2.13E+03 | 7.09E-03 | 2.35 | 3.32E-06 |
| Encorafenib (LGX818) | ALK5 (TGFBR1) | 8.09E+03 | 2.96E-02 | 0.56 | 3.66E-06 |
| Encorafenib (LGX818) | KDR (VEGFR2) | 7.11E+03 | 2.93E-02 | 0.57 | 4.13E-06 |
| Encorafenib (LGX818) | CDK16/cyclin Y | 2.67E+03 | 1.22E-02 | 1.37 | 4.56E-06 |
| Encorafenib (LGX818) | MNK1 (MKNK1) | 4.79E+03 | 2.41E-02 | 0.69 | 5.03E-06 |
| Encorafenib (LGX818) | ERK1 (MAPK3) | 2.45E+03 | 1.37E-02 | 1.22 | 5.60E-06 |
| Encorafenib (LGX818) | YSK1 (STK25) | 8.12E+03 | 5.21E-02 | 0.32 | 6.41E-06 |
| Encorafenib (LGX818) | CDK1/cyclin B1 | 2.49E+03 | 1.61E-02 | 1.03 | 6.47E-06 |
| Encorafenib (LGX818) | CDK2/cyclin E1 | 1.39E+03 | 9.03E-03 | 1.85 | 6.51E-06 |
| Encorafenib (LGX818) | MET (cMET) | 2.94E+03 | 2.21E-02 | 0.75 | 7.52E-06 |
| Encorafenib (LGX818) | HER2 (ERBB2) | 4.54E+03 | 3.42E-02 | 0.49 | 7.55E-06 |
| Encorafenib (LGX818) | DYRK1B | 9.08E+02 | 7.28E-03 | 2.29 | 8.01E-06 |
| Encorafenib (LGX818) | EGFR (ERBB1) E746_A750del | 1.74E+03 | 1.45E-02 | 1.15 | 8.31E-06 |
| Encorafenib (LGX818) | AMPK (A2/B1/G1) | >6.01E+04 | >5.00E-01 | <0.033 | 8.32E-06 |
| Encorafenib (LGX818) | CLK2 | 1.80E+02 | 1.79E-03 | 9.32 | 9.91E-06 |
| Encorafenib (LGX818) | HER4 (ERBB4) | 1.67E+03 | 1.68E-02 | 0.99 | 1.01E-05 |
| Encorafenib (LGX818) | ULK2 | 1.94E+04 | 1.96E-01 | 0.08 | 1.01E-05 |
| Encorafenib (LGX818) | CDK9/cyclin T1 | 2.81E+03 | 3.21E-02 | 0.52 | 1.14E-05 |
| Encorafenib (LGX818) | CDK8/cyclin C | 5.93E+01 | 6.90E-04 | 24.17 | 1.16E-05 |
| Encorafenib (LGX818) | IRAK3 | 9.60E+02 | 1.16E-02 | 1.43 | 1.21E-05 |
| Encorafenib (LGX818) | SGK1 (SGK) | 1.18E+03 | 1.46E-02 | 1.14 | 1.24E-05 |
| Encorafenib (LGX818) | AMPK (A1/B1/G1) | >3.75E+04 | >5.00E-01 | <0.033 | 1.33E-05 |
| Encorafenib (LGX818) | MYLK2 (SKMLCK) | 7.81E+02 | 1.07E-02 | 1.56 | 1.37E-05 |
| Encorafenib (LGX818) | MEK2 (MAP2K2) | 1.58E+03 | 2.21E-02 | 0.80 | 1.40E-05 |
| Encorafenib (LGX818) | CDK19/cyclin C | 9.25E+01 | 1.69E-03 | 9.87 | 1.83E-05 |
| Encorafenib (LGX818) | MER (MERTK) | 1.34E+04 | 2.56E-01 | 0.07 | 1.91E-05 |
| Encorafenib (LGX818) | CDK7/cyclin H/MNAT1 | 5.26E+02 | 1.03E-02 | 1.61 | 1.97E-05 |
| Encorafenib (LGX818) | EGFR (ERBB1) L858R | >2.43E+04 | >5.00E-01 | <0.033 | 2.06E-05 |
| Encorafenib (LGX818) | AKT1 | 2.49E+01 | 5.21E-04 | 31.99 | 2.09E-05 |
| Encorafenib (LGX818) | ULK1 | 4.82E+03 | 1.03E-01 | 0.16 | 2.14E-05 |
| Encorafenib (LGX818) | NIK (MAP3K14) | 1.41E+03 | 3.18E-02 | 0.52 | 2.26E-05 |
| Encorafenib (LGX818) | STK3 (MST2) | 6.11E+01 | 1.45E-03 | 11.51 | 2.37E-05 |
| Encorafenib (LGX818) | FGFR1 (non-activated) | 1.22E+03 | 2.93E-02 | 0.57 | 2.41E-05 |
| Encorafenib (LGX818) | MEK7 (MAP2K7) | 1.02E+02 | 2.47E-03 | 6.73 | 2.43E-05 |
| Encorafenib (LGX818) | MEK1 (MAP2K1) | 2.50E+02 | 6.19E-03 | 2.69 | 2.48E-05 |
| Encorafenib (LGX818) | CDK19/cyclin C | NE | NE | NE | >1.00E-06 |
| Encorafenib (LGX818) | CDK19/cyclin C | NE | NE | NE | >1.00E-07 |
| Encorafenib (LGX818) | ABL1 | NE | NE | NE | >2.50E-05 |
| Encorafenib (LGX818) | ABL2 (ARG) | NE | NE | NE | >2.50E-05 |
| Encorafenib (LGX818) | ACK (TNK2) | NE | NE | NE | >2.50E-05 |
| Encorafenib (LGX818) | ALK1 (ACVRL1) | NE | NE | NE | >2.50E-05 |
| Encorafenib (LGX818) | ASK1 (MAP3K5) | NE | NE | NE | >2.50E-05 |
| Encorafenib (LGX818) | AURKB (AURORA B)/INCENP | NE | NE | NE | >2.50E-05 |

| | | | | | |
|---|---|---|---|---|---|
| Encorafenib (LGX818) | AURORA A (AURKA) | NE | NE | NE | >2.50E-05 |
| Encorafenib (LGX818) | AXL | NE | NE | NE | >2.50E-05 |
| Encorafenib (LGX818) | BLK | NE | NE | NE | >2.50E-05 |
| Encorafenib (LGX818) | BTK (activated) | NE | NE | NE | >2.50E-05 |
| Encorafenib (LGX818) | CAMK2a | NE | NE | NE | >2.50E-05 |
| Encorafenib (LGX818) | CAMK2d | NE | NE | NE | >2.50E-05 |
| Encorafenib (LGX818) | CDK12/cyclin K | NE | NE | NE | >2.50E-05 |
| Encorafenib (LGX818) | CDK13/cyclin K | NE | NE | NE | >2.50E-05 |
| Encorafenib (LGX818) | CHEK1 (CHK1) | NE | NE | NE | >2.50E-05 |
| Encorafenib (LGX818) | CHEK2 (CHK2) | NE | NE | NE | >2.50E-05 |
| Encorafenib (LGX818) | CSF1R (FMS) | NE | NE | NE | >2.50E-05 |
| Encorafenib (LGX818) | DDR1 (activated) | NE | NE | NE | >2.50E-05 |
| Encorafenib (LGX818) | DDR1 (non-activated) | NE | NE | NE | >2.50E-05 |
| Encorafenib (LGX818) | EGFR (activated) | NE | NE | NE | >2.50E-05 |
| Encorafenib (LGX818) | EGFR (non-activated) | NE | NE | NE | >2.50E-05 |
| Encorafenib (LGX818) | FAK1 (PTK2) | NE | NE | NE | >2.50E-05 |
| Encorafenib (LGX818) | FER | NE | NE | NE | >2.50E-05 |
| Encorafenib (LGX818) | FGFR1 (activated) | NE | NE | NE | >2.50E-05 |
| Encorafenib (LGX818) | FGFR2 | NE | NE | NE | >2.50E-05 |
| Encorafenib (LGX818) | FGFR3 (activated) | NE | NE | NE | >2.50E-05 |
| Encorafenib (LGX818) | FGFR3 (non-activated) | NE | NE | NE | >2.50E-05 |
| Encorafenib (LGX818) | FGFR4 | NE | NE | NE | >2.50E-05 |
| Encorafenib (LGX818) | FLT3 (activated) | NE | NE | NE | >2.50E-05 |
| Encorafenib (LGX818) | FLT3 (non-activated) | NE | NE | NE | >2.50E-05 |
| Encorafenib (LGX818) | FLT3 D835Y | NE | NE | NE | >2.50E-05 |
| Encorafenib (LGX818) | FLT3 F691L | NE | NE | NE | >2.50E-05 |
| Encorafenib (LGX818) | FLT3 ITD | NE | NE | NE | >2.50E-05 |
| Encorafenib (LGX818) | FYN B | NE | NE | NE | >2.50E-05 |
| Encorafenib (LGX818) | INSR | NE | NE | NE | >2.50E-05 |
| Encorafenib (LGX818) | IRAK4 | NE | NE | NE | >2.50E-05 |
| Encorafenib (LGX818) | ITK | NE | NE | NE | >2.50E-05 |
| Encorafenib (LGX818) | JAK1 (JH1 JH2) | NE | NE | NE | >2.50E-05 |
| Encorafenib (LGX818) | JAK2 | NE | NE | NE | >2.50E-05 |
| Encorafenib (LGX818) | JAK3 | NE | NE | NE | >2.50E-05 |
| Encorafenib (LGX818) | KIT (activated) | NE | NE | NE | >2.50E-05 |
| Encorafenib (LGX818) | KIT (non-activated) | NE | NE | NE | >2.50E-05 |
| Encorafenib (LGX818) | KIT D816V | NE | NE | NE | >2.50E-05 |
| Encorafenib (LGX818) | LATS1 | NE | NE | NE | >2.50E-05 |
| Encorafenib (LGX818) | LATS2 | NE | NE | NE | >2.50E-05 |
| Encorafenib (LGX818) | LOK (STK10) | NE | NE | NE | >2.50E-05 |
| Encorafenib (LGX818) | LYNA | NE | NE | NE | >2.50E-05 |
| Encorafenib (LGX818) | MST1R (RON) | NE | NE | NE | >2.50E-05 |
| Encorafenib (LGX818) | MTOR | NE | NE | NE | >2.50E-05 |
| Encorafenib (LGX818) | PDGFRα | NE | NE | NE | >2.50E-05 |
| Encorafenib (LGX818) | PDGFRβ (activated) | NE | NE | NE | >2.50E-05 |
| Encorafenib (LGX818) | PDGFRβ (non-activated) | NE | NE | NE | >2.50E-05 |
| Encorafenib (LGX818) | PDK1 | NE | NE | NE | >2.50E-05 |
| Encorafenib (LGX818) | PI3Ka/P85a (PIK3CA/PIK3R1) | NE | NE | NE | >2.50E-05 |
| Encorafenib (LGX818) | PI3Ka/P85a (PIK3CA/PIK3R1) H1047R | NE | NE | NE | >2.50E-05 |
| Encorafenib (LGX818) | PI3Kb/P85a (PIK3CB/PIK3R1) | NE | NE | NE | >2.50E-05 |
| Encorafenib (LGX818) | PI3Kd/P85a (PIK3CD/PIK3R1) | NE | NE | NE | >2.50E-05 |
| Encorafenib (LGX818) | PLK1 | NE | NE | NE | >2.50E-05 |
| Encorafenib (LGX818) | PLK4 | NE | NE | NE | >2.50E-05 |
| Encorafenib (LGX818) | RET | NE | NE | NE | >2.50E-05 |
| Encorafenib (LGX818) | ROCK1 | NE | NE | NE | >2.50E-05 |
| Encorafenib (LGX818) | ROCK2 | NE | NE | NE | >2.50E-05 |
| Encorafenib (LGX818) | ROS1 | NE | NE | NE | >2.50E-05 |
| Encorafenib (LGX818) | SIK1 (SNF1LK) | NE | NE | NE | >2.50E-05 |
| Encorafenib (LGX818) | SIK2 (QIK) | NE | NE | NE | >2.50E-05 |
| Encorafenib (LGX818) | SIK3 | NE | NE | NE | >2.50E-05 |
| Encorafenib (LGX818) | SRC | NE | NE | NE | >2.50E-05 |
| Encorafenib (LGX818) | STK4 (MST1) | NE | NE | NE | >2.50E-05 |
| Encorafenib (LGX818) | SYK | NE | NE | NE | >2.50E-05 |
| Encorafenib (LGX818) | TLK2 | NE | NE | NE | >2.50E-05 |
| Encorafenib (LGX818) | TRKA (NTRK1) | NE | NE | NE | >2.50E-05 |
| Encorafenib (LGX818) | TRKB (NTRK2) | NE | NE | NE | >2.50E-05 |
| Encorafenib (LGX818) | TRKC (NTRK3) | NE | NE | NE | >2.50E-05 |
| Encorafenib (LGX818) | TYK2 | NE | NE | NE | >2.50E-05 |
| Encorafenib (LGX818) | ZAP70 | NE | NE | NE | >2.50E-05 |

**Table S6.** Table of TR-FRET (KINETICfinder) binding kinetic parameters for foretinib against a panel of kinases

| Compound ID | Target | $k_{on}$ ($M^{-1}$ $s^{-1}$) | $k_{off}$ ($s^{-1}$) | τ (min) | $K_d$ (M) |
|---|---|---|---|---|---|
| Foretinib | MET (cMET) | 1.20E+04 | 1.15E-05 | 1451.61 | 9.55E-10 |
| Foretinib | KDR | 1.17E+05 | 1.14E-04 | 146.31 | 9.75E-10 |
| Foretinib | CSFR1 (FMS) | 2.40E+04 | 2.57E-05 | 648.41 | 1.07E-09 |
| Foretinib | FLT4 (VEGFR3) | 8.45E+04 | 1.62E-04 | 102.92 | 1.92E-09 |
| Foretinib | LCK | 1.07E+06 | 2.37E-03 | 7.02 | 2.22E-09 |
| Foretinib | FLT3 (activated) | 7.30E+04 | 1.69E-04 | 98.52 | 2.32E-09 |
| Foretinib | RON (MST1R) | 5.76E+04 | 1.82E-04 | 91.57 | 3.16E-09 |
| Foretinib | FYN B | 1.07E+06 | 5.06E-03 | 3.29 | 4.74E-09 |
| Foretinib | MER (MERTK) | 3.50E+05 | 1.75E-03 | 9.52 | 4.99E-09 |
| Foretinib | KIT (activated) | 3.55E+04 | 3.89E-04 | 42.84 | 1.10E-08 |
| Foretinib | BLK | 1.05E+05 | 1.17E-03 | 14.19 | 1.12E-08 |
| Foretinib | RIPK2 | 4.27E+04 | 5.19E-04 | 32.11 | 1.22E-08 |
| Foretinib | PDGFRβ (activated) | 9.33E+03 | 1.91E-04 | 87.47 | 2.04E-08 |
| Foretinib | SRC | 7.94E+05 | 2.19E-02 | 0.76 | 2.75E-08 |
| Foretinib | KHS1 (MAP4K5) | 4.03E+05 | 1.23E-02 | 1.36 | 3.04E-08 |
| Foretinib | LYNA | 1.37E+05 | 4.34E-03 | 3.84 | 3.16E-08 |
| Foretinib | MEK2 (MAP2K2) | 3.39E+04 | 1.07E-03 | 15.55 | 3.16E-08 |
| Foretinib | PLK4 | 3.00E+05 | 9.90E-03 | 1.68 | 3.30E-08 |
| Foretinib | FER | 2.57E+05 | 8.78E-03 | 1.90 | 3.41E-08 |
| Foretinib | MNK2 (MKNK2) | 5.14E+04 | 1.93E-03 | 8.65 | 3.75E-08 |
| Foretinib | BTK (activated) | 1.07E+06 | 4.81E-02 | 0.35 | 4.50E-08 |
| Foretinib | MEK1 (MAP2K1) | 2.24E+04 | 1.02E-03 | 16.29 | 4.57E-08 |
| Foretinib | ALK1 (ACVRL1) | 1.38E+05 | 9.90E-03 | 1.68 | 7.19E-08 |
| Foretinib | PERK (EIF2AK3) | 3.08E+05 | 2.36E-02 | 0.70 | 7.68E-08 |
| Foretinib | ABL1 | 6.17E+05 | 5.50E-02 | 0.30 | 8.91E-08 |
| Foretinib | CDK8/Cyc C | 3.80E+03 | 3.47E-04 | 48.07 | 9.12E-08 |
| Foretinib | CDK19/Cyc C | 1.77E+03 | 1.69E-04 | 98.85 | 9.50E-08 |
| Foretinib | TNIK | 1.78E+05 | 1.76E-02 | 0.95 | 9.88E-08 |
| Foretinib | BTK (non-activated) | 4.35E+05 | 4.60E-02 | 0.36 | 1.06E-07 |
| Foretinib | ROS1 | 1.60E+05 | 2.16E-02 | 0.77 | 1.35E-07 |
| Foretinib | HPK1 (MAP4K1) | 1.52E+05 | 2.22E-02 | 0.75 | 1.45E-07 |
| Foretinib | KIT (non-activated) | 2.04E+03 | 3.01E-04 | 55.43 | 1.48E-07 |
| Foretinib | FLT3 D835Y | 2.94E+04 | 5.90E-03 | 2.82 | 2.01E-07 |
| Foretinib | IRAK3 | 7.45E+05 | 2.17E-01 | 0.08 | 2.91E-07 |
| Foretinib | CLK4 | 1.95E+05 | 6.56E-02 | 0.25 | 3.37E-07 |
| Foretinib | EGFR (ERBB1) (non-activated) | 1.73E+05 | 5.91E-02 | 0.28 | 3.42E-07 |
| Foretinib | ACK (TNK2) | 4.06E+04 | 1.49E-02 | 1.12 | 3.67E-07 |
| Foretinib | MINK1 | 7.78E+04 | 3.36E-02 | 0.50 | 4.31E-07 |
| Foretinib | EGFR (ERBB1) (activated) | 5.10E+04 | 2.28E-02 | 0.73 | 4.46E-07 |
| Foretinib | FLT3 F691L | 8.92E+01 | 4.01E-05 | 416.02 | 4.49E-07 |
| Foretinib | ITK | 3.11E+05 | 1.47E-01 | 0.11 | 4.72E-07 |
| Foretinib | LOK (STK10) | 1.55E+04 | 7.84E-03 | 2.12 | 5.06E-07 |
| Foretinib | AURKA (AURORA A) | 7.91E+05 | 4.08E-01 | 0.04 | 5.15E-07 |
| Foretinib | CLK1 | 7.34E+05 | 4.02E-01 | 0.04 | 5.48E-07 |
| Foretinib | HER2 (ERBB2) | 3.95E+03 | 2.18E-03 | 7.64 | 5.53E-07 |
| Foretinib | MNK1 (MKNK1) | 3.00E+04 | 1.85E-02 | 0.90 | 6.15E-07 |
| Foretinib | ABL2 (ARG) | >7.51E+05 | >5.00E-01 | <0.033 | 6.65E-07 |
| Foretinib | CHEK2 (CHK2) | 5.47E+04 | 3.74E-02 | 0.45 | 6.84E-07 |
| Foretinib | FGFR2 | 3.65E+05 | 2.53E-01 | 0.07 | 6.94E-07 |
| Foretinib | INSR | 2.70E+04 | 1.90E-02 | 0.88 | 7.03E-07 |
| Foretinib | FAK1 (PTK2) | 5.92E+04 | 4.26E-02 | 0.39 | 7.20E-07 |
| Foretinib | HER4 (ERBB4) | 4.75E+05 | 3.63E-01 | 0.05 | 7.65E-07 |
| Foretinib | KIT D816V | 6.45E+04 | 5.12E-02 | 0.33 | 7.93E-07 |
| Foretinib | LIMK1 | >5.76E+05 | >5.00E-01 | <0.033 | 8.69E-07 |
| Foretinib | EGFR (ERBB1) E746_A750del | 2.54E+04 | 2.33E-02 | 0.72 | 9.17E-07 |
| Foretinib | p38a (MAPK14) | 4.04E+04 | 3.75E-02 | 0.45 | 9.26E-07 |
| Foretinib | FGFR3 (activated) | >4.77E+05 | >5.00E-01 | <0.033 | 1.05E-06 |
| Foretinib | CDK13/cyclin K | 1.83E+04 | 1.92E-02 | 0.87 | 1.05E-06 |
| Foretinib | FGFR1 (activated) | >4.36E+05 | >5.00E-01 | <0.033 | 1.15E-06 |
| Foretinib | HGK (MAP4K4) | >4.18E+05 | >5.00E-01 | <0.033 | 1.20E-06 |
| Foretinib | ALK | 2.80E+04 | 5.97E-02 | 0.28 | 2.13E-06 |
| Foretinib | FGFR1 (non-activated) | >2.26E+05 | >5.00E-01 | <0.033 | 2.21E-06 |
| Foretinib | TAK1-TAB1 (MAP3K7-MAP3K7IP1) | 6.56E+03 | 1.47E-02 | 1.14 | 2.24E-06 |
| Foretinib | JAK2 | 2.13E+04 | 5.31E-02 | 0.30 | 2.49E-06 |
| Foretinib | ALK5 (TGFBR1) | 2.86E+04 | 8.27E-02 | 0.20 | 2.89E-06 |

| | | | | | |
|---|---|---|---|---|---|
| Foretinib | CDK16/cyclin Y | 8.31E+03 | 4.46E-02 | 0.37 | 5.37E-06 |
| Foretinib | EGFR (ERBB1) L858R | 1.94E+04 | 1.21E-01 | 0.14 | 6.27E-06 |
| Foretinib | AMPK (A2/B1/G1) | 6.92E+03 | 6.60E-02 | 0.25 | 9.54E-06 |
| Foretinib | CK1e (CSNK1E) | 3.61E+03 | 4.37E-02 | 0.38 | 1.21E-05 |
| Foretinib | IRAK4 | 1.06E+03 | 2.53E-02 | 0.66 | 2.39E-05 |
| Foretinib | CRAF (RAF1) | NE | NE | NE | >1.00E-05 |
| Foretinib | AKT1 | NE | NE | NE | >2.50E-05 |
| Foretinib | ARAF | NE | NE | NE | >2.50E-05 |
| Foretinib | ASK1 (MAP3K5) | NE | NE | NE | >2.50E-05 |
| Foretinib | BRAF | NE | NE | NE | >2.50E-05 |
| Foretinib | CAMK2a | NE | NE | NE | >2.50E-05 |
| Foretinib | CAMK2d | NE | NE | NE | >2.50E-05 |
| Foretinib | CDK1/cyclin B1 | NE | NE | NE | >2.50E-05 |
| Foretinib | CDK2/cyclin A2 | NE | NE | NE | >2.50E-05 |
| Foretinib | CDK2/cyclin E1 | NE | NE | NE | >2.50E-05 |
| Foretinib | CDK7/cyclin H/MNAT1 | NE | NE | NE | >2.50E-05 |
| Foretinib | CDK9/cyclin T1 | NE | NE | NE | >2.50E-05 |
| Foretinib | CHEK1 (CHK1) | NE | NE | NE | >2.50E-05 |
| Foretinib | CLK2 | NE | NE | NE | >2.50E-05 |
| Foretinib | DYRK1A | NE | NE | NE | >2.50E-05 |
| Foretinib | DYRK1B | NE | NE | NE | >2.50E-05 |
| Foretinib | ERK1 (MAPK3) | NE | NE | NE | >2.50E-05 |
| Foretinib | ERK2 (MAPK1) | NE | NE | NE | >2.50E-05 |
| Foretinib | FGFR4 | NE | NE | NE | >2.50E-05 |
| Foretinib | GSK3b (GSK3 BETA) | NE | NE | NE | >2.50E-05 |
| Foretinib | IRAK1 | NE | NE | NE | >2.50E-05 |
| Foretinib | JAK1 (JH1 JH2) | NE | NE | NE | >2.50E-05 |
| Foretinib | JAK3 | NE | NE | NE | >2.50E-05 |
| Foretinib | LATS1 | NE | NE | NE | >2.50E-05 |
| Foretinib | LATS2 | NE | NE | NE | >2.50E-05 |
| Foretinib | MEK7 (MAP2K7) | NE | NE | NE | >2.50E-05 |
| Foretinib | MYLK2 (SKMLCK) | NE | NE | NE | >2.50E-05 |
| Foretinib | NIK (MAP3K14) | NE | NE | NE | >2.50E-05 |
| Foretinib | PDK1 | NE | NE | NE | >2.50E-05 |
| Foretinib | PI3Ka/P85a (PIK3CA/PIK3R1) | NE | NE | NE | >2.50E-05 |
| Foretinib | PI3Ka/P85a (PIK3CA/PIK3R1) H1047R | NE | NE | NE | >2.50E-05 |
| Foretinib | PI3Kb/P85a (PIK3CB/PIK3R1) | NE | NE | NE | >2.50E-05 |
| Foretinib | PI3Kd/P85a (PIK3CD/PIK3R1) | NE | NE | NE | >2.50E-05 |
| Foretinib | PLK1 | NE | NE | NE | >2.50E-05 |
| Foretinib | ROCK1 | NE | NE | NE | >2.50E-05 |
| Foretinib | ROCK2 | NE | NE | NE | >2.50E-05 |
| Foretinib | SGK1 (SGK) | NE | NE | NE | >2.50E-05 |
| Foretinib | SIK1 (SNF1LK) | NE | NE | NE | >2.50E-05 |
| Foretinib | SIK2 (QIK) | NE | NE | NE | >2.50E-05 |
| Foretinib | SIK3 | NE | NE | NE | >2.50E-05 |
| Foretinib | STK3 (MST2) | NE | NE | NE | >2.50E-05 |
| Foretinib | TLK2 | NE | NE | NE | >2.50E-05 |
| Foretinib | TYK2 | NE | NE | NE | >2.50E-05 |
| Foretinib | ULK1 | NE | NE | NE | >2.50E-05 |
| Foretinib | ULK2 | NE | NE | NE | >2.50E-05 |
| Foretinib | YSK1 (STK25) | NE | NE | NE | >2.50E-05 |
| Foretinib | ZAP70 | NE | NE | NE | >2.50E-05 |
| Foretinib | AMPK (A1/B1/G1) | NE | NE | NE | >2.50E-06 |
| Foretinib | CDK12/cyclin K | NE | NE | NE | >2.50E-06 |
| Foretinib | FGFR3 (non-activated) | NE | NE | NE | >2.50E-06 |
| Foretinib | GCN2 (EIF2AK4) | NE | NE | NE | >2.50E-06 |
| Foretinib | STK4 (MST1) | NE | NE | NE | >2.50E-06 |
| Foretinib | SYK | NE | NE | NE | >2.50E-06 |

**Table S7.** Table of TR-FRET (KINETICfinder) binding kinetic parameters for tivozanib against a panel of kinases

| Compound ID | Target | $k_{on}$ ($M^{-1}$ $s^{-1}$) | $k_{off}$ ($s^{-1}$) | τ (min) | $K_d$ (M) |
|---|---|---|---|---|---|
| Tivozanib (AV-951) | KDR (VEGFR2) | 2.40E+05 | 5.48E-04 | 30.40 | 2.29E-09 |
| Tivozanib (AV-951) | FLT4 (VEGFR3) | 1.05E+06 | 2.88E-03 | 5.80 | 2.74E-09 |
| Tivozanib (AV-951) | KIT (activated) | 5.95E+04 | 1.80E-04 | 92.75 | 3.02E-09 |
| Tivozanib (AV-951) | RET | 1.60E+06 | 4.89E-03 | 3.41 | 3.06E-09 |
| Tivozanib (AV-951) | PDGFRα | 5.46E+05 | 2.55E-03 | 6.54 | 4.67E-09 |
| Tivozanib (AV-951) | PDGFRβ (activated) | 1.02E+04 | 9.64E-05 | 172.82 | 9.48E-09 |
| Tivozanib (AV-951) | DDR1 (non-activated) | 3.12E+03 | 3.25E-05 | 513.53 | 1.04E-08 |
| Tivozanib (AV-951) | CSF1R (FMS) | 9.30E+04 | 1.21E-03 | 13.79 | 1.30E-08 |
| Tivozanib (AV-951) | KIT (non-activated) | 6.93E+02 | 1.92E-05 | 870.26 | 2.76E-08 |
| Tivozanib (AV-951) | DDR1 (activated) | 4.25E+04 | 1.27E-03 | 13.10 | 3.00E-08 |
| Tivozanib (AV-951) | LYNA | 4.59E+05 | 1.48E-02 | 1.13 | 3.22E-08 |
| Tivozanib (AV-951) | LOK (STK10) | 2.56E+06 | 9.12E-02 | 0.18 | 3.56E-08 |
| Tivozanib (AV-951) | ABL1 | 2.22E+06 | 8.73E-02 | 0.19 | 3.94E-08 |
| Tivozanib (AV-951) | CDK19/cyclin C | 2.60E+04 | 1.41E-03 | 11.81 | 5.44E-08 |
| Tivozanib (AV-951) | RIPK2 | 9.47E+05 | 7.04E-02 | 0.24 | 7.44E-08 |
| Tivozanib (AV-951) | BLK | 9.09E+05 | 8.90E-02 | 0.19 | 9.79E-08 |
| Tivozanib (AV-951) | AURKB (AURORA B)/INCENP | 2.69E+05 | 2.88E-02 | 0.58 | 1.07E-07 |
| Tivozanib (AV-951) | CDK8/cyclin C | 2.07E+04 | 2.54E-03 | 6.55 | 1.23E-07 |
| Tivozanib (AV-951) | FLT3 (activated) | 2.44E+05 | 3.53E-02 | 0.47 | 1.45E-07 |
| Tivozanib (AV-951) | ARAF | 1.08E+05 | 1.77E-02 | 0.94 | 1.64E-07 |
| Tivozanib (AV-951) | TRKB (NTRK2) | 1.76E+06 | 3.03E-01 | 0.05 | 1.72E-07 |
| Tivozanib (AV-951) | PDGFRβ (non-activated) | 1.76E+02 | 3.19E-05 | 521.84 | 1.82E-07 |
| Tivozanib (AV-951) | TRKC (NTRK3) | 3.06E+05 | 5.82E-02 | 0.29 | 1.90E-07 |
| Tivozanib (AV-951) | EGFR (ERBB1) (activated) | 1.42E+05 | 3.11E-02 | 0.54 | 2.19E-07 |
| Tivozanib (AV-951) | BRAF | 2.03E+04 | 4.79E-03 | 3.48 | 2.35E-07 |
| Tivozanib (AV-951) | FER | 2.56E+05 | 7.38E-02 | 0.23 | 2.88E-07 |
| Tivozanib (AV-951) | KHS1 (MAP4K5) | 1.13E+06 | 3.46E-01 | 0.05 | 3.07E-07 |
| Tivozanib (AV-951) | EGFR (ERBB1) E746 A750del | 6.58E+04 | 2.44E-02 | 0.68 | 3.70E-07 |
| Tivozanib (AV-951) | EGFR (ERBB1) L858R | 1.50E+05 | 5.67E-02 | 0.29 | 3.77E-07 |
| Tivozanib (AV-951) | AXL | 7.19E+04 | 3.00E-02 | 0.56 | 4.17E-07 |
| Tivozanib (AV-951) | ABL2 (ARG) | 1.13E+05 | 5.35E-02 | 0.31 | 4.72E-07 |
| Tivozanib (AV-951) | JAK1 (JH1 JH2) | 3.84E+04 | 1.94E-02 | 0.86 | 5.05E-07 |
| Tivozanib (AV-951) | TRKA (NTRK1) | 5.18E+05 | 2.86E-01 | 0.06 | 5.53E-07 |
| Tivozanib (AV-951) | MET (cMET) | 5.77E+04 | 3.26E-02 | 0.51 | 5.66E-07 |
| Tivozanib (AV-951) | EGFR (ERBB1) (non-activated) | 5.69E+05 | 3.25E-01 | 0.05 | 5.71E-07 |
| Tivozanib (AV-951) | FGFR1 (activated) | 7.01E+05 | 4.57E-01 | 0.04 | 6.51E-07 |
| Tivozanib (AV-951) | PLK4 | >7.27E+05 | >5.00E-01 | <0.033 | 6.87E-07 |
| Tivozanib (AV-951) | MEK2 (MAP2K2) | 2.41E+04 | 1.68E-02 | 1.00 | 6.97E-07 |
| Tivozanib (AV-951) | TNIK | 2.96E+04 | 2.12E-02 | 0.80 | 7.18E-07 |
| Tivozanib (AV-951) | FLT3 (non-activated) | 2.72E+03 | 2.01E-03 | 8.28 | 7.41E-07 |
| Tivozanib (AV-951) | MEK1 (MAP2K1) | 2.86E+04 | 2.39E-02 | 0.70 | 8.37E-07 |
| Tivozanib (AV-951) | HPK1 (MAP4K1) | 1.34E+05 | 1.14E-01 | 0.15 | 8.47E-07 |
| Tivozanib (AV-951) | CK1e (CSNK1E) | 1.80E+04 | 1.62E-02 | 1.03 | 8.99E-07 |
| Tivozanib (AV-951) | SRC | 3.24E+05 | 2.97E-01 | 0.06 | 9.17E-07 |
| Tivozanib (AV-951) | FGFR2 | >5.01E+05 | >5.00E-01 | <0.033 | 9.98E-07 |
| Tivozanib (AV-951) | ROS1 | >4.57E+05 | >5.00E-01 | <0.033 | 1.09E-06 |
| Tivozanib (AV-951) | HER2 (ERBB2) | 1.81E+04 | 2.02E-02 | 0.82 | 1.12E-06 |
| Tivozanib (AV-951) | AURKA (AURORA A) | >4.03E+05 | >5.00E-01 | <0.033 | 1.24E-06 |
| Tivozanib (AV-951) | FGFR3 (activated) | >3.88E+05 | >5.00E-01 | <0.033 | 1.29E-06 |
| Tivozanib (AV-951) | p38a (MAPK14) | 2.85E+04 | 4.09E-02 | 0.41 | 1.43E-06 |
| Tivozanib (AV-951) | MER (MERTK) | 2.11E+04 | 4.54E-02 | 0.37 | 2.15E-06 |
| Tivozanib (AV-951) | FGFR1 (non-activated) | >2.15E+05 | >5.00E-01 | <0.033 | 2.33E-06 |
| Tivozanib (AV-951) | ALK1 (ACVRL1) | 1.88E+04 | 5.40E-02 | 0.31 | 2.88E-06 |
| Tivozanib (AV-951) | FYN B | >1.61E+05 | >5.00E-01 | <0.033 | 3.11E-06 |
| Tivozanib (AV-951) | ACK (TNK2) | >1.55E+05 | >5.00E-01 | <0.033 | 3.23E-06 |
| Tivozanib (AV-951) | MST1R (RON) | 2.03E+04 | 6.67E-02 | 0.25 | 3.28E-06 |
| Tivozanib (AV-951) | FLT3 ITD | 5.28E+03 | 1.75E-02 | 0.95 | 3.31E-06 |
| Tivozanib (AV-951) | CDK13/cyclin K | >1.35E+05 | >5.00E-01 | <0.033 | 3.72E-06 |
| Tivozanib (AV-951) | BTK (activated) | 1.25E+05 | 4.81E-01 | 0.03 | 3.83E-06 |
| Tivozanib (AV-951) | LIMK1 | 4.21E+04 | 1.64E-01 | 0.10 | 3.91E-06 |
| Tivozanib (AV-951) | FLT3 D835Y | 7.79E+03 | 3.58E-02 | 0.47 | 4.60E-06 |
| Tivozanib (AV-951) | MINK1 | 9.88E+03 | 6.58E-02 | 0.25 | 6.66E-06 |
| Tivozanib (AV-951) | BTK (non-activated) | >5.65E+04 | >5.00E-01 | <0.033 | 8.85E-06 |
| Tivozanib (AV-951) | ALK5 (TGFBR1) | >5.54E+04 | >5.00E-01 | <0.033 | 9.03E-06 |
| Tivozanib (AV-951) | JAK3 | >4.58E+04 | >5.00E-01 | <0.033 | 1.09E-05 |
| Tivozanib (AV-951) | AMPK (A1/B1/G1) | >4.40E+04 | >5.00E-01 | <0.033 | 1.14E-05 |

| | | | | | |
|---|---|---|---|---|---|
| Tivozanib (AV-951) | HGK (MAP4K4) | >4.30E+04 | >5.00E-01 | <0.033 | 1.16E-05 |
| Tivozanib (AV-951) | CLK1 | >4.29E+04 | >5.00E-01 | <0.033 | 1.17E-05 |
| Tivozanib (AV-951) | STK3 (MST2) | 5.92E+03 | 7.99E-02 | 0.21 | 1.35E-05 |
| Tivozanib (AV-951) | CHEK2 (CHK2) | 2.48E+03 | 3.77E-02 | 0.44 | 1.52E-05 |
| Tivozanib (AV-951) | IRAK3 | 3.47E+03 | 6.27E-02 | 0.27 | 1.81E-05 |
| Tivozanib (AV-951) | ROCK2 | >2.14E+04 | >5.00E-01 | <0.033 | 2.34E-05 |
| Tivozanib (AV-951) | AKT1 | NE | NE | NE | >2.50E-05 |
| Tivozanib (AV-951) | ALK | NE | NE | NE | >2.50E-05 |
| Tivozanib (AV-951) | AMPK (A2/B1/G1) | NE | NE | NE | >2.50E-05 |
| Tivozanib (AV-951) | ASK1 (MAP3K5) | NE | NE | NE | >2.50E-05 |
| Tivozanib (AV-951) | CAMK2a | NE | NE | NE | >2.50E-05 |
| Tivozanib (AV-951) | CAMK2d | NE | NE | NE | >2.50E-05 |
| Tivozanib (AV-951) | CDK1/cyclin B1 | NE | NE | NE | >2.50E-05 |
| Tivozanib (AV-951) | CDK12/cyclin K | NE | NE | NE | >2.50E-05 |
| Tivozanib (AV-951) | CDK16/cyclin Y | NE | NE | NE | >2.50E-05 |
| Tivozanib (AV-951) | CDK2/cyclin A2 | NE | NE | NE | >2.50E-05 |
| Tivozanib (AV-951) | CDK2/cyclin E1 | NE | NE | NE | >2.50E-05 |
| Tivozanib (AV-951) | CDK7/cyclin H/MNAT1 | NE | NE | NE | >2.50E-05 |
| Tivozanib (AV-951) | CDK9/cyclin T1 | NE | NE | NE | >2.50E-05 |
| Tivozanib (AV-951) | CHEK1 (CHK1) | NE | NE | NE | >2.50E-05 |
| Tivozanib (AV-951) | CLK2 | NE | NE | NE | >2.50E-05 |
| Tivozanib (AV-951) | CLK4 | NE | NE | NE | >2.50E-05 |
| Tivozanib (AV-951) | CRAF (RAF1) | NE | NE | NE | >2.50E-05 |
| Tivozanib (AV-951) | DYRK1A | NE | NE | NE | >2.50E-05 |
| Tivozanib (AV-951) | DYRK1B | NE | NE | NE | >2.50E-05 |
| Tivozanib (AV-951) | ERK1 (MAPK3) | NE | NE | NE | >2.50E-05 |
| Tivozanib (AV-951) | ERK2 (MAPK1) | NE | NE | NE | >2.50E-05 |
| Tivozanib (AV-951) | FAK1 (PTK2) | NE | NE | NE | >2.50E-05 |
| Tivozanib (AV-951) | FGFR3 (non-activated) | NE | NE | NE | >2.50E-05 |
| Tivozanib (AV-951) | GCN2 (EIF2AK4) | NE | NE | NE | >2.50E-05 |
| Tivozanib (AV-951) | GSK3b (GSK3 BETA) | NE | NE | NE | >2.50E-05 |
| Tivozanib (AV-951) | HER4 (ERBB4) | NE | NE | NE | >2.50E-05 |
| Tivozanib (AV-951) | INSR | NE | NE | NE | >2.50E-05 |
| Tivozanib (AV-951) | IRAK1 | NE | NE | NE | >2.50E-05 |
| Tivozanib (AV-951) | IRAK4 | NE | NE | NE | >2.50E-05 |
| Tivozanib (AV-951) | ITK | NE | NE | NE | >2.50E-05 |
| Tivozanib (AV-951) | JAK2 | NE | NE | NE | >2.50E-05 |
| Tivozanib (AV-951) | KIT D816V | NE | NE | NE | >2.50E-05 |
| Tivozanib (AV-951) | LATS1 | NE | NE | NE | >2.50E-05 |
| Tivozanib (AV-951) | LATS2 | NE | NE | NE | >2.50E-05 |
| Tivozanib (AV-951) | MEK7 (MAP2K7) | NE | NE | NE | >2.50E-05 |
| Tivozanib (AV-951) | MNK1 (MKNK1) | NE | NE | NE | >2.50E-05 |
| Tivozanib (AV-951) | MNK2 (MKNK2) | NE | NE | NE | >2.50E-05 |
| Tivozanib (AV-951) | MYLK2 (SKMLCK) | NE | NE | NE | >2.50E-05 |
| Tivozanib (AV-951) | NIK (MAP3K14) | NE | NE | NE | >2.50E-05 |
| Tivozanib (AV-951) | PDK1 | NE | NE | NE | >2.50E-05 |
| Tivozanib (AV-951) | PI3Ka/P85a (PIK3CA/PIK3R1) | NE | NE | NE | >2.50E-05 |
| Tivozanib (AV-951) | PI3Ka/P85a (PIK3CA/PIK3R1) H1047R | NE | NE | NE | >2.50E-05 |
| Tivozanib (AV-951) | PI3Kb/P85a (PIK3CB/PIK3R1) | NE | NE | NE | >2.50E-05 |
| Tivozanib (AV-951) | PI3Kd/P85a (PIK3CD/PIK3R1) | NE | NE | NE | >2.50E-05 |
| Tivozanib (AV-951) | PLK1 | NE | NE | NE | >2.50E-05 |
| Tivozanib (AV-951) | ROCK1 | NE | NE | NE | >2.50E-05 |
| Tivozanib (AV-951) | SGK1 (SGK) | NE | NE | NE | >2.50E-05 |
| Tivozanib (AV-951) | SIK1 | NE | NE | NE | >2.50E-05 |
| Tivozanib (AV-951) | SIK2 (QIK) | NE | NE | NE | >2.50E-05 |
| Tivozanib (AV-951) | SIK3 | NE | NE | NE | >2.50E-05 |
| Tivozanib (AV-951) | STK4 (MST1) | NE | NE | NE | >2.50E-05 |
| Tivozanib (AV-951) | SYK | NE | NE | NE | >2.50E-05 |
| Tivozanib (AV-951) | TLK2 | NE | NE | NE | >2.50E-05 |
| Tivozanib (AV-951) | TYK2 | NE | NE | NE | >2.50E-05 |
| Tivozanib (AV-951) | ULK1 | NE | NE | NE | >2.50E-05 |
| Tivozanib (AV-951) | ULK2 | NE | NE | NE | >2.50E-05 |
| Tivozanib (AV-951) | YSK1 (STK25) | NE | NE | NE | >2.50E-05 |
| Tivozanib (AV-951) | ZAP70 | NE | NE | NE | >2.50E-05 |
| Tivozanib (AV-951) | FGFR4 | NE | NE | NE | >2.50E-06 |
| Tivozanib (AV-951) | FLT3 F691L | NE | NE | NE | >2.50E-06 |
| Tivozanib (AV-951) | TAK1-TAB1 (MAP3K7-MAP3K7IP1) | NE | NE | NE | >2.50E-06 |

**Table S8.** Table of TR-FRET (KINETICfinder) binding kinetic parameters for tucatinib against a panel of kinases

| Compound ID | Target | $k_{on}$ ($M^{-1}$ $s^{-1}$) | $k_{off}$ ($s^{-1}$) | τ (min) | $K_d$ (M) |
|---|---|---|---|---|---|
| Tucatinib | HER2 (ERBB2) | 3.38E+05 | 3.42E-05 | 487.75 | 1.01E-10 |
| Tucatinib | HER4 (ERBB4) | 1.60E+06 | 1.18E-02 | 1.41 | 7.37E-09 |
| Tucatinib | EGFR (ERBB1) (non-activated) | 1.09E+06 | 1.10E-02 | 1.51 | 1.01E-08 |
| Tucatinib | EGFR (ERBB1) E746_A750del | 1.27E+06 | 1.39E-02 | 1.20 | 1.10E-08 |
| Tucatinib | EGFR (ERBB1) (activated) | 2.23E+05 | 3.27E-03 | 5.10 | 1.47E-08 |
| Tucatinib | EGFR (ERBB1) L858R | 2.79E+05 | 1.03E-02 | 1.62 | 3.68E-08 |
| Tucatinib | KIT D816V | 1.27E+05 | 2.00E-01 | 0.08 | 1.57E-06 |
| Tucatinib | MEK1 (MAP2K1) | 3.38E+04 | 6.45E-02 | 0.26 | 1.91E-06 |
| Tucatinib | MER (MERTK) | 1.07E+04 | 2.73E-02 | 0.61 | 2.55E-06 |
| Tucatinib | LRRK2 | >1.26E+05 | >5.00E-01 | <0.033 | 3.95E-06 |
| Tucatinib | CDK19/cyclin C | >1.19E+05 | >5.00E-01 | <0.033 | 4.21E-06 |
| Tucatinib | LCK | 8.39E+03 | 4.43E-02 | 0.38 | 5.28E-06 |
| Tucatinib | RET | 1.53E+04 | 8.32E-02 | 0.20 | 5.44E-06 |
| Tucatinib | BTK (activated) | 6.49E+03 | 3.95E-02 | 0.42 | 6.09E-06 |
| Tucatinib | BLK | 2.37E+03 | 1.50E-02 | 1.11 | 6.32E-06 |
| Tucatinib | RIPK2 | 8.50E+03 | 6.43E-02 | 0.26 | 7.56E-06 |
| Tucatinib | PLK4 | >5.53E+04 | >5.00E-01 | <0.033 | 9.04E-06 |
| Tucatinib | BTK (non-activated) | 5.80E+03 | 5.95E-02 | 0.28 | 1.03E-05 |
| Tucatinib | PDGFRα | >4.58E+04 | >5.00E-01 | <0.033 | 1.09E-05 |
| Tucatinib | AURKB (AURORA B)/INCENP | 1.43E+04 | 2.04E-01 | 0.08 | 1.42E-05 |
| Tucatinib | CRAF (RAF1) | >2.96E+04 | >5.00E-01 | <0.033 | 1.69E-05 |
| Tucatinib | ROS1 | 1.18E+04 | 2.83E-01 | 0.06 | 2.39E-05 |
| Tucatinib | LIMK1 | 8.58E+03 | 2.07E-01 | 0.08 | 2.41E-05 |
| Tucatinib | CLK4 | 1.56E+02 | 3.82E-03 | 4.36 | 2.45E-05 |
| Tucatinib | ABL2 (ARG) | NE | NE | NE | >2.50E-05 |
| Tucatinib | ACK (TNK2) | NE | NE | NE | >2.50E-05 |
| Tucatinib | AKT1 | NE | NE | NE | >2.50E-05 |
| Tucatinib | ALK5 (TGFBR1) | NE | NE | NE | >2.50E-05 |
| Tucatinib | AMPK (A1/B1/G1) | NE | NE | NE | >2.50E-05 |
| Tucatinib | AMPK (A2/B1/G1) | NE | NE | NE | >2.50E-05 |
| Tucatinib | BRAF | NE | NE | NE | >2.50E-05 |
| Tucatinib | CAMK2a | NE | NE | NE | >2.50E-05 |
| Tucatinib | CAMK2d | NE | NE | NE | >2.50E-05 |
| Tucatinib | CDK16/cyclin Y | NE | NE | NE | >2.50E-05 |
| Tucatinib | CDK7/cyclin H/MNAT1 | NE | NE | NE | >2.50E-05 |
| Tucatinib | CHEK1 (CHK1) | NE | NE | NE | >2.50E-05 |
| Tucatinib | CHEK2 (CHK2) | NE | NE | NE | >2.50E-05 |
| Tucatinib | CK1e (CSNK1E) | NE | NE | NE | >2.50E-05 |
| Tucatinib | CLK2 | NE | NE | NE | >2.50E-05 |
| Tucatinib | CSF1R (FMS) | NE | NE | NE | >2.50E-05 |
| Tucatinib | DYRK1B | NE | NE | NE | >2.50E-05 |
| Tucatinib | ERK1 (MAPK3) | NE | NE | NE | >2.50E-05 |
| Tucatinib | ERK2 (MAPK1) | NE | NE | NE | >2.50E-05 |
| Tucatinib | FAK1 (PTK2) | NE | NE | NE | >2.50E-05 |
| Tucatinib | FER | NE | NE | NE | >2.50E-05 |
| Tucatinib | FGFR4 | NE | NE | NE | >2.50E-05 |
| Tucatinib | FLT3 (activated) | NE | NE | NE | >2.50E-05 |
| Tucatinib | FLT3 (non-activated) | NE | NE | NE | >2.50E-05 |
| Tucatinib | FLT4 (VEGFR3) | NE | NE | NE | >2.50E-05 |
| Tucatinib | GCN2 (EIF2AK4) | NE | NE | NE | >2.50E-05 |
| Tucatinib | INSR | NE | NE | NE | >2.50E-05 |
| Tucatinib | IRAK1 | NE | NE | NE | >2.50E-05 |
| Tucatinib | IRAK3 | NE | NE | NE | >2.50E-05 |
| Tucatinib | IRAK4 | NE | NE | NE | >2.50E-05 |
| Tucatinib | ITK | NE | NE | NE | >2.50E-05 |
| Tucatinib | KHS1 (MAP4K5) | NE | NE | NE | >2.50E-05 |
| Tucatinib | KIT (activated) | NE | NE | NE | >2.50E-05 |
| Tucatinib | KIT (non-activated) | NE | NE | NE | >2.50E-05 |
| Tucatinib | LOK (STK10) | NE | NE | NE | >2.50E-05 |
| Tucatinib | LYNA | NE | NE | NE | >2.50E-05 |
| Tucatinib | MEK7 (MAP2K7) | NE | NE | NE | >2.50E-05 |
| Tucatinib | MET (CMET) | NE | NE | NE | >2.50E-05 |
| Tucatinib | MINK1 | NE | NE | NE | >2.50E-05 |
| Tucatinib | MTOR | NE | NE | NE | >2.50E-05 |
| Tucatinib | NIK (MAP3K14) | NE | NE | NE | >2.50E-05 |

| | | | | | |
|---|---|---|---|---|---|
| Tucatinib | p38a (MAPK14) | NE | NE | NE | >2.50E-05 |
| Tucatinib | PDGFRβ (activated) | NE | NE | NE | >2.50E-05 |
| Tucatinib | PDGFRβ (non-activated) | NE | NE | NE | >2.50E-05 |
| Tucatinib | PDK1 | NE | NE | NE | >2.50E-05 |
| Tucatinib | PLK1 | NE | NE | NE | >2.50E-05 |
| Tucatinib | SIK1 | NE | NE | NE | >2.50E-05 |
| Tucatinib | SIK2 (QIK) | NE | NE | NE | >2.50E-05 |
| Tucatinib | SIK3 | NE | NE | NE | >2.50E-05 |
| Tucatinib | SRC | NE | NE | NE | >2.50E-05 |
| Tucatinib | SYK | NE | NE | NE | >2.50E-05 |
| Tucatinib | TAK1/TAB1 (MAP3K7/MAP3K7IP1) | NE | NE | NE | >2.50E-05 |
| Tucatinib | TLK2 | NE | NE | NE | >2.50E-05 |
| Tucatinib | TNIK | NE | NE | NE | >2.50E-05 |
| Tucatinib | TRKA (NTRK1) | NE | NE | NE | >2.50E-05 |
| Tucatinib | TRKB (NTRK2) | NE | NE | NE | >2.50E-05 |
| Tucatinib | TRKC (NTRK3) | NE | NE | NE | >2.50E-05 |
| Tucatinib | TYK2 | NE | NE | NE | >2.50E-05 |
| Tucatinib | ULK1 | NE | NE | NE | >2.50E-05 |
| Tucatinib | ULK2 | NE | NE | NE | >2.50E-05 |
| Tucatinib | ZAP70 | NE | NE | NE | >2.50E-05 |
| Tucatinib | MNK1 (MKNK1) | NE | NE | NE | >25000.0 |
| Tucatinib | MST1R (RON) | NE | NE | NE | >25000.0 |

**Table S9.** Table of TR-FRET (KINETICfinder) binding kinetic parameters for zimlovisertib against a panel of kinases

| Compound ID | Target | $k_{on}$ ($M^{-1}$ $s^{-1}$) | $k_{off}$ ($s^{-1}$) | $\tau$ (min) | $K_d$ (M) |
|---|---|---|---|---|---|
| Zimlovisertib (PF-06650833) | IRAK4 | 3.75E+06 | 8.14E-05 | 204.63 | 2.17E-11 |
| Zimlovisertib (PF-06650833) | CLK4 | 1.62E+06 | 7.22E-03 | 2.31 | 4.45E-09 |
| Zimlovisertib (PF-06650833) | IRAK1 | 2.61E+06 | 1.21E-02 | 1.38 | 4.63E-09 |
| Zimlovisertib (PF-06650833) | PLK4 | 1.05E+07 | 8.19E-02 | 0.20 | 7.78E-09 |
| Zimlovisertib (PF-06650833) | LRRK2 | 1.54E+06 | 1.40E-02 | 1.19 | 9.11E-09 |
| Zimlovisertib (PF-06650833) | IRAK3 | 1.00E+06 | 1.57E-02 | 1.06 | 1.57E-08 |
| Zimlovisertib (PF-06650833) | RET | 2.61E+06 | 6.95E-02 | 0.24 | 2.66E-08 |
| Zimlovisertib (PF-06650833) | LIMK1 | 1.51E+06 | 5.21E-02 | 0.32 | 3.46E-08 |
| Zimlovisertib (PF-06650833) | CK1e (CSNK1E) | 3.42E+05 | 1.94E-02 | 0.86 | 5.66E-08 |
| Zimlovisertib (PF-06650833) | ULK1 | 6.35E+05 | 3.88E-02 | 0.43 | 6.11E-08 |
| Zimlovisertib (PF-06650833) | CDK19/cyclin C | 7.37E+05 | 8.01E-02 | 0.21 | 1.09E-07 |
| Zimlovisertib (PF-06650833) | MEK1 (MAP2K1) | 1.31E+05 | 1.53E-02 | 1.09 | 1.16E-07 |
| Zimlovisertib (PF-06650833) | MNK1 (MKNK1) | 2.17E+05 | 2.56E-02 | 0.65 | 1.18E-07 |
| Zimlovisertib (PF-06650833) | ALK5 (TGFBR1) | 1.10E+06 | 1.32E-01 | 0.13 | 1.20E-07 |
| Zimlovisertib (PF-06650833) | ITK | 2.28E+05 | 3.39E-02 | 0.49 | 1.48E-07 |
| Zimlovisertib (PF-06650833) | MEK7 (MAP2K7) | 3.04E+05 | 5.36E-02 | 0.31 | 1.76E-07 |
| Zimlovisertib (PF-06650833) | CLK2 | 1.08E+06 | 1.96E-01 | 0.08 | 1.83E-07 |
| Zimlovisertib (PF-06650833) | CHEK2 (CHK2) | 2.37E+05 | 5.30E-02 | 0.31 | 2.24E-07 |
| Zimlovisertib (PF-06650833) | ROS1 | 2.25E+05 | 5.29E-02 | 0.32 | 2.35E-07 |
| Zimlovisertib (PF-06650833) | CAMK2a | 1.15E+04 | 3.42E-03 | 4.88 | 2.97E-07 |
| Zimlovisertib (PF-06650833) | RIPK2 | 2.92E+05 | 9.11E-02 | 0.18 | 3.12E-07 |
| Zimlovisertib (PF-06650833) | DYRK1B | 9.93E+04 | 3.14E-02 | 0.53 | 3.16E-07 |
| Zimlovisertib (PF-06650833) | CDK7/cyclin H/MNAT1 | 4.40E+04 | 1.44E-02 | 1.16 | 3.27E-07 |
| Zimlovisertib (PF-06650833) | TLK2 | 4.22E+04 | 1.57E-02 | 1.06 | 3.72E-07 |
| Zimlovisertib (PF-06650833) | CAMK2d | 1.41E+04 | 7.60E-03 | 2.19 | 5.37E-07 |
| Zimlovisertib (PF-06650833) | TRKB (NTRK2) | 6.79E+05 | 3.72E-01 | 0.04 | 5.47E-07 |
| Zimlovisertib (PF-06650833) | CDK16/cyclin Y | 7.02E+04 | 3.93E-02 | 0.42 | 5.59E-07 |
| Zimlovisertib (PF-06650833) | TAK1/TAB1 (MAP3K7/MAP3K7IP1) | 1.67E+04 | 9.82E-03 | 1.70 | 5.88E-07 |
| Zimlovisertib (PF-06650833) | GCN2 (EIF2AK4) | 2.21E+04 | 1.45E-02 | 1.15 | 6.53E-07 |
| Zimlovisertib (PF-06650833) | AURKB (AURORA B)/INCENP | 4.43E+04 | 3.18E-02 | 0.52 | 7.18E-07 |
| Zimlovisertib (PF-06650833) | ULK2 | 3.91E+04 | 4.21E-02 | 0.40 | 1.08E-06 |
| Zimlovisertib (PF-06650833) | PDK1 | 2.93E+04 | 3.59E-02 | 0.46 | 1.23E-06 |
| Zimlovisertib (PF-06650833) | TYK2 | 7.86E+04 | 1.09E-01 | 0.15 | 1.39E-06 |
| Zimlovisertib (PF-06650833) | AMPK (A1/B1/G1) | 5.85E+04 | 8.28E-02 | 0.20 | 1.41E-06 |
| Zimlovisertib (PF-06650833) | KIT D816V | 2.16E+05 | 3.39E-01 | 0.05 | 1.57E-06 |
| Zimlovisertib (PF-06650833) | MST1R (RON) | 2.58E+04 | 4.52E-02 | 0.37 | 1.75E-06 |
| Zimlovisertib (PF-06650833) | FER | 2.40E+04 | 5.08E-02 | 0.33 | 2.11E-06 |
| Zimlovisertib (PF-06650833) | SIK1 | 6.89E+03 | 1.49E-02 | 1.12 | 2.16E-06 |
| Zimlovisertib (PF-06650833) | AMPK (A2/B1/G1) | 1.32E+05 | 2.95E-01 | 0.06 | 2.24E-06 |
| Zimlovisertib (PF-06650833) | FLT4 (VEGFR3) | >2.19E+05 | >5.00E-01 | <0.033 | 2.29E-06 |
| Zimlovisertib (PF-06650833) | PLK1 | 8.87E+03 | 2.13E-02 | 0.78 | 2.40E-06 |
| Zimlovisertib (PF-06650833) | KHS1 (MAP4K5) | 6.17E+03 | 1.54E-02 | 1.09 | 2.49E-06 |
| Zimlovisertib (PF-06650833) | HER2 (ERBB2) | 7.51E+03 | 2.21E-02 | 0.75 | 2.95E-06 |
| Zimlovisertib (PF-06650833) | EGFR (ERBB1) L858R | 2.95E+03 | 1.07E-02 | 1.56 | 3.62E-06 |
| Zimlovisertib (PF-06650833) | PDGFRα | 9.95E+04 | 3.83E-01 | 0.04 | 3.85E-06 |
| Zimlovisertib (PF-06650833) | TRKA (NTRK1) | 1.04E+04 | 4.76E-02 | 0.35 | 4.56E-06 |
| Zimlovisertib (PF-06650833) | MER (MERTK) | 4.04E+03 | 2.27E-02 | 0.73 | 5.63E-06 |
| Zimlovisertib (PF-06650833) | JAK1 (JH1 JH2) | >8.57E+04 | >5.00E-01 | <0.033 | 5.84E-06 |
| Zimlovisertib (PF-06650833) | EGFR (ERBB1) E746_A750del | 1.35E+04 | 7.90E-02 | 0.21 | 5.86E-06 |
| Zimlovisertib (PF-06650833) | FLT3 (activated) | >8.06E+04 | >5.00E-01 | <0.033 | 6.20E-06 |
| Zimlovisertib (PF-06650833) | TRKC (NTRK3) | >8.04E+04 | >5.00E-01 | <0.033 | 6.22E-06 |
| Zimlovisertib (PF-06650833) | AKT1 | 4.98E+03 | 3.14E-02 | 0.53 | 6.31E-06 |
| Zimlovisertib (PF-06650833) | BLK | 4.76E+03 | 3.16E-02 | 0.53 | 6.64E-06 |
| Zimlovisertib (PF-06650833) | LOK (STK10) | 4.95E+03 | 3.32E-02 | 0.50 | 6.71E-06 |
| Zimlovisertib (PF-06650833) | FAK1 (PTK2) | 3.45E+03 | 2.79E-02 | 0.60 | 8.10E-06 |
| Zimlovisertib (PF-06650833) | BRAF | 4.07E+03 | 3.31E-02 | 0.50 | 8.13E-06 |
| Zimlovisertib (PF-06650833) | LCK | 5.98E+03 | 4.95E-02 | 0.34 | 8.29E-06 |
| Zimlovisertib (PF-06650833) | SIK2 (QIK) | 3.34E+03 | 2.90E-02 | 0.58 | 8.66E-06 |
| Zimlovisertib (PF-06650833) | EGFR (ERBB1) (activated) | 4.02E+03 | 3.50E-02 | 0.48 | 8.70E-06 |
| Zimlovisertib (PF-06650833) | SRC | 1.40E+04 | 1.69E-01 | 0.10 | 1.21E-05 |
| Zimlovisertib (PF-06650833) | FGFR4 | 3.47E+03 | 4.38E-02 | 0.38 | 1.26E-05 |
| Zimlovisertib (PF-06650833) | SYK | 4.38E+03 | 5.59E-02 | 0.30 | 1.28E-05 |
| Zimlovisertib (PF-06650833) | LYNA | 2.40E+03 | 3.39E-02 | 0.49 | 1.41E-05 |
| Zimlovisertib (PF-06650833) | ACK (TNK2) | 1.80E+04 | 2.77E-01 | 0.06 | 1.54E-05 |
| Zimlovisertib (PF-06650833) | TNIK | 1.17E+03 | 2.16E-02 | 0.77 | 1.84E-05 |
| Zimlovisertib (PF-06650833) | ERK2 (MAPK1) | 2.52E+03 | 5.00E-02 | 0.33 | 1.98E-05 |

| | | | | | |
|---|---|---|---|---|---|
| Zimlovisertib (PF-06650833) | MET (CMET) | 1.33E+03 | 3.19E-02 | 0.52 | 2.40E-05 |
| Zimlovisertib (PF-06650833) | EGFR (ERBB1) (non-activated) | 1.52E+03 | 3.73E-02 | 0.45 | 2.46E-05 |
| Zimlovisertib (PF-06650833) | ABL2 (ARG) | NE | NE | NE | >2.50E-05 |
| Zimlovisertib (PF-06650833) | CHEK1 (CHK1) | NE | NE | NE | >2.50E-05 |
| Zimlovisertib (PF-06650833) | CRAF (RAF1) | NE | NE | NE | >2.50E-05 |
| Zimlovisertib (PF-06650833) | CSF1R (FMS) | NE | NE | NE | >2.50E-05 |
| Zimlovisertib (PF-06650833) | ERK1 (MAPK3) | NE | NE | NE | >2.50E-05 |
| Zimlovisertib (PF-06650833) | FLT3 (non-activated) | NE | NE | NE | >2.50E-05 |
| Zimlovisertib (PF-06650833) | HER4 (ERBB4) | NE | NE | NE | >2.50E-05 |
| Zimlovisertib (PF-06650833) | INSR | NE | NE | NE | >2.50E-05 |
| Zimlovisertib (PF-06650833) | KIT (activated) | NE | NE | NE | >2.50E-05 |
| Zimlovisertib (PF-06650833) | KIT (non-activated) | NE | NE | NE | >2.50E-05 |
| Zimlovisertib (PF-06650833) | MINK1 | NE | NE | NE | >2.50E-05 |
| Zimlovisertib (PF-06650833) | MTOR | NE | NE | NE | >2.50E-05 |
| Zimlovisertib (PF-06650833) | NIK (MAP3K14) | NE | NE | NE | >2.50E-05 |
| Zimlovisertib (PF-06650833) | p38a (MAPK14) | NE | NE | NE | >2.50E-05 |
| Zimlovisertib (PF-06650833) | PDGFRβ (activated) | NE | NE | NE | >2.50E-05 |
| Zimlovisertib (PF-06650833) | PDGFRβ (non-activated) | NE | NE | NE | >2.50E-05 |
| Zimlovisertib (PF-06650833) | SIK3 | NE | NE | NE | >2.50E-05 |
| Zimlovisertib (PF-06650833) | ZAP70 | NE | NE | NE | >2.50E-05 |

**Table S10.** Limiting $k_{on}$ and $k_{off}$ values and affinity thresholds for the different target classes and inhibitors analyzed in **Figures 1-4**, calculated by nonlinear regression of the $k_{on}$ data to *Equation 1*.

| Target(s) | Ligand(s) | Data Source | Limiting $k_{on}$ (x10$^6$ M$^{-1}$s$^{-1}$) [95% CI] | Limiting $k_{off}$ (s$^{-1}$) [95% CI] | $K_d$ threshold (nM) [95% CI] |
|---|---|---|---|---|---|
| Kinases | | | | | |
| Phospho-BTK | Various | This study (Fig 1A) | 1.4 [0.7-2.1] | 0.022 [0.011-0.032] | 16 [5-46] |
| Unphos-BTK | Various | This study (Fig 1A) | 1.0 [0.6-1.5] | 0.030 [0.014-0.045] | 30 [9-75] |
| Kinases | Various | This study (Fig 2A) | 0.77 [0.56-1.1] | 0.032 [0.026-0.038] | 42 [24-68] |
| Kinases (136) | Dasatinib | This study (Fig 2B) | 2.8 [1.6-4.8] | 0.029 [0.021-0.040] | 10 [4.4-24] |
| Kinases (134) | Danusertib | This study (Fig 2C) | 1.0 [0.68-1.5] | 0.040 [0.031-0.052] | 39 [21-76] |
| Kinases (123) | Foretinib | This study (Fig 2D) | 0.10 [0.06-0.20] | 0.061 [0.021-0.17] | 580 [110-3200] |
| Kinases (113) | Tivozanib | This study (Fig 2E) | 0.15 [0.06-0.36] | 0.077 [0.027-0.22] | 510 [75-3500] |
| Kinases (135) | Encorafenib | This study (Fig 2F) | 0.32 [0.08-1.3] | 0.010 [0.006-0.016] | 31 [5-200] |
| Kinases (87) | Tucatinib | This study (Fig 2G) | 0.65 [0.2-2.1] | 0.11 [0.058-0.21] | 170 [27-1100] |
| Kinases (86) | Zimlovisertib | This study (Fig 2H) | 4.3 [1.4-14] | 0.043 [0.033-0.056] | 9.9 [2.4-41] |
| Kinases | Various | KIND (Fig 3A) | 4.3 [3.5-5.1] | 0.042 [0.039-0.044] | 9.8 [7.6-13] |
| GPCRs | | | | | |
| D2 | Various | KIND (Fig 3C, S3) | 26 [18-38] | 0.027 [0.018-0.041] | 1.0 [0.5-2.3] |
| mGlu | Various | KIND (Fig 3D, S3) | ≥1 | 0.0014 [0.0012-0.0016] | ≤1.4 |
| A1A-R | Various | KIND (Fig S3) | 0.42 [0.18-0.96] | ≥0.5 | ≥1200 |
| A2A-R | Various | KIND (Fig S3) | 2.9 [1.6-5.5] | 0.024 [0.013-0.046] | 8.3 [2.4-29] |
| Beta-2 | Various | KIND (Fig S3) | 17 [8.7-34] | 0.052 [0.025-0.11] | 3.1 [0.74-13] |
| CRF1 | Various | KIND (Fig S3) | 0.26 [0.13-0.53] | 0.0028 [0.0009-0.0086] | 11 [1.7-66] |
| H1 | Various | KIND (Fig S3) | 0.95 [0.12-7.4] | 0.0031 [0.0006-0.016] | 3.3 [0.8-133] |
| M3 | Various | KIND (Fig S3) | 16 [9.8-25] | ≥0.004 | ≥0.25 |
| Other | | | | | |
| HSP90 | Various | KIND (Fig 3E) | 0.19 [0.15-0.24] | 0.20 [0.09-0.45] | 1100 [375-3000] |
| hERG | Various | KIND (Fig 3F) | ≥100 | 0.0024 [0.0016-0.0035] | ≤0.02 |
| InhA | Various | *Spagnuolo et al.*[7] (Fig 4C) | ≥0.05 | 0.00023 [0.00016-0.00034] | n/a |

**Table S11.** SPR and TR-FRET (KINETICfinder) binding kinetic data.
**A. BTK:** SPR values determined in this work and TR-FRET (KINETICfinder) values from Bravo et al.

| | $k_{on}$ ($M^{-1}$ $s^{-1}$) | $k_{off}$ ($s^{-1}$) | $K_d$ (M) | τ |
|---|---|---|---|---|
| P301390 | (4.9±0.9)x10$^{6}$<br>(1.75±0.31)x10$^{6}$ | (7.35±1.20) x10$^{-2}$<br>(4.90±1.43)x10$^{-2}$ | (1.51±0.03) x10$^{-8}$<br>(2.75±0.48)x10$^{-8}$ | 13.6 s<br>22.8 s |
| BTK-IN-1 | (6.87±0.73)x10$^{5}$<br>(1.58±0.23)x10$^{5}$ | (6.74±2.73) x10$^{-2}$<br>(3.39±0.52)x10$^{-2}$ | (1.01±0.51) x10$^{-7}$<br>(2.14±0.04)x10$^{-7}$ | 32 s<br>29.4 s |
| GDC0834 | (2.16±1.53)x10$^{6}$<br>(1.23±0.05)x10$^{6}$ | (1.90±0.19) x10$^{-3}$<br>(2.34±0.12)x10$^{-3}$ | (8.14±0.03) x10$^{-10}$<br>(1.91±0.04)x10$^{-9}$ | 8.77 min<br>7.12 min |
| GDC0853 | (1.91±0.34)x10$^{6}$<br>(8.17±0.84)x10$^{5}$ | (2.08±0.23) x10$^{-4}$<br>(6.34±0.30)x10$^{-5}$ | (1.10±0.08) x10$^{-10}$<br>(7.84±0.88)x10$^{-11}$ | 80 min<br>263 min |

Errors are expressed as SD (n=2). Values from *Bravo et al.*[2]

**B: Data for other kinases:** TR-FRET (KINETICfinder) binding kinetic data determined here and SPR data from the literature.[3,8-16]

| | **TR-FRET (KINETICfinder)** | | | **SPR**[1] | | |
|---|---|---|---|---|---|---|
| **Enzyme-ligand** | Log $K_d$ | Log $k_{on}$ | Log $k_{off}$ | Log $K_d$ | Log $k_{on}$ | Log $k_{off}$ |
| Imatinib-SRC | -4.477 | 2.726 | -1.751 | -5.006 | 3.386 | -1.620 |
| Ponatinib-SRC | -8.316 | 4.133 | -4.183 | -8.219 | 4.415 | -3.804 |
| Pazopanib-KIT | -8.805 | 4.835 | -3.971 | -7.266 | 3.567 | -3.699 |
| Dasatinib-FLT3 | -5.022 | 2.790 | -2.232 | -5.607 | 3.380 | -2.227 |
| Dasatinib-CSF1R | -9.660 | 6.474 | -3.186 | -8.413 | 5.455 | -2.959 |
| Sorafenib-p38a | -6.377 | 4.718 | -1.659 | -6.220 | 4.280 | -1.940 |
| Axitinib-ABL1 | -8.671 | 7.335 | -1.336 | -8.398 | 6.381 | -2.017 |
| Dasatinib-ABL1 | -10.732 | 7.042 | -3.690 | -9.886 | 5.869 | -4.017 |
| Ponatinib -ABL1 | -9.486 | 5.355 | -4.130 | -9.444 | 5.036 | -4.407 |
| Pictilisib-PI3Kg | -7.459 | 5.807 | -1.652 | -7.614 | 6.346 | -1.268 |
| Idelalisib-PI3Kg | -7.894 | 6.062 | -1.832 | -7.780 | 6.013 | -1.767 |
| Pictilisib-PI3Kd | -8.576 | 6.337 | -2.239 | -8.635 | 6.604 | -2.031 |
| Duvelisib-PI3Kd | -10.884 | 7.018 | -3.866 | -10.538 | 6.725 | -3.917 |
| Idelalisib-PI3Kd | -9.655 | 6.835 | -2.820 | -9.137 | 6.773 | -2.365 |
| Pictilisib-PI3Ka | -8.177 | 6.118 | -2.059 | -8.959 | 6.612 | -2.348 |
| Idelalisib-PI3Ka | -6.754 | 5.738 | -1.016 | -6.914 | 6.053 | -0.863 |
| SNS-032-CDK2/A | -8.099 | | -1.477 | | | -1.954 |
| Dinaciclib-CDK2/A | -8.925 | | -2.924 | | | -2.556 |
| AT-7519-CDK2/A | -7.971 | | -2.079 | | | -1.477 |
| SNS-032-CDK9/T | -8.877 | | -2.079 | | | -2.255 |
| Dinaciclib-CDK9/T | -9.499 | | -3.635 | | | -3.681 |
| AT-7519-CDK9/T | -9.286 | | -3.193 | | | -2.556 |
| AT-7519-CDK2/E | -7.353 | | -1.853 | | | -1.477 |
| Staurosporine-ASK1 | -8.270 | 6.219 | -2.051 | -8.155 | 5.648 | -2.506 |
| Bosutinib-ABL1 | -8.801 | 6.497 | -2.304 | -8.664 | 5.858 | -2.804 |
| Dasatinib-BTK | -9.617 | 6.577 | -3.041 | -9.071 | 5.690 | -3.380 |
| Ponatinib-BTK | -6.988 | 5.702 | -1.286 | -6.106 | 4.332 | -1.775 |
| Staurosporine-BTK | -7.680 | 6.389 | -1.291 | -7.676 | 5.801 | -2.000 |
| Ponatinib-LCK | -9.126 | 5.650 | -3.475 | -8.754 | 5.104 | -3.650 |
| Bosutinib-TNIK | -7.971 | 5.514 | -2.457 | -7.845 | 6.097 | -1.750 |
| Gefitinib-EGFR | -9.949 | 7.273 | -2.676 | -8.854 | 6.737 | -2.117 |
| Erlotinib-EGFR | -10.078 | 7.559 | -2.519 | -9.156 | 6.759 | -2.480 |
| Lapatinib-EGFR | -8.363 | 4.881 | -3.482 | -8.375 | 5.045 | -3.328 |
| Dasatinib-EGFR | -8.083 | 6.363 | -1.720 | -7.759 | 5.893 | -2.000 |
| Staurosporine-EGFR | -7.559 | 6.125 | -1.434 | -7.291 | 5.693 | -1.523 |
| Crizotinib-ALK | -9.088 | 6.737 | -2.351 | -8.518 | 7.068 | -1.450 |
| Ponatinib-FGFR1 | -7.553 | 4.352 | -3.201 | -8.125 | 4.380 | -3.745 |

| | | | | | | |
|---|---|---|---|---|---|---|
| Sunitinib-IRAK4 | -7.252 | 5.644 | -1.608 | -6.834 | 5.856 | -0.979 |
| SORAFENIB-CDK8/C | -7.112 | 3.290 | -3.822 | -7.002 | 3.509 | -3.492 |
| LINIFANIB-CDK8/C | -7.858 | 4.057 | -3.801 | -8.165 | 3.918 | -4.248 |
| PONATINIB-CDK8/C | -8.425 | 4.113 | -4.311 | -8.169 | 3.682 | -4.487 |
| Danusertib-AURKA | -8.465 | 5.535 | -2.930 | -8.772 | 5.709 | -3.061 |
| Tozasertib-AURKA | -8.486 | 6.157 | -2.329 | -8.523 | 5.771 | -2.752 |
| AMG900-AURKB | -9.159 | 4.682 | -4.477 | -9.406 | 4.769 | -4.636 |
| Danusertib-AURKB | -9.022 | 4.691 | -4.331 | -8.842 | 4.637 | -4.203 |
| GSK1070916-AURKB | -9.527 | 4.705 | -4.823 | -8.349 | 4.146 | -4.202 |
| MK5108-AURKB | -8.272 | 6.135 | -2.138 | -8.021 | 5.972 | -2.049 |
| MLN8054-AURKB | -8.574 | 5.405 | -3.169 | -8.580 | 4.815 | -3.764 |
| Tozasertib-AURKB | -8.698 | 4.998 | -3.701 | -8.558 | 4.498 | -4.059 |
| BI 2536-PLK1 | -9.217 | 5.670 | -3.547 | -7.971 | 4.670 | -3.302 |
| Crizotinib-MET | -9.577 | 6.489 | -3.087 | -7.625 | 4.870 | -2.754 |
| Cabozantinib-AURKA | -5.269 | 4.327 | -0.941 | -5.498 | 4.629 | -0.869 |
| Crizotinib-AURKA | -6.792 | 6.141 | -0.651 | -7.081 | 5.888 | -1.193 |
| Danusertib-AURKA | -8.399 | 5.458 | -2.941 | -8.006 | 5.325 | -2.681 |
| Barasertib -AURKA | -6.434 | 5.900 | -0.534 | -6.833 | 6.249 | -0.584 |
| GSK1070916-AURKA | -7.287 | 5.260 | -2.027 | -7.821 | 5.530 | -2.291 |
| Midostaurin-AURKA | -7.195 | 5.072 | -2.122 | -7.718 | 5.377 | -2.341 |
| Pazopanib-AURKA | -6.228 | 5.851 | -0.377 | -6.012 | 5.245 | -0.767 |
| Tozasertib-AURKA | -8.486 | 6.157 | -2.329 | -8.579 | 6.578 | -2.001 |
| GSK1070916-AURKA | -7.287 | 5.260 | -2.027 | -7.880 | 5.260 | -2.620 |
| Dasatinib-BMX | -10.100 | 6.551 | -3.549 | -9.480 | 5.820 | -3.660 |
| Dabrafenib-BRAF | -9.495 | 6.156 | -3.339 | -9.210 | 5.670 | -3.540 |
| Crizotinib-FAK | -7.849 | 6.957 | -0.892 | -7.870 | 6.930 | -0.940 |
| Staurosporine-MAP3K5 | -8.152 | 5.503 | -2.648 | -8.160 | 5.650 | -2.510 |
| Duvelisib-PI3Ka/P85a H1047R | -7.411 | 6.262 | -1.150 | -7.523 | 6.230 | -1.292 |
| BIRB796-p38a | -8.959 | 3.982 | -4.959 | -10.000 | 4.924 | -5.081 |

[1]References for SPR data[3,8-16]

(15) Willemsen-Seegers, N.; Uitdehaag, J. C. M.; Prinsen, M. B. W.; de Vetter, J. R. F.; de Man, J.; Sawa, M.; Kawase, Y.; Buijsman, R. C.; Zaman, G. J. R. Compound Selectivity and

Target Residence Time of Kinase Inhibitors Studied with Surface Plasmon Resonance *J Mol Biol*, **2017**, *429*, 574-586. DOI: 10.1016/j.jmb.2016.12.019

(16) Buckbinder, L.; St Jean, D. J., Jr.; Tieu, T.; Ladd, B.; Hilbert, B.; Wang, W.; Alltucker, J. T.; Manimala, S.; Kryukov, G. V.; Brooijmans, N.; Dowdell, G.; Jonsson, P.; Huff, M.; Guzman-Perez, A.; Jackson, E. L.; Goncalves, M. D.; Stuart, D. D. STX-478, a Mutant-Selective, Allosteric PI3Kalpha Inhibitor Spares Metabolic Dysfunction and Improves Therapeutic Response in PI3Kalpha-Mutant Xenografts *Cancer Discov*, **2023**, *13*, 2432-2447. DOI: 10.1158/2159-8290.CD-23-0396
